# Tractography dissection variability: what happens when 42 groups dissect 14 white matter bundles on the same dataset?

**DOI:** 10.1101/2020.10.07.321083

**Authors:** Kurt G. Schilling, François Rheault, Laurent Petit, Colin B. Hansen, Vishwesh Nath, Fang-Cheng Yeh, Gabriel Girard, Muhamed Barakovic, Jonathan Rafael-Patino, Thomas Yu, Elda Fischi-Gomez, Marco Pizzolato, Mario Ocampo-Pineda, Simona Schiavi, Erick J. Canales-Rodríguez, Alessandro Daducci, Cristina Granziera, Giorgio Innocenti, Jean-Philippe Thiran, Laura Mancini, Stephen Wastling, Sirio Cocozza, Maria Petracca, Giuseppe Pontillo, Matteo Mancini, Sjoerd B. Vos, Vejay N. Vakharia, John S. Duncan, Helena Melero, Lidia Manzanedo, Emilio Sanz-Morales, Ángel Peña-Melián, Fernando Calamante, Arnaud Attyé, Ryan P. Cabeen, Laura Korobova, Arthur W. Toga, Anupa Ambili Vijayakumari, Drew Parker, Ragini Verma, Ahmed Radwan, Stefan Sunaert, Louise Emsell, Alberto De Luca, Alexander Leemans, Claude J. Bajada, Hamied Haroon, Hojjatollah Azadbakht, Maxime Chamberland, Sila Genc, Chantal M. W. Tax, Ping-Hong Yeh, Rujirutana Srikanchana, Colin D. Mcknight, Joseph Yuan-Mou Yang, Jian Chen, Claire E. Kelly, Chun-Hung Yeh, Jerome Cochereau, Jerome J. Maller, Thomas Welton, Fabien Almairac, Kiran K Seunarine, Chris A. Clark, Fan Zhang, Nikos Makris, Alexandra Golby, Yogesh Rathi, Lauren J. O’Donnell, Yihao Xia, Dogu Baran Aydogan, Yonggang Shi, Francisco Guerreiro Fernandes, Mathijs Raemaekers, Shaun Warrington, Stijn Michielse, Alonso Ramírez-Manzanares, Luis Concha, Ramón Aranda, Mariano Rivera Meraz, Garikoitz Lerma-Usabiaga, Lucas Roitman, Lucius S. Fekonja, Navona Calarco, Michael Joseph, Hajer Nakua, Aristotle N. Voineskos, Philippe Karan, Gabrielle Grenier, Jon Haitz Legarreta, Nagesh Adluru, Veena A. Nair, Vivek Prabhakaran, Andrew L. Alexander, Koji Kamagata, Yuya Saito, Wataru Uchida, Christina Andica, Masahiro Abe, Roza G. Bayrak, Claudia A.M. Gandini Wheeler-Kingshott, Egidio D’Angelo, Fulvia Palesi, Giovanni Savini, Nicolò Rolandi, Pamela Guevara, Josselin Houenou, Narciso López-López, Jean-François Mangin, Cyril Poupon, Claudio Román, Andrea Vázquez, Chiara Maffei, Mavilde Arantes, José Paulo Andrade, Susana Maria Silva, Vince D. Calhoun, Eduardo Caverzasi, Simone Sacco, Michael Lauricella, Franco Pestilli, Daniel Bullock, Yang Zhan, Edith Brignoni-Perez, Catherine Lebel, Jess E Reynolds, Igor Nestrasil, René Labounek, Christophe Lenglet, Amy Paulson, Stefania Aulicka, Sarah R. Heilbronner, Katja Heuer, Bramsh Qamar Chandio, Javier Guaje, Wei Tang, Eleftherios Garyfallidis, Rajikha Raja, Adam W. Anderson, Bennett A. Landman, Maxime Descoteaux

## Abstract

White matter bundle segmentation using diffusion MRI fiber tractography has become the method of choice to identify white matter fiber pathways *in vivo* in human brains. However, like other analyses of complex data, there is considerable variability in segmentation protocols and techniques. This can result in different reconstructions of the same intended white matter pathways, which directly affects tractography results, quantification, and interpretation. In this study, we aim to evaluate and quantify the variability that arises from different protocols for bundle segmentation. Through an open call to users of fiber tractography, including anatomists, clinicians, and algorithm developers, 42 independent teams were given processed sets of human whole-brain streamlines and asked to segment 14 white matter fascicles on six subjects. In total, we received 57 different bundle segmentation protocols, which enabled detailed volume-based and streamline-based analyses of agreement and disagreement among protocols for each fiber pathway. Results show that even when given the exact same sets of underlying streamlines, the variability across protocols for bundle segmentation is greater than all other sources of variability in the virtual dissection process, including variability within protocols and variability across subjects. In order to foster the use of tractography bundle dissection in routine clinical settings, and as a fundamental analytical tool, future endeavors must aim to resolve and reduce this heterogeneity. Although external validation is needed to verify the anatomical accuracy of bundle dissections, reducing heterogeneity is a step towards reproducible research and may be achieved through the use of standard nomenclature and definitions of white matter bundles and well-chosen constraints and decisions in the dissection process.

## Introduction

Diffusion MRI fiber tractography [1, 2] offers unprecedented insight into the structural connections of the human brain. In a process that parallels post-mortem microdissection, tractography – in combination with a set of rules, constraints, and procedures to dissect and segment major white matter fascicles of the brain – allows noninvasive visualization and quantification of the shape, location, connectivity, and biophysical properties of white matter bundles. This process of *in vivo* “virtual dissection” [3, 4], also called *bundle segmentation*, has led to new insight into how structural connectivity underlies brain function, cognition, and development, in addition to dysfunction in neurological diseases, mental health disorders, and aging [5]. Additionally, bundle segmentation is used routinely to provide critical clinical information in both pre-operative and intra-operative mapping of brain tumor resections [6, 7].

Despite widespread use in clinical and research domains, there are a large number of variations in workflows for bundle segmentation that have been adopted by the neuroimaging community. Normally, workflows either generate bundles of streamlines, i.e., digital representations of fiber trajectories, or dissect subsets of streamlines from an ensemble of streamlines throughout the whole brain. These protocols typically differ in the rules and constraints used to isolate a given pathway, ranging from manual delineation of inclusion and exclusion regions of interest, to fully automated segmentations based on shape, location, or connectivity. Contributing to this variability, agreements on the anatomical definitions of pathways in the human brain are far from settled [8–11], in part hindered by the lack of a consistent framework for defining tracts. Descriptive tract definitions have traditionally focused on the shape and area of convergence of axons deep in the white matter, but may also focus on the specific regions to which these fibers connect [9, 11–15]. Consequently, and coming full circle, differences and disagreements in anatomical definitions and their interpretation may lead to further variations in protocols used in the virtual dissection process.

For these reasons, the process of bundle segmentation has been described as existing somewhere between science and art [16]. Variation in protocols can result in different segmentations which can lead to different scientific conclusions or clinical decisions [17]. This inter-protocol variability adds “noise” to the literature when it comes to the process of bundle segmentation [18, 19], a variability that prevents a direct comparison of the outcomes of different studies, and hinders the translation of these techniques from the research laboratory to the clinic. Yet, an estimate of the variability that exists across different protocols remains unclear. In order to ultimately harmonize the anatomical definition of tracts and standardize the bundle segmentation process, we propose a first step is to quantify this variability, and understand the similarities and differences in bundle segmentation results across protocols.

Towards this end, the aims of this study are twofold: (1) to understand how much variability exists across different protocols for bundle segmentation, and (2) to quantify which fascicles exhibit the most agreement/disagreement across protocols. To do this we take a “many analysts, one dataset” approach previously used to study workflows for diffusion analysis [20], hippocampus segmentation [21], fMRI analysis [19, 22], and psychology research [23]. Through an open call to the community, we invited collaborations from expert scientists and clinicians who use tractography for bundle segmentation, provided them all with the same sets of tractography streamlines, and gave them the task of segmenting 14 white matter pathways from each dataset. This enabled streamline-based and volume-based quantification of inter-protocol agreement and disagreement for each fiber pathway and the results highlight the problem of variation of definitions and protocols for bundle segmentation.

## Results

### Submissions

We surveyed the protocols for bundle segmentation of 14 white matter bundles: Superior Longitudinal Fasciculus (SLF), Arcuate Fasciculus (AF), Optic Radiation (OR), Corticospinal Tract (CST), Cingulum (CG), Uncinate Fasciculus (UF), Corpus Callosum (CC), Middle Longitudinal Fasciculus (MdLF), Inferior Fronto-Occipital Fasciculus (IFOF), Inferior Longitudinal Fasciculus (ILF), Fornix (FX), Anterior Commissure (AC), Posterior Commissure (PC), and Parieto-Occipital Pontine Tract (POPT).

To isolate the effects of bundle segmentation from all other sources of variation, we directly provided six sets of whole-brain streamlines (both deterministic and probabilistic) to all collaborators, derived from 3 subjects with scan-rescan data acquired from the Human Connectome Project test-retest database [24]. Collaborators were given the choice of utilizing streamlines generated from one of two commonly used tractography methods, a deterministic or a probabilistic algorithm, which are known to generate different representations of white matter bundles and have different uses and applications as described in the literature [25, 26].

In total, this collaborative effort involved 144 collaborators from 42 teams (**Figure 2**, **top**). 57 unique sets of protocols were submitted, of which 28 submissions used the deterministic streamlines and 29 used probabilistic. A total of 3138 bundle tractograms were submitted. Because collaborators did not have to submit all bundles, pathways showed varying representation across submissions (**Figure 2**, **bottom**), ranging from as low as 16 protocols for the PC, up to 50 protocols for the CST.

**Figure 1.**
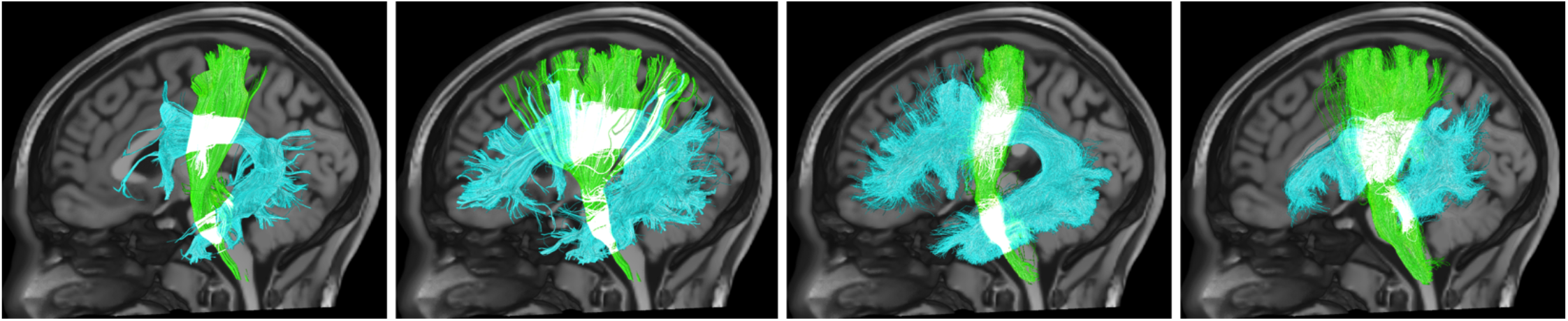
Variation in white matter bundle segmentation. Four example segmentations of the corticospinal tract (green) and arcuate fasciculus (cyan) show variability in the size, shape, densities, and connections of these reconstructed white matter pathways.

**Figure 2.**
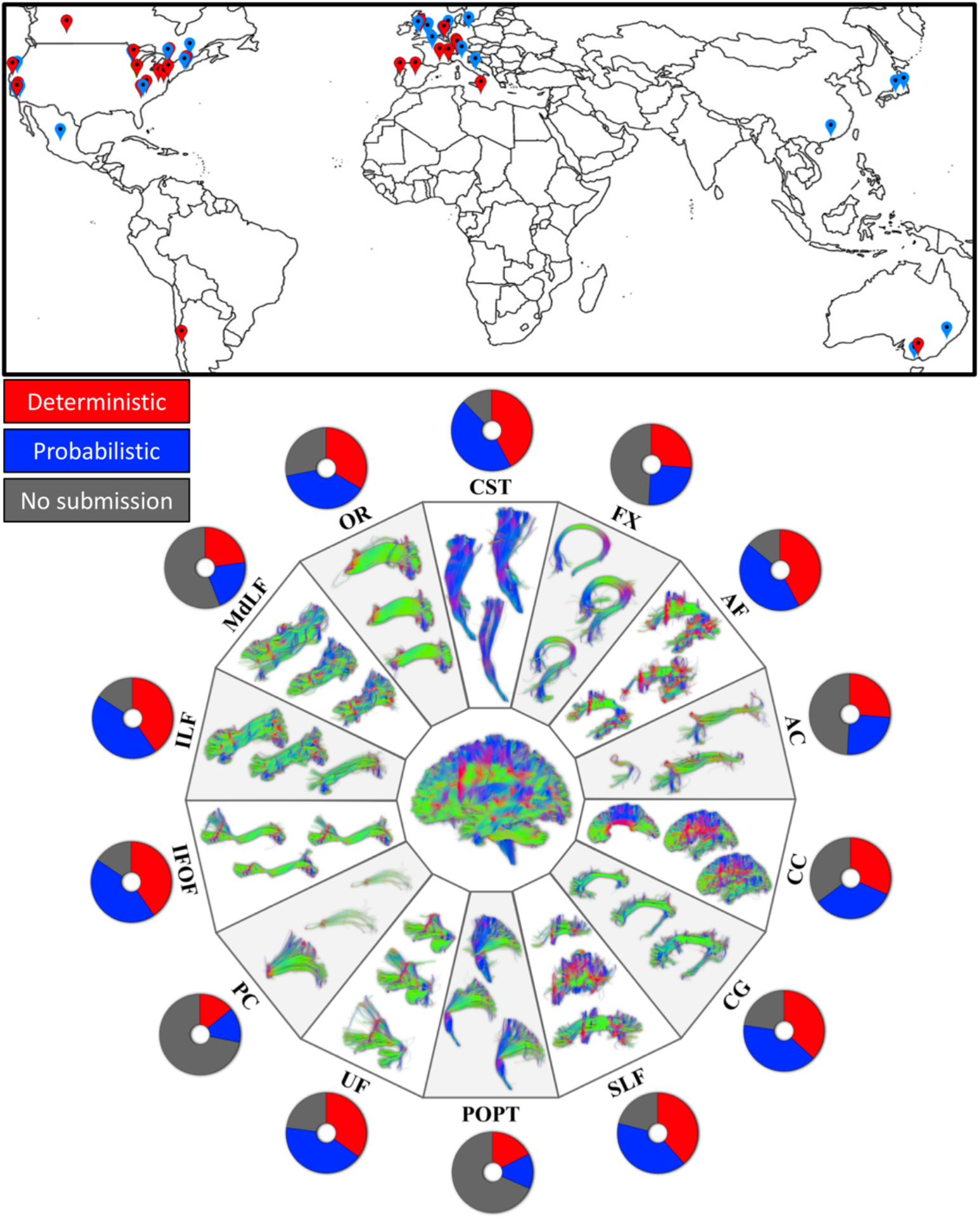
Summary of teams and submissions. Location of the teams’ affiliated lab (top). In total, 42 teams submitted 57 unique sets of bundle dissections, 28 utilized the provided deterministic streamlines, and 29 utilized probabilistic. Map icons are colored based on the set of streamlines utilized, with the same color-scheme as bar plots. Example submissions are shown for 14 pathways (bottom) along with a pie chart indicating the number of submissions for each bundle. Acronyms: see text.

### Qualitative Results

Example visualizations of randomly selected segmentations from a single subject are shown for exemplar projection, association, and commissural pathways (CST, AF, CC) in **Figure 3**. These are visualized as both streamlines directly, and also as 3D streamline density maps. The primary result from this figure is that there are many ways to segment these structures that result in qualitatively different representations of the same white matter pathways. These examples demonstrate visibly apparent variations in the size, shape, and connectivity patterns of streamlines. In contrast, different protocols result in similar patterns of high streamline density in the deep white matter and midbrain, with similar overall shape and central location. Similar visualizations, for all submitted pathways, both probabilistic and deterministic, are provided in supplementary documentation. These observations apply to all dissected pathways, however the commissural AC and PC contained very few streamlines, with little-to-no agreement across protocols.

**Figure 3.**
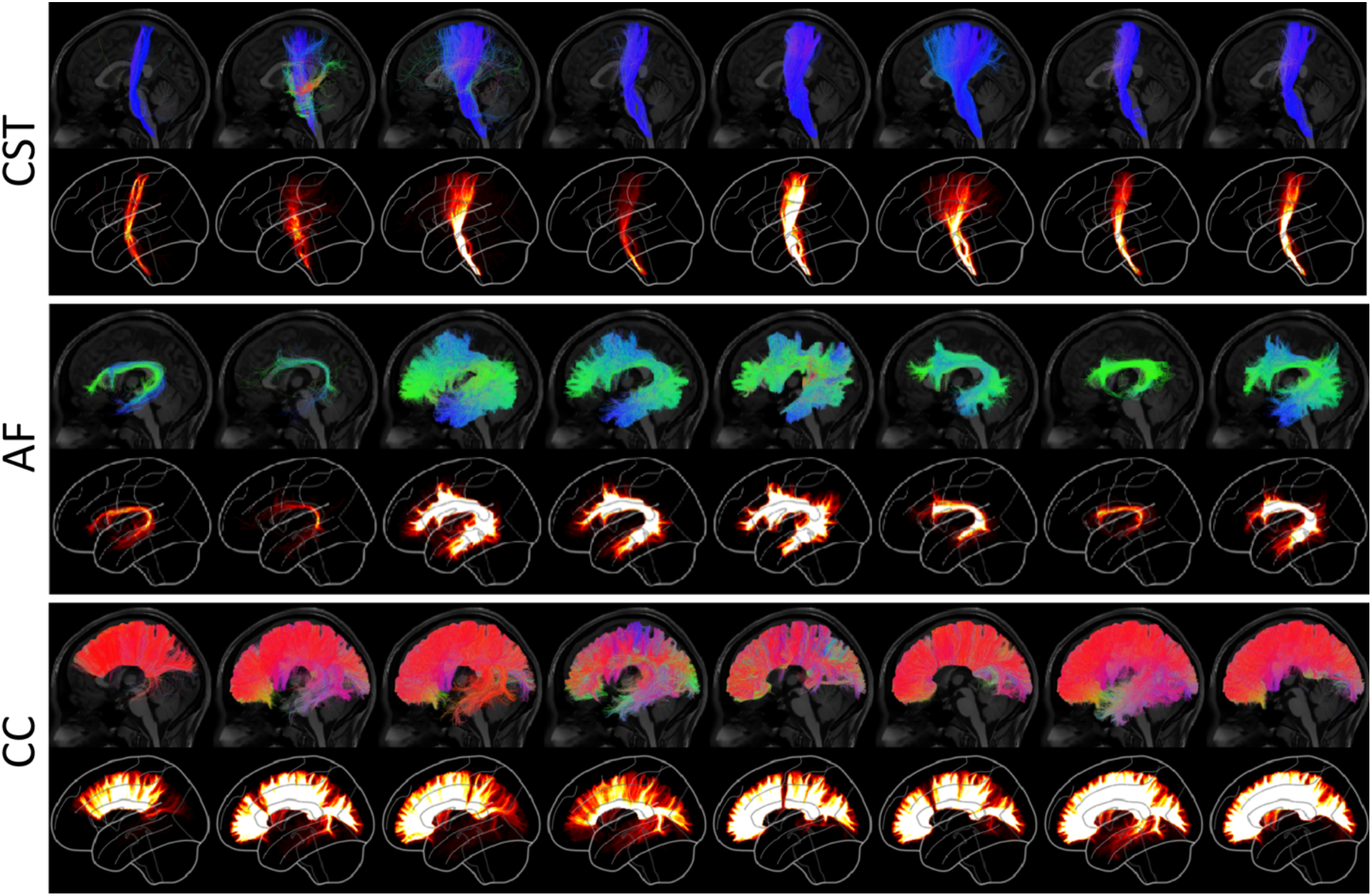
Variation in protocols for bundle segmentation of example pathways (CST, AF, and CC) on the same subject from the same set of whole-brain streamlines. Eight randomly selected bundle segmentation approaches for each pathway are shown as segmented streamlines and rendered as 3D streamline density maps. Variations in size, shape, density, and connectivity are qualitatively apparent. Probabilistic streamlines are shown, see supplementary material for Deterministic submissions. Random selections generated independently for each pathway. Streamlines are colored by orientation and all density maps are windowed to the same range.

### Pathway-Specific results

To understand the variability that exists across protocols for a given pathway, we visualize volume-based and streamline-based overlaps among the protocols and show boxplots of agreement measures that quantify inter-protocol, intra-protocol, and inter-subject variation. The volume overlap is displayed as the volume of voxels in which a given percent of protocols agree that the voxel was occupied by a given pathway, where a streamline overlap is displayed as the individual streamlines in which a given percent of protocols agree that streamline is representative of a given pathway. For quantitative analysis, we use several measures to describe similarity and dissimilarity of streamlines, streamline density, and pathway volume (**Figure 4**). This includes (1) v*olume Dice overlap* which describes the overall volume similarity, (2) *density correlation* which describes insight into similarity of streamline density, (3) *bundle adjacency* which describes the average distance of disagreement between two bundles, and (4) *streamline Dice* which describes the overlap of streamlines common between protocols (which can only be calculated because bundles come from the same original set of streamlines). We calculate geometric measures of pathways including number of streamlines, mean length, and volume, as well as microstructural measures of the average fractional anisotropy (FA) of the entire pathway volume and the FA weighted by streamline density (wFA).

**Figure 4.**
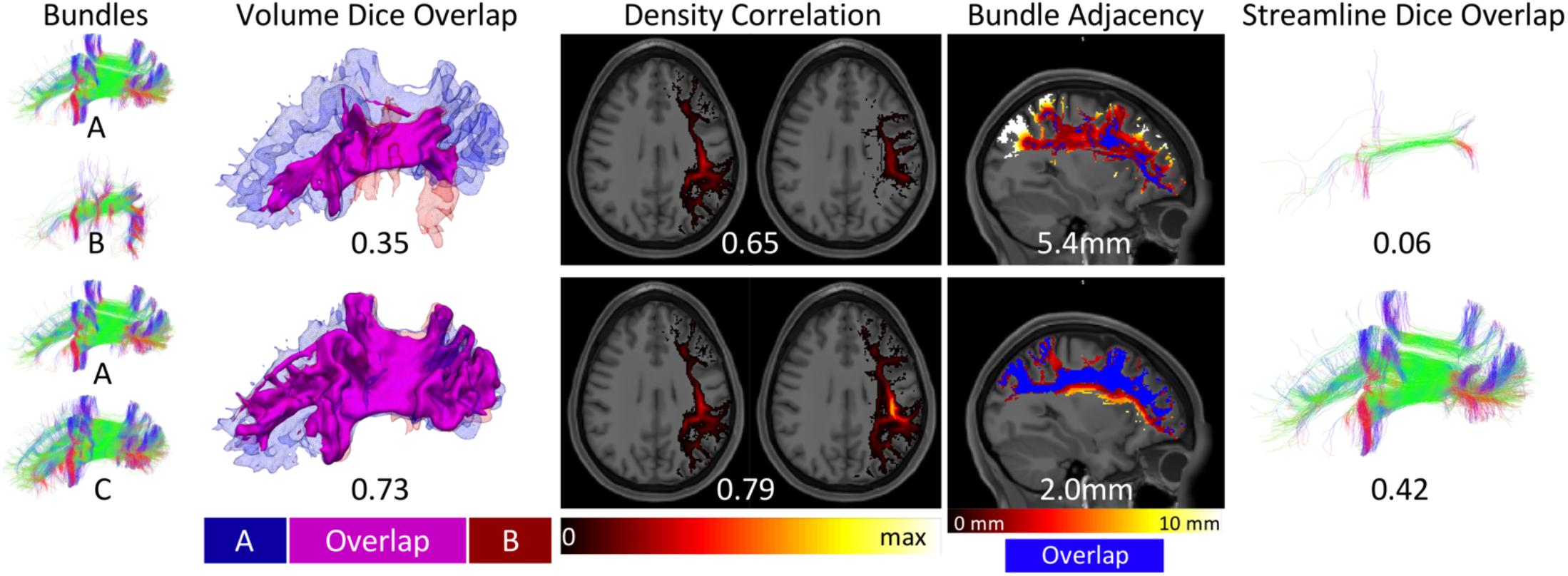
Similarity and dissimilarity metrics to assess reproducibility. Example SLF datasets are used to illustrate a range of similarity values between bundles A and B (top) and between bundles A and C (bottom). Dice overlap is a volume-based measure calculated as twice the intersection of two bundles (magenta) divided by the union (red and blue). Density correlation is calculated as the correlation coefficient between the voxel-wise streamline densities (shown as a hot-cold colormap ranging from 0 to maximum streamline density) of the two bundles being compared. Bundle adjacency is calculated by taking the average distance of disagreement (not including overlapping voxels in blue) between bundles (distances shown as hot-cold colormap). Finally, streamline Dice is taken as the intersection of common streamlines divided by the union of all streamlines in a bundle and requires input bundles to be segmented from the same set of underlying streamlines (intersection shown in figure).

For simplicity, we show results of the CST, AF, and CC. Analysis was conducted on all tracts, and results are provided in supplementary documentation.

#### Corticospinal Tract (CST)

**Figure 5** shows the results for the CST, and **Appendix A** summarizes the descriptive definitions and decisions made in the bundle segmentation workflow. Looking at the volume of agreement on a single subject, nearly all methods agree on the convergence of axons through the internal capsule and midbrain, with some disagreements on cortical terminations, and only a minority of protocols suggesting lateral projections of this tract. Streamline-based agreements show similar trends. The most striking result is that there were not any streamlines which were common to at least 75% of either the deterministic or probabilistic protocols.

**Figure 5.**
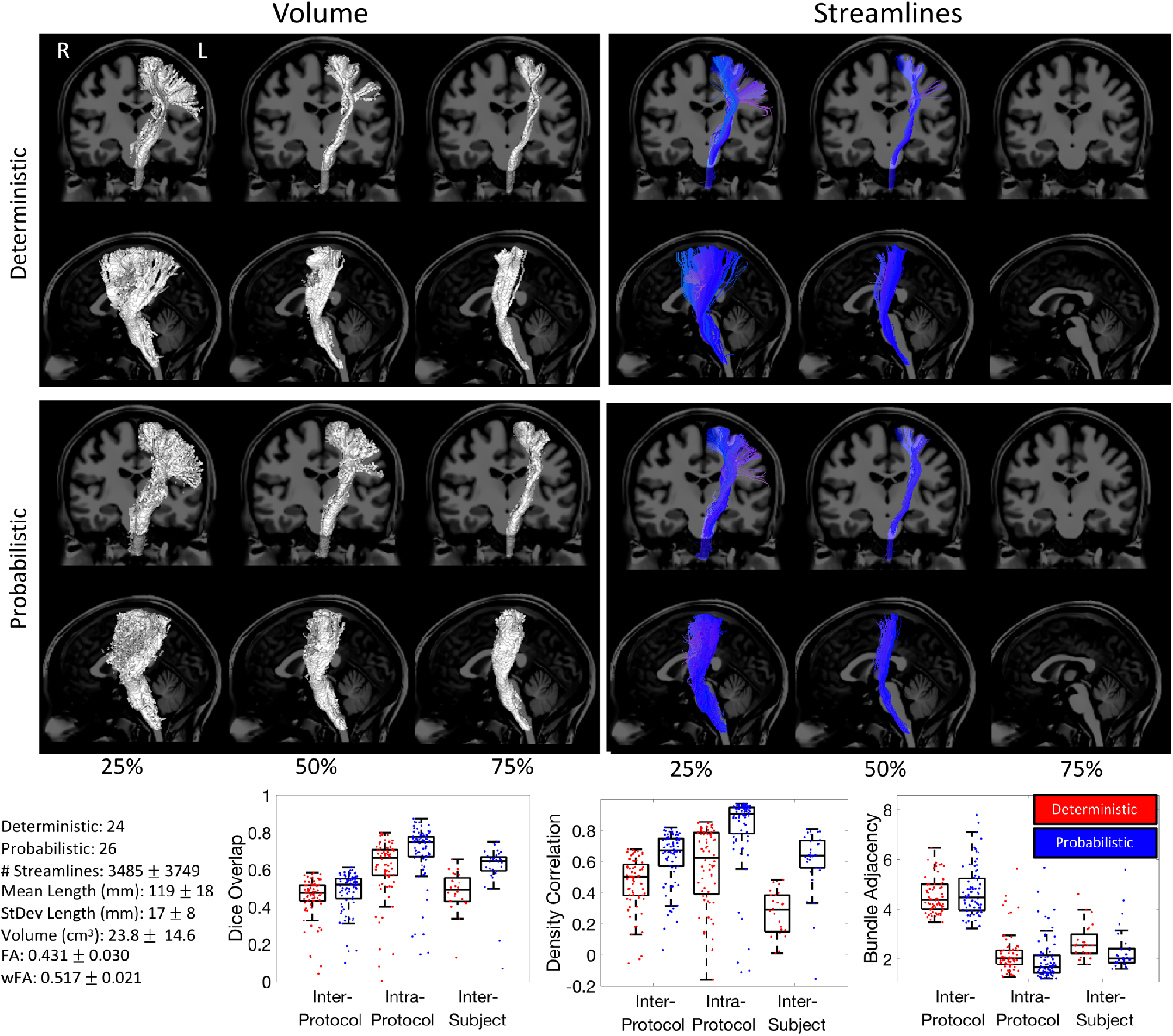
Corticospinal Tract (CST) inter-protocol variability. Renderings show 25%, 50%, and 75% agreement on volume and streamlines for deterministic and probabilistic tractograms. Box-and-whisker plots of Dice overlap, density correlation, and bundle adjacency quantify inter-protocol, intra-protocol, and inter-subject variability (deterministic: red; probabilistic: blue). Each data-point in the plots is derived from the summary statistic of a single submission. Note that there were no streamlines which were common to at least 75% of the protocols.

Quantitative analysis indicates fairly low agreement across protocols. Inter-protocol Dice overlap coefficients largely fall between 0.4 and 0.6 (median Dice of 0.47 and 0.51 for probabilistic and deterministic, respectively), with a larger tail towards much lower Dice values indicating some outlier protocols that are substantially different from others. Protocols show moderate density correlation coefficients (median correlations of 0.51 and 0.67), and an average difference between protocols of >4mm (median bundle adjacency of 4.3mm and 3.9mm). Reproducibility within protocols is much higher, resulting in higher Dice coefficients, higher density correlations, and lower bundle adjacency. The variation across protocols is even greater than the variation across subjects when quantified using Dice overlap. However, the density correlation across protocols is higher than that across subjects, indicating that while the volume overlap decreases, measures of bundle density are more consistent across protocols. Finally, bundle adjacency is higher for inter-protocol analysis than inter-subjects, suggesting that volume-based differences across protocols are greater than volume-based differences across subjects. The quantitative index FA shows a coefficient of variation across protocols of 7% relative to its average value and the density weighted FA shows a variation of 4%.

#### Arcuate Fasciculus (AF)

**Figure 6** shows the results of the inter-protocol analysis for the AF, and **Appendix B** summarizes the descriptive definitions and decisions made in the bundle segmentation workflow. A majority of the extracted bundles agree on the volume occupied by the bundle, with both deterministic and probabilistic submissions showing the characteristic arching shape as the pathway bends from the frontal to temporal lobes. The volume of the 75% agreement is significantly smaller and much more specific than that of the 25% of agreement, occupying only the deep white matter core of this trajectory. Similar results are shown for streamlines. Very few streamlines were agreed upon by 75% of protocols for deterministic tractography, and no single streamline was observed in 75% of probabilistic submissions. Cortical connections show significant variation. Qualitatively, as we become more strict with agreement, the connections become much more refined to the frontal and temporal lobes only, with fewer connections to the parietal cortex.

**Figure 6.**
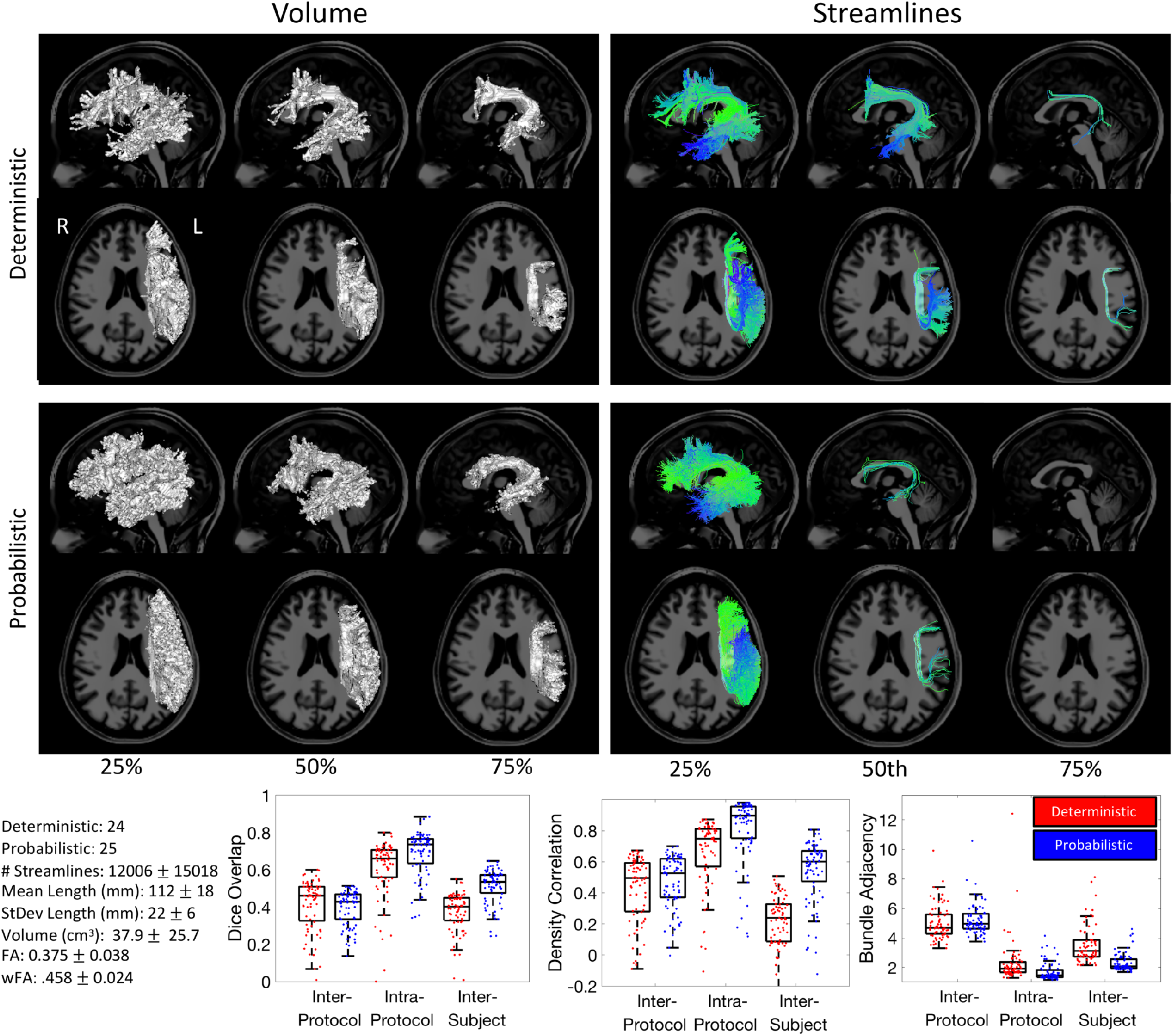
Arcuate Fasciculus (AF) inter-protocol variability. Renderings show 25%, 50%, and 75% agreement on volume and streamlines for deterministic and probabilistic tractograms. Box-and-whisker plots of Dice overlap, density correlation, and bundle adjacency quantify inter-protocol, intra-protocol, and inter-subject variability (deterministic: red; probabilistic: blue). Note that there were no streamlines which were common to at least 75% of the protocols.

Quantitative analyses of similarity and agreement closely follow that of the CST. The Dice overlap indicates relatively poor inter-protocol agreement (median values 0.46 and 0.43 for probabilistic and deterministic, respectively), with a much higher intra-protocol agreement (median of 0.66 and 0.74). However, the inter-protocol overlap is similar to the variation across subjects (0.40 and 0.53). Similar trends are observed for density correlations. In this case, the inter-subject variation is lower than inter-protocol for deterministic, but higher for probabilistic, although both measures are lower than within protocol agreement. Finally, differences across protocols are on average >5mm of distance, whereas the disagreement is much less within protocols and even between subjects. Finally, the coefficient of variation of FA and wFA across protocols is 10% and 5% that of the average FA and wFA, respectively.

#### Corpus Callosum

**Figure 7** shows the results of inter-protocol analysis of the CC, and **Appendix C** presents a summary of the descriptive definitions and decisions made in the bundle segmentation workflow. Most protocols generally agree that this structure takes up a large portion of the cerebral white matter in both hemispheres. While many streamlines were consistent across methods, when looking at the 75% agreement, many submissions do not include lateral projections – although they exist within the dataset – as well as fibers of the splenium (or forceps major) connecting to the occipital lobe and connections to temporal cortex.

**Figure 7.**
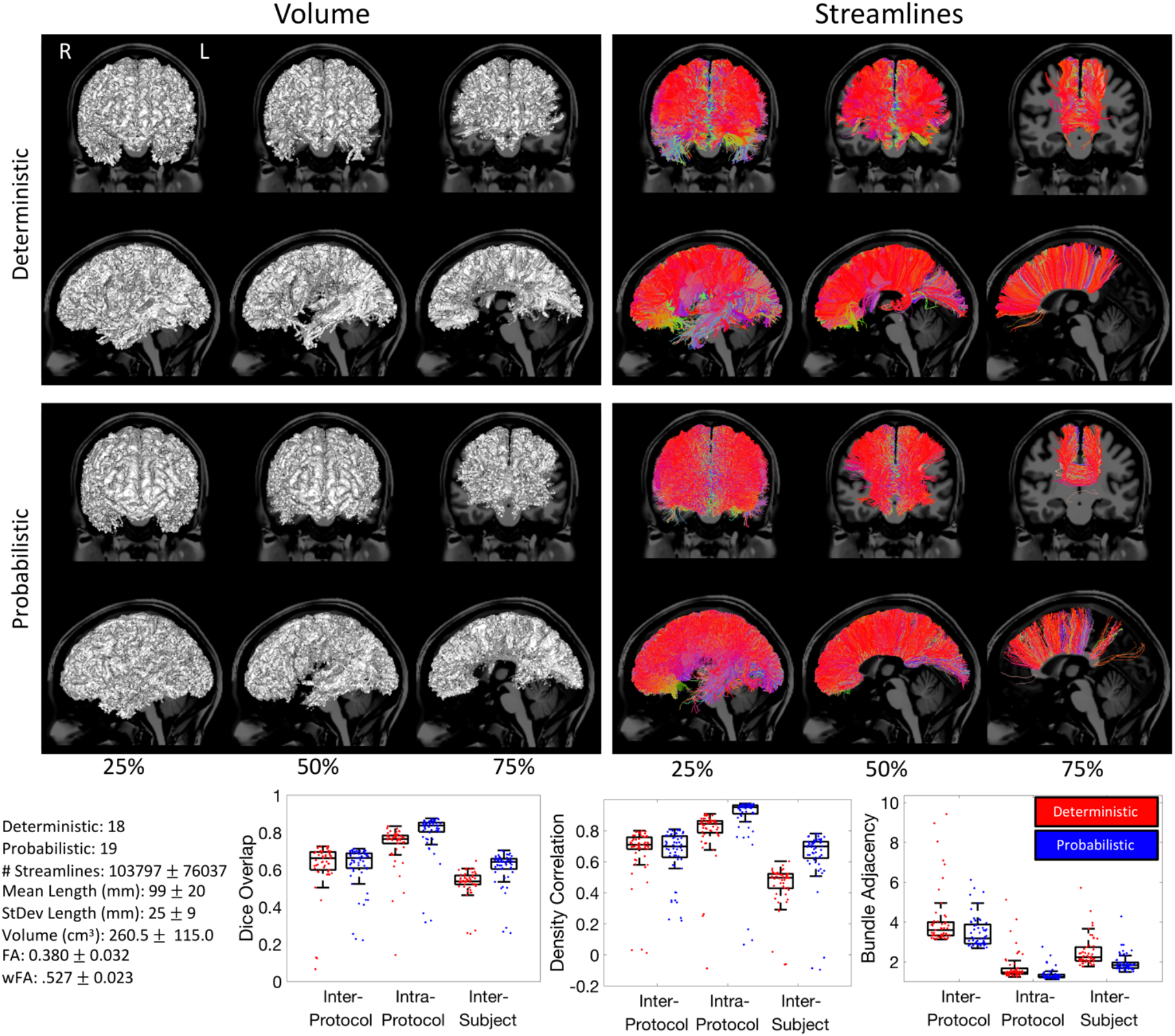
Corpus callosum (CC) inter-protocol variability. Renderings show 25%, 50%, and 75% agreement on volume and streamlines for deterministic and probabilistic tractograms. Box-and-whisker plots of Dice overlap, density correlation, and bundle adjacency quantify inter-protocol, intra-protocol, and inter-subject variability (deterministic: red; probabilistic: blue).

Quantitative analysis shows much higher reproducibility than for the AF and CST, with mean Dice values across protocols of 0.66 and 0.72, which are again lower than intra-protocol reproducibility, but in this case, both slightly higher than that across subjects. The density correlation shows similar trends. Finally, bundle adjacency is higher across protocols than across subjects, with measures indicating that disagreement is generally 3mm or greater across protocols. Even though this structure is quite expansive throughout the white matter, variation across quantitative FA measures are still on the order of 8% and 4% for FA and wFA, respectively.

### Inter-protocol variability

To understand which pathways exhibit the most agreement/disagreement across protocols, intra-protocol volume-based variation measures of Dice overlap, density correlation, bundle adjacency, and Dice streamlines are plotted in **Figure 8.**

**Figure 8.**
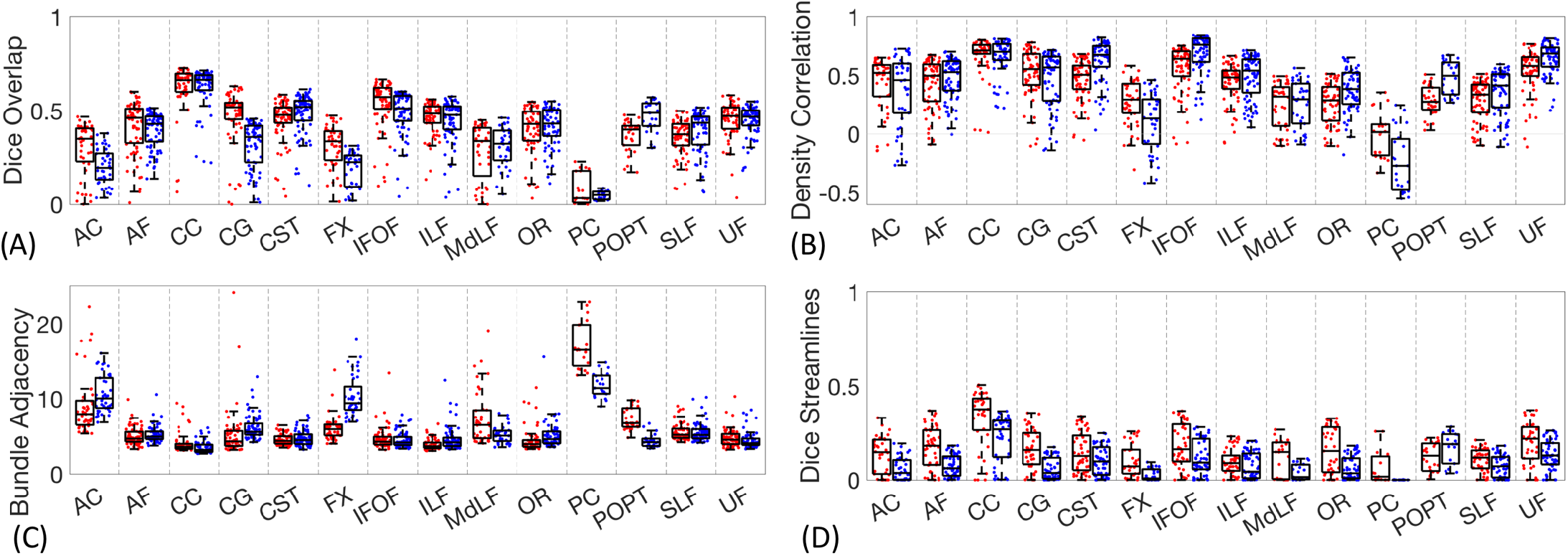
Inter-protocol variability. Dice overlap coefficients, density correlation, bundle adjacency, and Dice streamlines for all studied pathways. Deterministic results shown in red, probabilistic in blue.

There is a fairly large variation across pathways in the overall protocol agreement as measured by Dice volume overlap (**Figure 8A)**. Volume-wise, the most reproducible were the CC, the CST, and the IFOF. Reproducible results from the CC were expected due to its large size and unambiguous location of the CC proper, while the CST is arguably one of the most well-studied tracts. The IFOF, while one of the more controversial fasciculi [8, 9, 27, 28], likely results in higher overlap because it is a long anterior-posterior directed pathway spanning from the occipital to frontal lobe, passing through the temporal stem, a tight and small bottleneck region [29] and most protocols agree that nearly any streamline spanning this extent through a ventral route, will belong to this pathway. In all cases, the overlap across protocols is fairly low, with median values of the CC of 0.66 and 0.72 being the highest among all pathways studied.

The least reproducible structures are those of the commissures, AC and PC, which are largely defined only by a single location along the midline with very little information on their routes or connections. The FX represented a unique case. Many groups submitted the left FX as expected, while others considered the left and right FX as a single structure due to its commissural component. Thus, while it is indeed a small structure, the quantitative value of overlap is overly critical based on qualitative observations.

In agreement with qualitative results, the density correlations (**Figure 8B**) are moderate to high for most pathways, meaning that areas of high streamline density and low streamline density are generally in agreement across protocols. Pathways such as the CC, IFOF, CG, CST, and UF have high agreement in streamline densities, whereas pathways with generally lower number of streamlines and hence lower densities (i.e., PC, and FX) show lower density correlations.

Similar results are observed for dissimilarity (**Figure 8C**). Again, AC, PC, show very large distances of disagreement, along with the FX and in this case the MdLF. For nearly all pathways, the range of disagreements across protocols are most typically on the order of 4-6mm. Looking at Dice overlap of the streamlines (**Figure 8D**), it is immediately apparent that the overlap is very low in all cases, much lower than overlap of volume. For all pathways, a large majority of all comparisons yield streamline Dice coefficients less than 0.2, with many indicating no overlap at all. A trend observed in the streamline comparisons is that the overlap is generally greater for deterministic than probabilistic algorithms.

**Figure 9** shows protocol variability for pathway-specific measures of the mean fractional anisotropy, weighted fractional anisotropy, pathway volume, and pathway length across all protocols. In agreement with results on the CST, AF, and CC, the FA derived from different protocols varies by more than 8-12%, an effect greater than that observed in the literature across study cohorts [30–32]. Weighted-FA (wFA), however, varies much less across protocols (4-7%) and is of greater overall magnitude than the unweighted metric. The volume measurements show that different protocols can result in an order of magnitude difference in pathway volume, an effect observed for all pathways. Finally, pathways with more variation in average streamline length (**Figure 9**) agree well with those with more variation in overlap measures. For example, AC, PC, and FX result in large differences in average length, while protocols on the IFOF consistently agree on the length of this structure.

**Figure 9.**
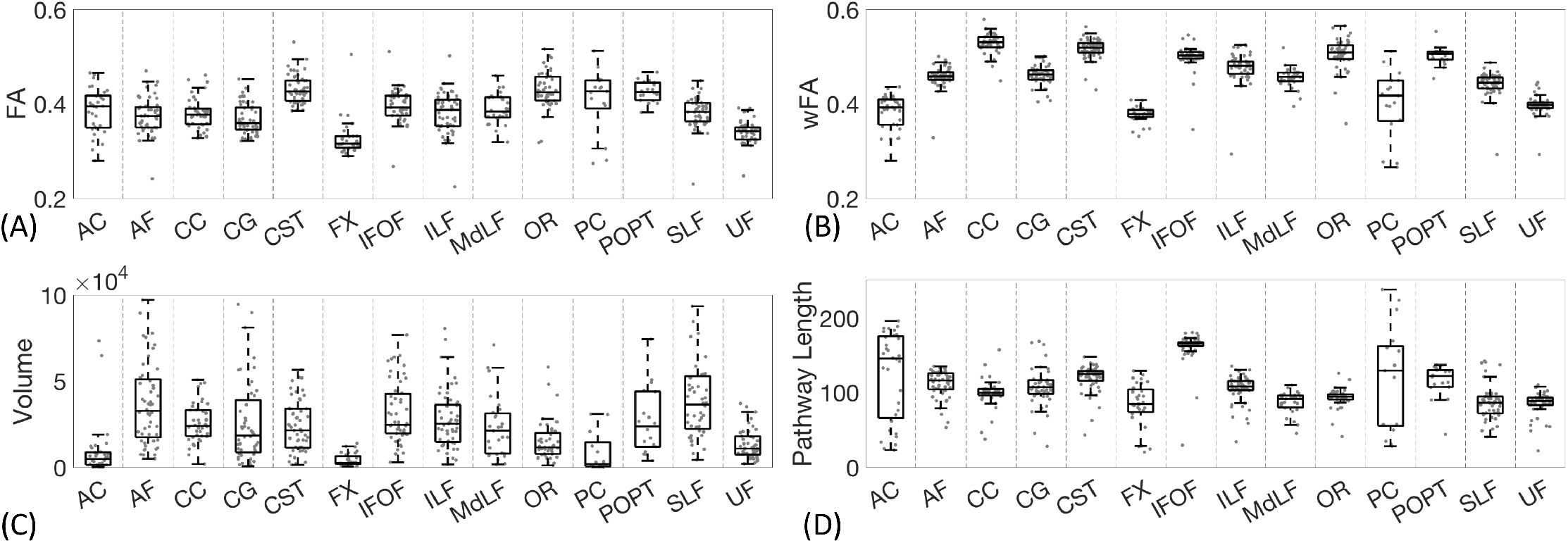
Inter-protocol variation in mean FA, weighted-FA, volume (mm3), and pathway length (mm) for all studied pathways. Note that CC volume is an order of magnitude larger than all other pathways and is shown on a 10^3^ mm^3^ scale.

### Variability within and across pathways

To assess similarity and differences in submissions without a priori user-defined metrics of similarity, we utilized the Uniform Manifold Approximate and Projection (UMAP) [33] technique to visualize all bundle segmentation techniques in a low-dimensional space. The UMAP is a general nonlinear dimensionality reduction that is particularly well suited for visualizing high-dimensional datasets, in this case, on a 2D plane. **Figure 10** shows all submissions, for all pathways, projected on a 2D plane. While there are differences across protocols for a given pathway, all submissions for a given pathway generally cluster together and show similar low-order commonalities, for both probabilistic and deterministic. However, overlap between different pathways does occur in some instances, for example between the SLF and AF (**Figure 10, A**), POPT and CST (**Figure 10, B**), and MLF, ILF, and OR (**Figure 10, C**). This suggests similar low-order representation of some submissions in these pathways.

**Figure 10.**
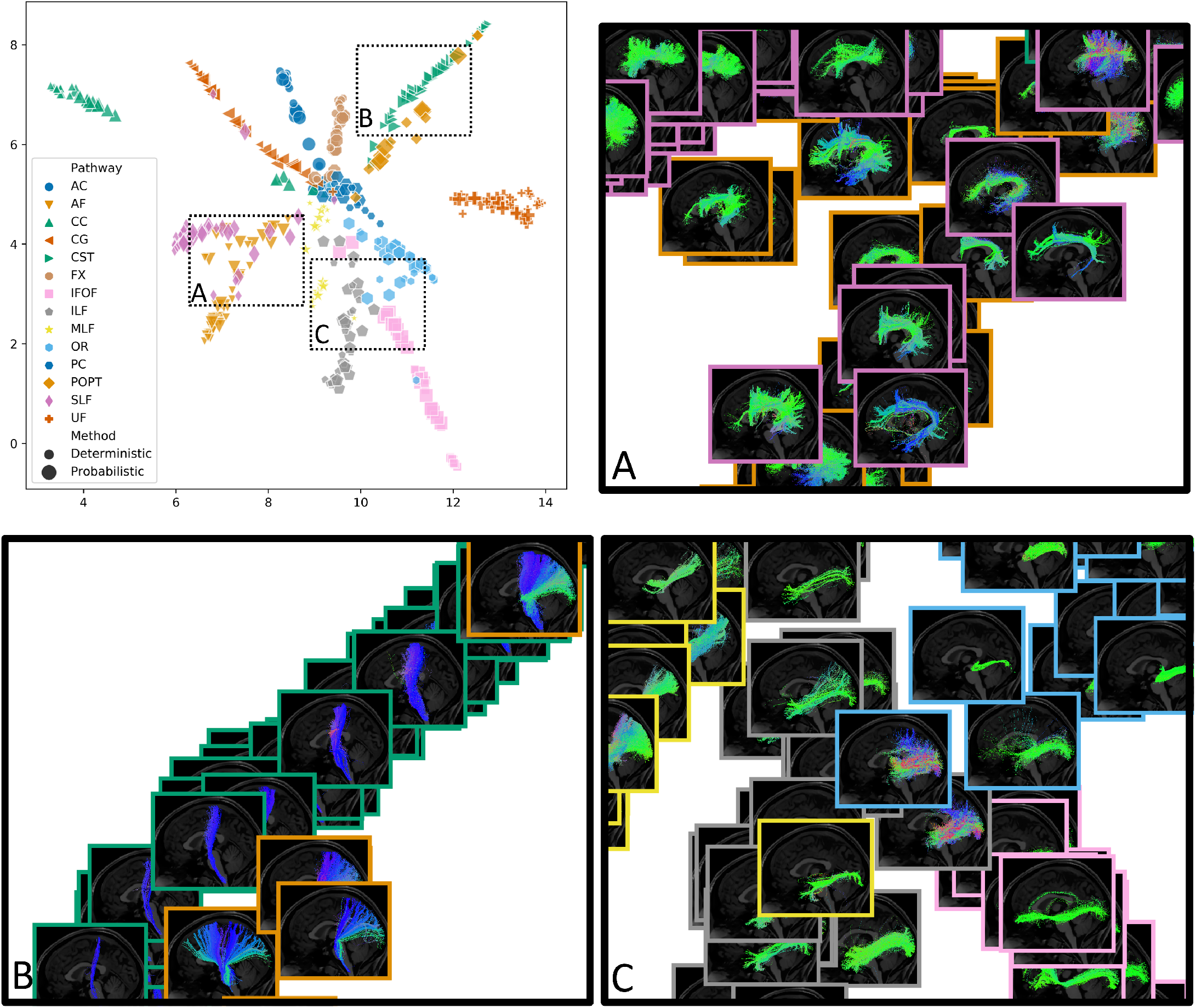
UMAP dimensionality reduction projected bundles onto an un-scaled 2D plane. Object color and shape represent pathways, and object size designates deterministic/probabilistic. While variation exists within pathways and within deterministic/probabilistic streamlines, the white matter pathways generally cluster together in low dimensional space. Insets visualize data points as streamline renderings, and highlight areas where similarity and/or overlap is shown across different pathways.

## Discussion

These results identify and quantify differences and the significant heterogeneity of white matter structures introduced by the use of different protocols for bundle segmentation with tractography. This variability may present difficulties interpreting differences in bundle segmentation results obtained by different labs, or meta-analyses extending and comparing findings from one study to other studies. Additionally, this variation in protocols can lead to variability in quantitative metrics that are greater than true biological variability across populations or subjects and may hinder translation of these techniques from the research laboratory to the clinic.

We propose that a major source of this variation stems from a lack of consensuses on the anatomical definition of pathways [8–11]. There is no standard framework for defining a tract, with some descriptive definitions focusing on the shape and locations of convergence of axons in the deep white matter, while others may focus on specific regions to which fibers connect [9, 11–15]. Consequently, differences, misconceptions, and ambiguities in anatomical definitions and their interpretation may lead to different rules used in the dissection process. For example, workflows used to dissect a bundle range from manual to automated delineation of regions through which streamlines must pass, to shape-based, signal-based, or connection-based methods of segmentation. Importantly, the appropriateness and usefulness of the chosen reconstruction method is application dependent, and no single method is clearly wrong and/or better than the others.

This study was not intended to detract from the value of tractography and bundle segmentation, but rather the aim was to clearly define a current inherent problem and its scope. Looking forward, with a number of well-validated and valuable tools, pipelines, software, and processes at our disposal, it becomes fairly straightforward to modify bundle segmentation protocols to match what we would ultimately strive for in a “consensus definition” of white matter bundles. Thus, instead of describing these results as revealing a problem, we see this as an opportunity, or a call-to-action to harmonize the field of bundle segmentation – both in the nomenclature and definition of white matter pathways, and in the best way to virtually segment these using tractography. Moreover, optimistically, it may be quite useful to have a supply of tools available to dissect and investigate the same white matter bundle in different ways depending on the research question, or the anatomy or functional system under investigation.

Our first main result is that *the inter-protocol agreement is generally poor across protocols for many pathways*, with limited agreement on the brain volume occupied by the pathway. With few exceptions, the average Dice coefficients from both deterministic and probabilistic streamlines were below 0.5, with many considerably lower. For most streamlines, the inter-protocol bundle adjacency is between 4-6 mm, meaning that when protocols disagree, they do so by an average of ~3-5 voxels. Shape and geometry-based measures (i.e., length and volume) of the streamline bundles vary by an order of magnitude across protocols. Consequently, quantitative metrics calculated based on this volume will vary, for example the average FA within a bundle varies by ~8-12% across protocols. Because our analysis was based on the same set of streamlines, these results represent a best-case measure of inter-protocol agreement, and would almost certainly result in increased variability if participants performed their own reconstruction and streamline generation procedures.

Our second main result is that *bundle segmentation protocols have better agreement in areas with high streamline densities*. Measures of streamline density correlation coefficients across submissions are on average >0.5, with few exceptions, which suggests that high density areas in tractograms generally correspond to high density areas of other tractograms, while low density areas correspond to low-density areas (or, in fact, regions with no streamlines). This agrees with observations of 3D density maps where areas of high streamline density are consistently observed in the same location across submissions. These areas of higher streamline density correspond to the core or stem of most of the bundles, generally located in the deep white matter of the brain. Because of this, weighting quantification by streamline density will reduce variability across protocols, for example, wFA varied by ~4-7% across protocols.

Third, we find that the *variability across protocols is greater than the variability within protocols*, and more importantly, similar to (or greater than) the variability across subjects. These results are in agreement with previous studies showing high overlap, high density correlations, and low disagreements *within* a protocol [34–36]. Most importantly, in our study, this represents a worst-case intra-protocol measure. It includes sources of variability related to acquisition (and associated noise and artifacts), registration, reconstruction, and streamline generation – sources of variation which are shown to be still smaller than that across protocols. Thus, while there is little consensus on bundle dissection protocols, a study that uses a consistent protocol has been shown to have the power to reliably detect consistent differences within and across populations; however, there may be limitations in how the findings from a given study can be extended, applied, or compared to others with different protocols.

Fourth, we find that there is *variability per bundle in how much agreement there is across protocols*. The commissural CC has a higher reproducibility due to its large size and very clear anatomical definition, despite more ambiguous definitions of its cortical terminations. However, the PC and AC commissures showed very poor agreement, despite having a very clear location along the midline. This is in part due to smaller sizes, but also scarce literature on the location and connections of the bundles that pass through these regions. CST and IFOF also show moderate agreement across protocols, in part due to their length and at least one location that is moderately specific to these bundles (i.e., the pyramids of the medulla for the CST and the floor of the external capsules for the IFOF). Even here, the Dice overlap across protocols is 0.6 or less, on average. The MdLF and CG show relatively poor agreement. The MdLF is much less studied, and a relatively recent addition to the literature [37, 38], with some disagreement on parietal terminations [11]. The CG is a tract that is likely composed of both longer fibers extending throughout the whole tract, as well as multiple short fibers across its structure which may be both hard for tractography to entirely delineate the long fibers, and hard to capture and constrain segmentation of the shorter fibers that enter and leave throughout [39, 40]. The POPT showed relatively higher agreement. This bundle was included as a relatively ambiguous nomenclature (seen in the literature) of pontine tracts. Whereas both occipito-pontine and parieto-pontine fibers exist, they are not usually defined as a specific tract or fasciculus. Finally, some of the more commonly delineated structures (OR, ILF, SLF, UF) show inter-protocol variabilities somewhere in between, but still exhibit poor-to-moderate volume and streamline overlaps.

For many applications, end-users of bundle segmentation technologies are interested in gross differences in connectivity and location, and what matters is not so much that tracts are reconstructed in their entirety, but that they are not confused with one another. For example, misunderstanding or inapt nomenclature, and/or non-specific constraints in the bundle segmentation process could lead to misidentification of the desired pathway (possibly as another pathway or subset of another pathway) and would lead to confusion in the literature. Based on our results, an experienced neuroanatomist or neuroimager can easily classify the submitted pathways based on visual inspection of the streamlines. Thus, these *inter-protocol bundle segmentations represent the same basic structure*, even if some variability in spatial extent and connections is observed. This is confirmed using an unsupervised data exploration tool for dimensionality reduction, where within-pathway submissions are clearly clustered (for both probabilistic and deterministic algorithms) in low dimensional space. However, there are a few exceptions. Notably, several AF and SLF submissions overlap significantly, which is not unexpected because these have often been defined and/or used interchangeably in the literature [41]. Relatedly, several submissions of the POPT contain a subset of streamlines often assigned as CST, which is again expected because both are often (or can be) described as having parietal connections in common. Finally, several ventral longitudinal systems of fibers (MdLF, OR, ILF, and IFOF) are not clearly separated in this space, suggesting that in many instances they share similar spatial overlap and densities of streamlines across submissions.

Finally, while there is low volume-based agreement, streamline-based agreement is lower still. In fact, many protocols did not agree on a single streamline belonging to a pathway of interest. Protocols agreed on consistently 20% or less of deterministic streamlines and less than 10% of probabilistic streamlines. Put another way, given a set of streamlines from which to select, very few streamlines were consistently determined to be a part of a given pathway across all groups performing the segmentation. With the wide variety of workflows to select streamlines, few streamlines met inclusion criteria associated with cortical connectivity, shape and spatial location, and survived possible exclusion criteria such as filtering based on length, curvature, or diffusions signal, as well as personal preference of the person performing dissection (for example eliminating streamlines to reduce complexity of manual segmentation). Thus, the final main result is that *the measured variability depends on the scale upon which the variability is analyzed*. Protocols show little-to-no agreement in assigning individual streamlines to a pathway, whereas protocols show higher agreement in assessing spatial overlap of pathway, and even higher agreement when taking into account density of streamlines over a volume. This means that while selected streamlines may occupy the same volume, the streamlines that make up this volume are different.Thus, *the effects of this variability are dependent upon how these bundles are ultimately utilized in practice*, and there are a number of ways in which these bundles are used and applied. For this reason, we state that no submissions are inherently “wrong”, and instead emphasize that they are simply “different from one another”.

We have identified variability in the protocols for bundle segmentation, which parallels variability in the literature of other techniques that have been used to elucidate the structure and function of the brain for the last 20 years. These types of disagreements and the challenge in advancing science beyond them are not new to computational neuroanatomy. Indeed, as we look at the history of brain science differences in opinions and associated results can be traced back a long way. Key examples of the inherent variability in anatomical and functional definitions and associated disagreements include the definition and functional specialization of cortical areas [42–44]. Hence, our findings here highlight the complexity of the scientific concepts and the difficulty in making progress towards understanding. The fact that the engineering of new methods needs to be refined because we still have (and have had for over hundreds of years in neuroanatomy) substantial variability in results does not necessarily mean that science is not progressing.

We postulate that the problem stems from two sources (1) the anatomical definition of a white matter pathway and (2) the constraints used to dissect this pathway. The descriptions of the white matter pathways given in the appendix highlight the problem of “definition”. Pathways may be defined by their shape, their endpoints, or by regions through which they pass. Descriptions and definition approaches may vary based on the pathway itself (i.e., some may lend themselves more easily to descriptions of shape rather than endpoints), by the system or functions under investigation, by the training and/or occupation of the researcher/clinician, or by the modality used to define the tract. For example, cadaveric microdissection may facilitate description of fascicular organization and regional descriptions over highly specific lobular connectivity descriptions provided by histological tracers. Further, definitions do not always facilitate binary decision making in the bundle dissection process due to biological reasons. The brain is a complex structure, there are not always hard or unique borders between cortical or subcortical regions, and the location of endpoints or regions may not always be precisely determined. The goal of tractography bundle segmentation then is to recreate these definitions in the bundle dissection process [45]; however, certain algorithms, software packages, and manual pipelines lend themselves more naturally to one type of constraint than the other, and may implement them in different ways or with different levels of precision. Even if a definition has been entirely met, a sensitivity/specificity tradeoff is possible, influenced by potentially every step in the fiber tractography process from acquisition and reconstruction to the final constraints and streamline filtering techniques [46–48].

The question becomes “whose problem is this?”. We propose that there may be shared responsibility on the part of classical anatomists, those developing tractography algorithms, and those implementing or performing segmentations. The endeavor to digitally segment the white matter is predicated upon there being some consensus of what structures are there to be segmented, this is the task of classical neuroanatomists. Next, tractography providers must endeavor to create candidate tractomes that resemble the white matter of the brain as closely as possible, as the resultant tractomes must contain viable anatomy for extraction. Finally, those who perform digital segmentations must decide an appropriate level of precision (sensitivity/specificity) and be clear and precise as they describe the methods of their segmentations as this will permit comparison and refinement between segmentations. This must be an iterative process, utilizing orthogonal information in the form of non-human model brains, micro-dissection, and alternative neuroimaging contrasts, in order to validate the existence and location or connections of a pathway, validate the rules and constraints that allow accurate dissection of this pathway, then iteratively refining the location and/or connections based on knowledge gained through the bundle segmentation process. Thus, we hope that this paper acts as a call to action on two efforts of consensus: both standardization of the anatomical definition (in addition to nomenclature) and the adoption of protocols to fulfill this definition.

Even without a consensus, there could be a convergence towards appropriate, or more specific, nomenclature and clustering of streamlines, or alternative accepted definitions. Additionally, a consensus on the healthy, young adult, individual may not lead to satisfactory results on developing, aging, or diseased populations. The effect of protocols and their adherence to definitions should be investigated in the presence of tumors, on the pediatric and elderly populations, and also with varying acquisition, reconstruction, and streamline generation conditions. While we cannot currently give a recommended dissection protocol for a given pathway, we can recommend good practices to be used in all studies. First, we suggest transparency and explicit descriptions of pathway definition, dissection protocol, and ROIs [3, 49]. Second, understanding and quantifying the intra-protocol variability, for both automatic and manual approaches, is a necessary prerequisite to determine quantification variability and subsequent statistical power. Third, with the knowledge that the dense core of the pathway is consistent across protocols, weighting by density (or a focus on deep white matter, as is common in many statistical analyses [50, 51]) will be more appropriate for evaluating inter-subject difference in microstructural properties, given its smaller inter-site and inter-lab differences. Finally, the results obtained by (and inferences made from) tractography must be interpreted with appropriate level of coarseness, by considering the existence of inter-protocol variability and coarse spatial scale of diffusion MRI measurements. Since some of statistical properties of tractography (streamline counts and densities, and geometry/volume of tracts) have dependency on method selections at this point, it is important to encourage studies by independent groups testing how much conclusions in a single original paper can be generalizable to a different segmentation protocol or datasets.

This study has several limitations which constrain the generalizability of the results. First, there is a low number of subjects and low number of repeats. While automated methods can be run on several hundred subjects using only CPU-hours, this study would have become prohibitive for manual or semi-automated methods with more than 14 pathways over six datasets (84 total possible dissections), and many of these methods would have been under-represented. Next, we did not include a number of pathways with functional relevance in the literature, but chose a sample representative of the commonly studied projection, association, and commissural bundles, and, again, a compromise was made between the number of pathways requested and expected time and effort. Future studies should consider studying pathway sub-divisions specifically, as well as additional major white matter pathways and superficial U-fibers [52]. Further, because we wanted to isolate the effect of bundle segmentation protocols, we forced the use of our own generated streamlines. This may not be optimal for a given segmentation process where streamlines are generated using different parameters or propagation methods, and filtered or excluded in various ways. However, allowing the creation of different streamlines would only increase the variability seen across protocols. Finally, there is no “right” measure to quantify variability [36]. No single measure can paint a complete picture of the similarities and differences of this complex technology across all applications. The measures used in this study were chosen as intuitive quantifications of volume-based, voxel-wise, and streamline-based agreement, as well as measures based on binary volumes and streamline densities. We also quantified measures of geometry which are often used in quantification or to modulate connectivity measures, as well as measures of microstructure within pathways (both weighted and unweighted by densities). Finally, the UMAP approach represents an analysis that is not dependent on user-defined criteria, and allows an intuitive visualization of primary components that explain the data. The best measure of bundle variability is ultimately dependent on how the bundle is used.

## Materials and Methods

We surveyed the protocols for bundle segmentation of 14 white matter bundles, chosen to represent a variety of white matter pathways studied in the literature, including association, projection, and commissural fibers, fibers with clinical and neurosurgical relevance, as well as covering a range from frequently to relatively infrequently studied and/or described in the literature.

We made available the same datasets to be analyzed by a large number of groups in order to uncover variability across analysis teams. To isolate the effects of bundle segmentation from all other sources of variation, we directly provided six sets of whole-brain streamlines (both deterministic and probabilistic) to all collaborators, derived from 3 subjects with scan-rescan data acquired from the Human Connectome Project test-retest database [24]. We extended invitations for collaboration, disseminated data and the protocol with clearly defined tasks, and received streamlines from collaborators for analysis. In addition to streamlines, we requested a written “definition” of the pathways and a description of the constraints used to dissect it. Importantly, this dataset allows us to quantify and compare variability across protocols (inter-protocol), variability within protocols (intra-protocol), and variability across subjects (inter-subject). Detailed procedures are provided in supplementary material.

### Data and Protocol

The diffusion data for this study were selected from the Human Connectome Project test-retest database [24]. A total of three subjects (HCP IDs: 144226, 103818, 783462) were chosen that had repeat diffusion MRI scans, resulting in six high-quality datasets, free of any significant artifacts. This dataset was chosen as a compromise between quantification and inclusivity - the use of this small database still provides enough information to detect and quantify the variability among results with great enough participation across laboratories and scientists.

Collaborators were not informed that the six datasets represented only three subjects in order to not bias intra-protocol analysis. Distortion, motion correction and estimation of nonlinear transformations with the MNI space was performed using the HCP preprocessing pipelines [24]. Whole-brain tractograms were generated using the DIPY-based Tractoflow processing pipeline [53, 54], producing both deterministic and probabilistic sets of streamlines to be given to participants. Importantly, to be as inclusive as possible to all definitions and constraints, streamlines were not filtered in any way. Streamlines were separated into left, right, and commissural fibers in order to minimize file sizes. Also provided were the b0 images, Fractional Anisotropy (FA) maps [55], directionally-encoded color maps [55], T1 weighted images, and masks for the cerebrospinal fluid, gray matter, and white matter [55].

The task given to collaborators was (see supplementary material) to dissect 14 major white matter pathways on the left hemisphere on the six diffusion MRI datasets provided. Collaborators were free to choose either deterministic or probabilistic streamlines, and free to utilize any software they desired. In order to maximize the quality of submitted results, investigators did not have to provide segmentations for all pathways if they did not have protocols or experience in some areas.

### Submissions

For submission, we asked for a written definition of the white matter bundles, a description of the protocol to dissect these pathways, all code and/or temporary files in order to facilitate reproducibility of methods, and finally the streamline files themselves. Quality assurance was performed on file organization, naming conventions, and streamline spatial attributes, and visual inspection was performed for all streamlines of all subjects. Tools for quality assurance (QA) can be found at (https://github.com/scilus/scilpy).

### Pathway-specific Analysis

For all pathways, we focused on quantifying volume-based and streamline-based similarities and differences in the dissected bundles across protocols. Qualitatively, we assessed volume overlap and streamline overlap. Volume overlap was displayed as the volume of voxels in which 25%, 50%, and 75% of all protocols agreed that a given voxel was occupied by the pathway under investigation. Similarly, we viewed the individual streamlines in which 25%, 50%, and 75% of all protocols agreed that this streamline is representative of a given pathway. These qualitative observations were shown as volume-renderings or streamlines visualizations directly.

Next, quantitative analysis used three voxel-based measures (based on volume and streamline density) and one streamline-based measure [36]. The Dice overlap coefficient, density correlation coefficient, bundle adjacency, and streamline Dice overlap are illustrated in **Figure 4**. Dice overlap measures the overall volume similarity between two binarized bundles (i.e., all voxels that contain a streamline), by taking twice the intersection of two bundles divided by the union of both bundles. A value of 1 indicates perfect overlap, a value of 0 indicates no overlap. The density correlation coefficient is a measure of the Pearson’s correlation coefficient obtained from the streamline density maps. This provides insight into not only overlap, but also agreement in streamline density. Bundle adjacency is a volume-based metric that describes the average distance of disagreement between two bundles. This was calculated by taking all non-overlapping voxels from one bundle, and calculating the nearest distance to the second bundle (and repeating from the second to the first bundle) and taking the average of these distances. By defining this metric, we are using a convenient symmetric distance between two binary volumes, which is a modification of the Hausdorff distance. A value of 3mm, for example, indicates that when the bundles disagree, they are an average of 3mm apart. Finally, streamline Dice is the streamline-equivalent of Dice overlap. Because all submissions for a given subject were derived from the same set of whole-brain streamlines, we had the ability to quantify whether an individual streamline was common to both submitted bundles. Streamline Dice was calculated by taking the total amount of streamlines common to both protocols (i.e., intersection) divided by the total number of unique streamlines in both bundles (i.e., union). Again, a value of 1 indicates that all streamlines are exactly the same, a value of 0 indicates no overlap in streamlines. Note that this final measure can be calculated only for datasets that are derived from the same original set of streamlines.

### Quantifying variability across protocols

The measures introduced above were used to quantify variability across protocols (inter-protocol), variability within protocols (intra-protocol), and variability across subjects (inter-subject), with separate analyses for deterministic and probabilistic results. Below, we describe these three levels of variability assuming there were “N” submissions for a given pathway.

For inter-protocol variability, each bundle was compared to its counterpart as produced by each of the other N-1 protocols, and the results averaged, representing the average similarity/dissimilarity of that protocol with all others. This was done for all N submissions, for all 3 subjects, resulting in Nx3 data-points for each pathway.

For intra-protocol variability, we aimed to compare the same protocol performed on the same subject. For each of the N submissions, we calculated the similarity/dissimilarity measures with respect to the same submission on the repeated scan. This was repeated for all subjects, resulting in again Nx3 data-points for each pathway. A “precise” measure of intra-protocol variability would have been possible if the same set of streamlines had been provided twice for each subject. Instead, the study used scan/re-scan data to measure not only intra-protocol variability, but *the variability of everything up to, and including protocol*. Thus, this measure includes acquisition variability (i.e., noise and possible artifacts), registration (to a common space), reconstruction, and generation of whole brain streamlines.

Finally, for inter-subject variability, we sought to characterize how similar/dissimilar a bundle is across subjects within a single protocol. All streamlines were normalized to MNI space using nonlinear registration (*antsRegistrationSyn)* [56] of the T1 image to the MNI ICBM 152 asymmetric template [57]. For each of N protocols, the agreement measures were calculated from subject 1 to subject 2, from subject 2 to subject 3, and from subject 1 to subject 3, again resulting in Nx3 data-points for each pathway.

## Supporting information

Number of submissions per pathway

Visualization of Pathways

Results for all pathways

Protocol given to collaborators

## Appendix A Cortico Spinal Tract (CST)

The CST is the major descending tract that mediates voluntary skilled movements [58, 59]. At its most basic, this tract is a pathway of fibers coursing primarily from the motor cortex down the spinal cord. Despite this apparent simplicity, dissecting this tract can be quite variable. Moderately increasing the complexity of the definition, the CST can be (unanimously) described as starting from the cortex, traveling through the corona radiata, converging into the internal capsule, continuing into the brainstem through the medulla, and finally extending to the spinal cord. Decisions to be made include choosing specific cortical terminations (which span both frontal and parietal lobes) and how these are delineated, selecting regions through which the streamlines must pass (“cortex to medulla” or “cortex to lower brainstem” or “motor cortex to medulla”), and implementing additional inclusion and exclusion regions throughout the extent of the pathway to further refine where it goes and where it does not go. Adding further ambiguity, the CST together with the corticobulbar tract make up the pyramidal tract, and because these are not easily (or not possibly) separated due to inherent tractography limitations and field of view restrictions, these have sometimes been used interchangeably and/or incorrectly in the literature. In this study, the CST was divided into precentral and postcentral divisions based on endpoints, hand-foot-face divisions based on regions of interest, anterior-posterior-central-cingulate divisions based on endpoints, combined/separated with ascending pathways with thalamic synapses, as well as combined/separated with the peri-Rolandic component based on endpoints, and divided into lateral and anterior components based on definition (but not dissected).

## Appendix B Arcuate Fasciculus (AF)

The AF plays a key role in language processing. This is an association tract that is well-understood to connect Wernicke’s area (somewhere in the posterior temporal lobe) to Broca’s area (located in the inferior frontal lobe). It gets its name (Latin for *curved* bundle) from the distinctive arch shape it makes as it curves from the anterior-posterior direction in the frontal-parietal cortex ventrally into the temporal cortex around the Sylvian fissure (lateral sulcus) [60, 61]. This description of the AFs shape is generally agreed upon. A third area (inferior parietal lobule) is also traditionally included in this tract’s connections, representing the pathway that Geschwind postulated to be damaged in conduction aphasia [60]. For this reason, many descriptions include multiple segments of the AF - a direct pathway traversing the entire tract from temporal to frontal lobes, and an indirect pathway of shorter fibers connecting temporal to parietal to frontal lobes. Consequently, the AF can be described as connecting a number of areas of the perisylvian cortex of the frontal, parietal, and temporal lobes. To further complicate the literature, because the AF is a dorsal longitudinal system of tracts, it is occasionally considered to be part of the SLF system of tracts [41, 62] and considered synonymous or used interchangeably in the literature [41]. For these reasons, we hypothesized that we would see large variability when giving collaborators the task to “segment the arcuate fasciculus”. Variability is observed due to differences in defining the location and method of delineating Wernicke’s and Broca’s areas, or selection of regions to capture the arch-like shape. Approximately 1/5 of submissions indicated dividing the AF into the long direct segment (often described as more medially located), and the anterior and posterior indirect segments (described as laterally located shorter segments).

## Appendix C Corpus Callosum (CC)

The CC is the largest, and arguably most easily recognizable, white matter structure of the brain. This structure is not a single tract, but rather a commissure, composed of axons coursing in the left-right orientation at the midline, and interconnecting the cerebral cortex of the two hemispheres. Many subdivisions of the CC have been proposed [63] with most partitioning the CC based on axon location in the mid-sagittal section. Most commonly, subcomponents are rostrum, genu, body, isthmus, splenium, and (sometimes) tapetum, although others include genu, splenium, and callosal body, or anterior, mid-anterior, central, mid-posterior, and posterior based on (FreeSurfer) parcellation schemes. Alternative subdivisions included separating according to the major lobes of the brain (frontal, parietal, occipital, and temporal) or numerical subdivisons (ranging between 5 and 12) based on cadaveric and histological dissections [64], or homologous connections, or clusters of fibers. Common to all protocols is the large, easily distinguishable region near the midline. Constraints, decisions, and filters include choices of where these bundles cannot go (various temporal lobe regions, through or near subcortical structures, cingulum and parahippocampal gyri, etc), filtering by connection regions or lengths, or rules enforcing homologous connections.

## Acknowledgments

This work was conducted in part using the resources of the Advanced Computing Center for Research and Education at Vanderbilt University, Nashville, TN. KS, BL, CH were supported by the National Institutes of Health under award numbers R01EB017230, and T32EB001628, and in part by ViSE/VICTR VR3029 and the National Center for Research Resources, Grant UL1 RR024975-01. This work was also possible thanks to the support of the Institutional Research Chair in NeuroInformatics of Université de Sherbrooke, NSERC and Compute Canada (MD, FR). MP received funding from the European Union’s Horizon 2020 research and innovation programme under the Marie Skłodowska-Curie grant agreement No 754462. The Wisconsin group acknowledges the support from a core grant to the Waisman Center from the National Institute of Child Health and Human Development (IDDRC U54 HD090256). NSF OAC-1916518, NSF IIS-1912270, NSF IIS-1636893, NSF BCS-1734853, NIH NIBIB 1R01EB029272-01, and a Microsoft Faculty Fellowship to F.P. LF acknowledges the support of the Cluster of Excellence *Matters of Activity. Image Space Material* funded by the Deutsche Forschungsgemeinschaft (DFG, German Research Foundation) under Germany’s Excellence Strategy – EXC 2025. SW is supported by a Medical Research Council PhD Studentship UK [MR/N013913/1]. The Nottingham group’s processing was performed using the University of Nottingham’s Augusta HPC service and the Precision Imaging Beacon Cluster. JPA, MA and SMS acknowledges the support of FCT - Fundação para a Ciência e a Tecnologia within CINTESIS, R&D Unit (reference UID/IC/4255/2013). MM was funded by the Wellcome Trust through a Sir Henry Wellcome Postdoctoral Fellowship [213722/Z/18/Z]. EJC-R is supported by the Swiss National Science Foundation (SNSF, Ambizione grant PZ00P2 185814/1). CMWT is supported by a Sir Henry Wellcome Fellowship (215944/Z/19/Z) and a Veni grant from the Dutch Research Council (NWO) (17331). FC acknowledges the support of the National Health and Medical Research Council of Australia (APP1091593 and APP1117724) and the Australian Research Council (DP170101815). NSF OAC-1916518, NSF IIS-1912270, NSF IIS-1636893, NSF BCS-1734853, Microsoft Faculty Fellowship to F.P. D.B. was partially supported by NIH NIMH T32-MH103213 to William Hetrick (Indiana University). CL is partly supported by NIH grants P41 EB027061 and P30 NS076408 “Institutional Center Cores for Advanced Neuroimaging. JYMY received positional funding from the Royal Children’s Hospital Foundation (RCH 1000). JYMY, JC, and CEK acknowledge the support of the Royal Children’s Hospital Foundation, Murdoch Children’s Research Institute, The University of Melbourne Department of Paediatrics, and the Victorian Government’s Operational Infrastructure Support Program. C-HY is grateful to the Ministry of Science and Technology of Taiwan (MOST 109-2222-E-182-001-MY3) for the support. LC acknowledges support from CONACYT and UNAM. ARM acknowledges support from CONACYT. LJO, YR, and FZ were supported by NIH P41EB015902 and R01MH119222. AJG was supported by P41EB015898. NM was supported by R01MH119222, K24MH116366, and R01MH111917. This project has received funding from the European Union’s Horizon 2020 Research and Innovation Programme under Grant Agreement No. 785907 & 945539 (HBP SGA2 & SGA3), and from the ANR IFOPASUBA-19-CE45-0022-01. PG, CR, NL and AV were partially supported by ANID-Basal FB0008 and ANID-FONDECYT 1190701 grants. We would like to acknowledge John C Gore, Hiromasa Takemura, Anastasia Yendiki, and Riccardo Galbusera for their helplful suggestions regarding the analysis, figures, and discussions.

