## Supplementary material for "Tractography dissection variability: what happens when 42 groups dissect 14 white matter bundles on the same dataset?": Number of submissions per pathway

**what happens when 42 groups**

Supplementary Information

Abbreviations: Superior Longitudinal Fasciculus (SLF), Arcuate Fasciculus (AF), Optic Radiation (OR), Corticospinal Tract (CST), Cingulum (CG), Uncinate Fasciculus (UF), Corpus Callosum (CC), Middle Longitudinal Fasciculus (MdLF), Inferior Fronto-Occipital Fasciculus (IFOF), Inferior Longitudinal Fasciculus (ILF), Fornix (FX), Anterior Commissure (AC), Posterior Commissure (PC), and Parieto-Occipital Pontine Tract (POPT).

### Number of Submissions

This work resulted in 57 unique bundle segmentation protocols from the provided set of deterministic or probabilistic streamlines. Not all pathways were represented by all protocols, thus there are varying numbers of submissions for each pathway. Additionally, six groups submitted protocols that generated their own streamlines. These were not included in the current work in order to isolate effects of bundle dissection protocol alone. The number of submissions for each pathway is shown in Supplementary Table 1.

Supplementary Table 1. Number of submitted pathways.

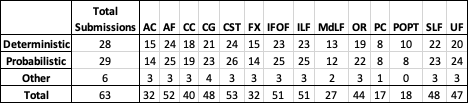
