## Supplementary material for "Tractography dissection variability: what happens when 42 groups dissect 14 white matter bundles on the same dataset?": Visualization of Pathways

**what happens when 42 groups**

Supplementary Information

Abbreviations: Superior Longitudinal Fasciculus (SLF), Arcuate Fasciculus (AF), Optic Radiation (OR), Corticospinal Tract (CST), Cingulum (CG), Uncinate Fasciculus (UF), Corpus Callosum (CC), Middle Longitudinal Fasciculus (MdLF), Inferior Fronto-Occipital Fasciculus (IFOF), Inferior Longitudinal Fasciculus (ILF), Fornix (FX), Anterior Commissure (AC), Posterior Commissure (PC), and Parieto-Occipital Pontine Tract (POPT).

### Pathway Visualization

The following figures demonstrate variability in bundle segmentation protocols for the 14 selected pathways, separated based on projection, association, and commissural pathways. The dissected bundles are shown on the same subject, from the same set of whole brain streamlines. In each case, eight randomly selected bundle segmentation approaches for each pathway are shown as segmented streamlines and rendered as 3D streamline density maps. Variations in size, shape, density, and connectivity are qualitatively apparent. Random selections generated independently for each pathway.

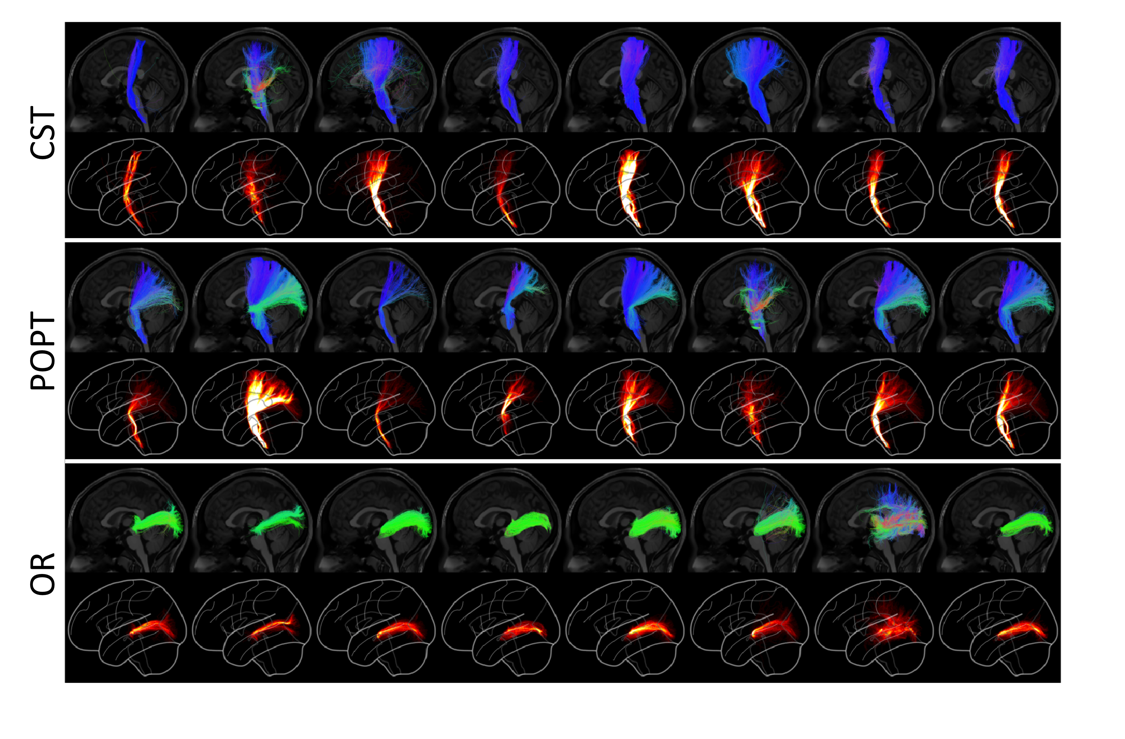

**Supplementary Figure 1.** Streamline and 3D density rendering of projection pathways dissected from the probabilistic set of streamlines.

**Supplementary Figure 2.** Streamline and 3D density rendering of projection pathways dissected from the deterministic set of streamlines.

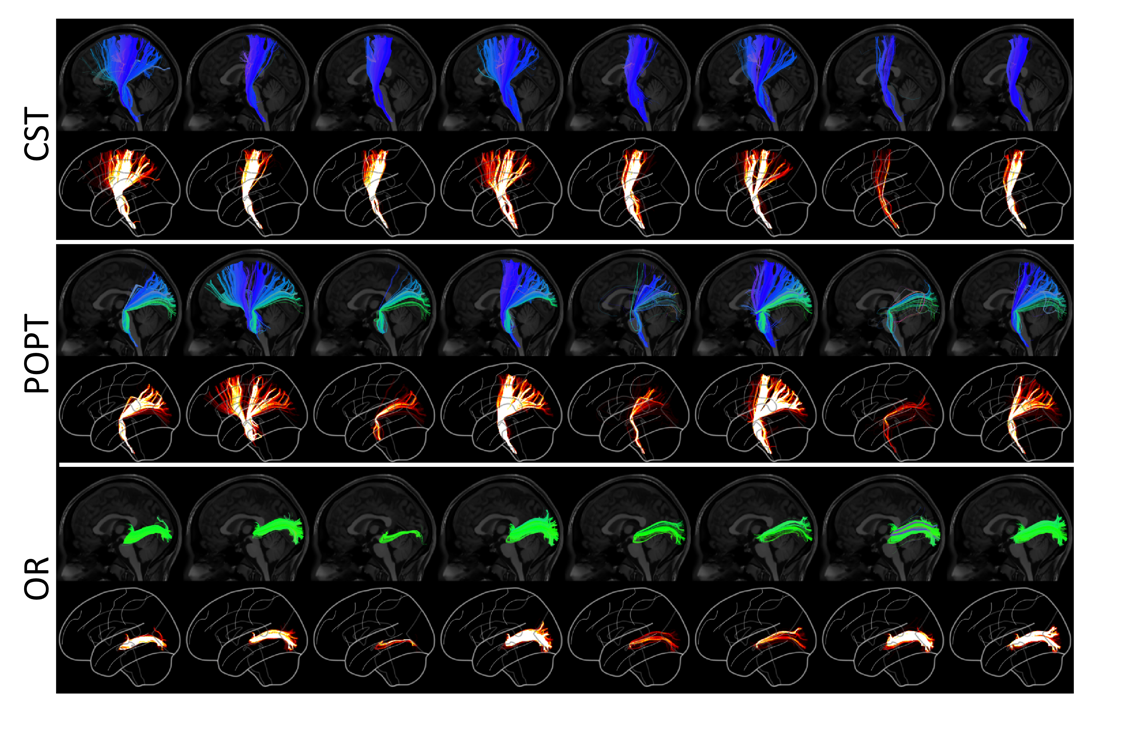

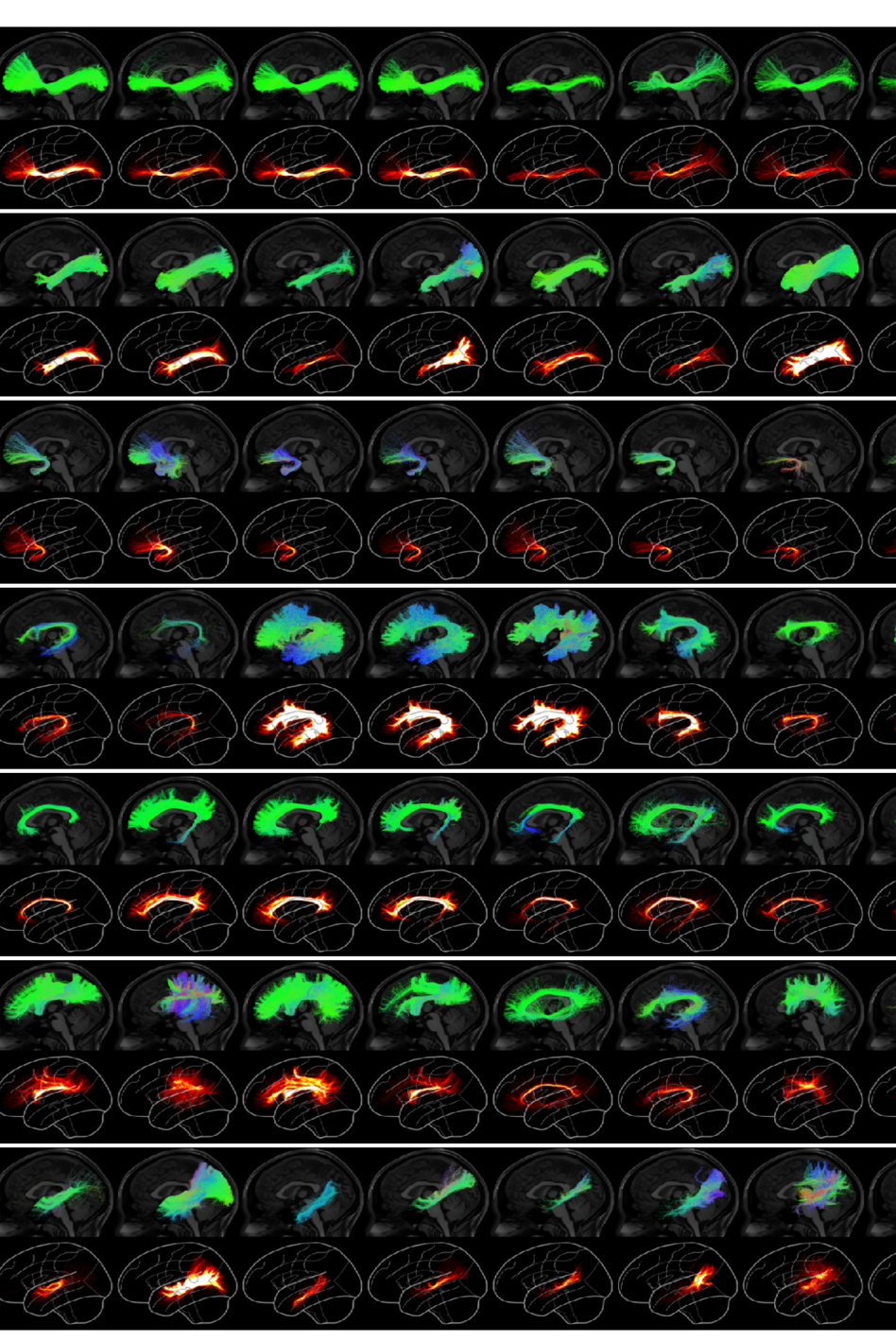

**Supplementary Figure 3.** Streamline and 3D density rendering of association pathways dissected from the probabilistic set of streamlines.

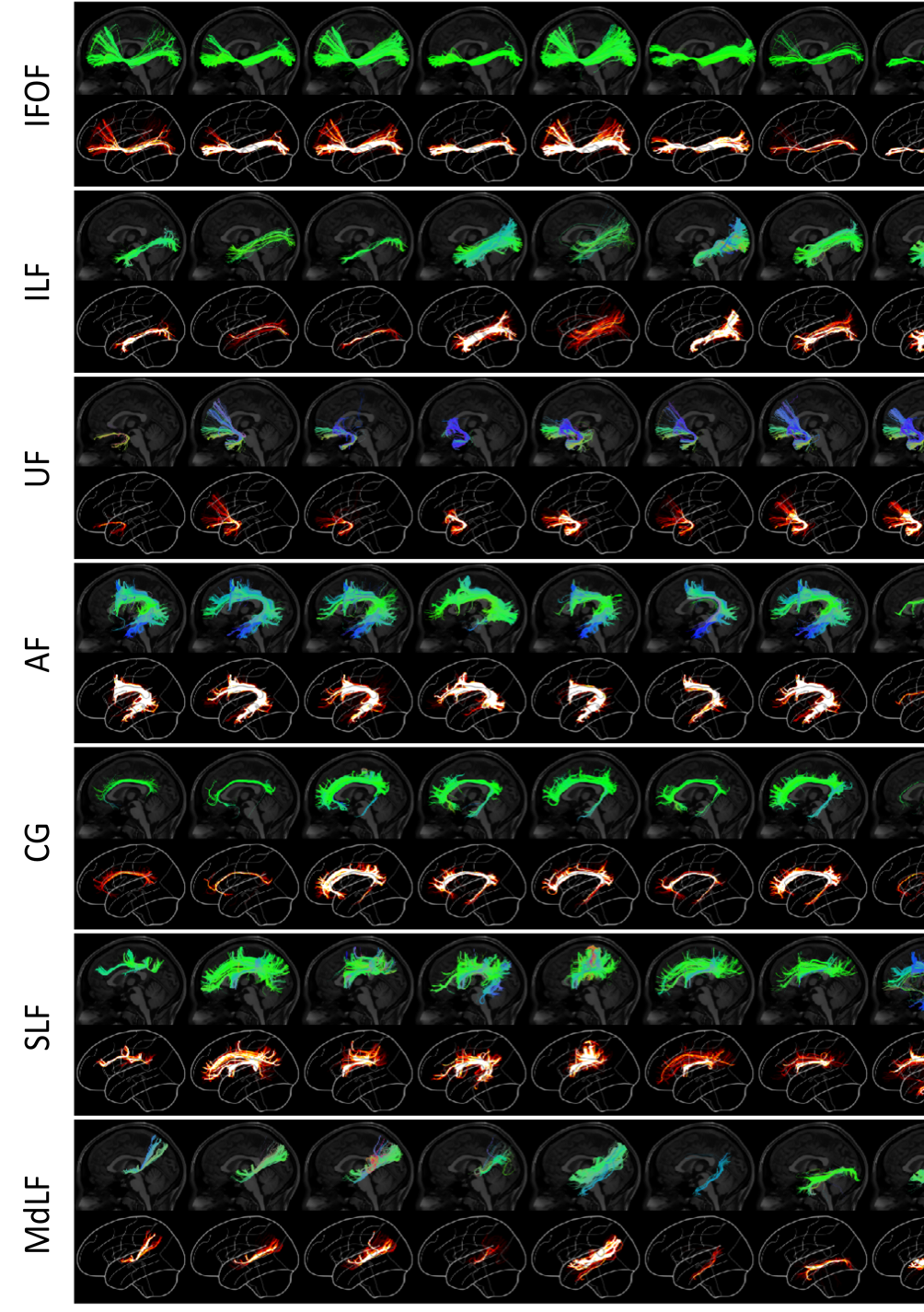

**Supplementary Figure 4.** Streamline and 3D density rendering of association pathways dissected from the deterministic set of streamlines.

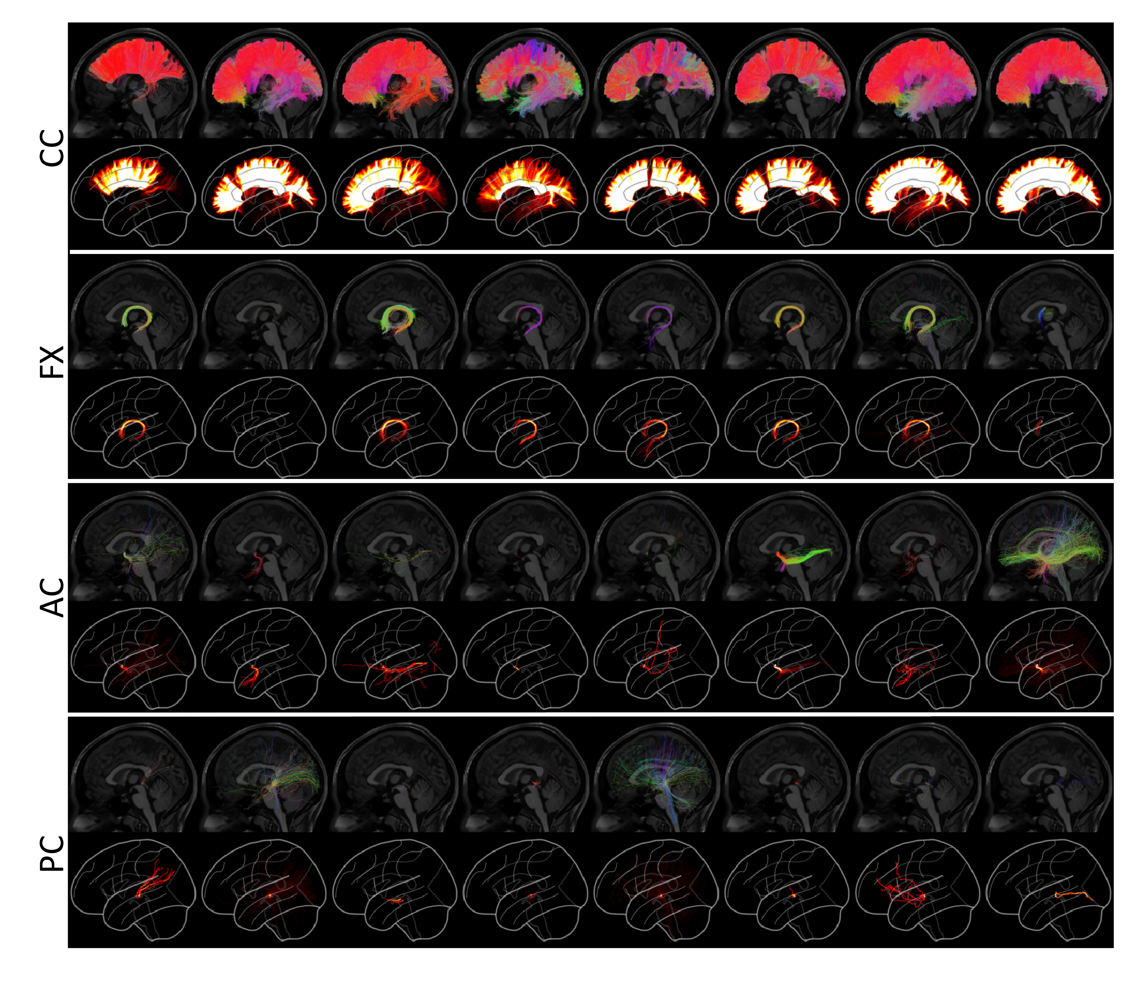

**Supplementary Figure 5.** Streamline and 3D density rendering of commissural pathways dissected from the probabilistic set of streamlines.

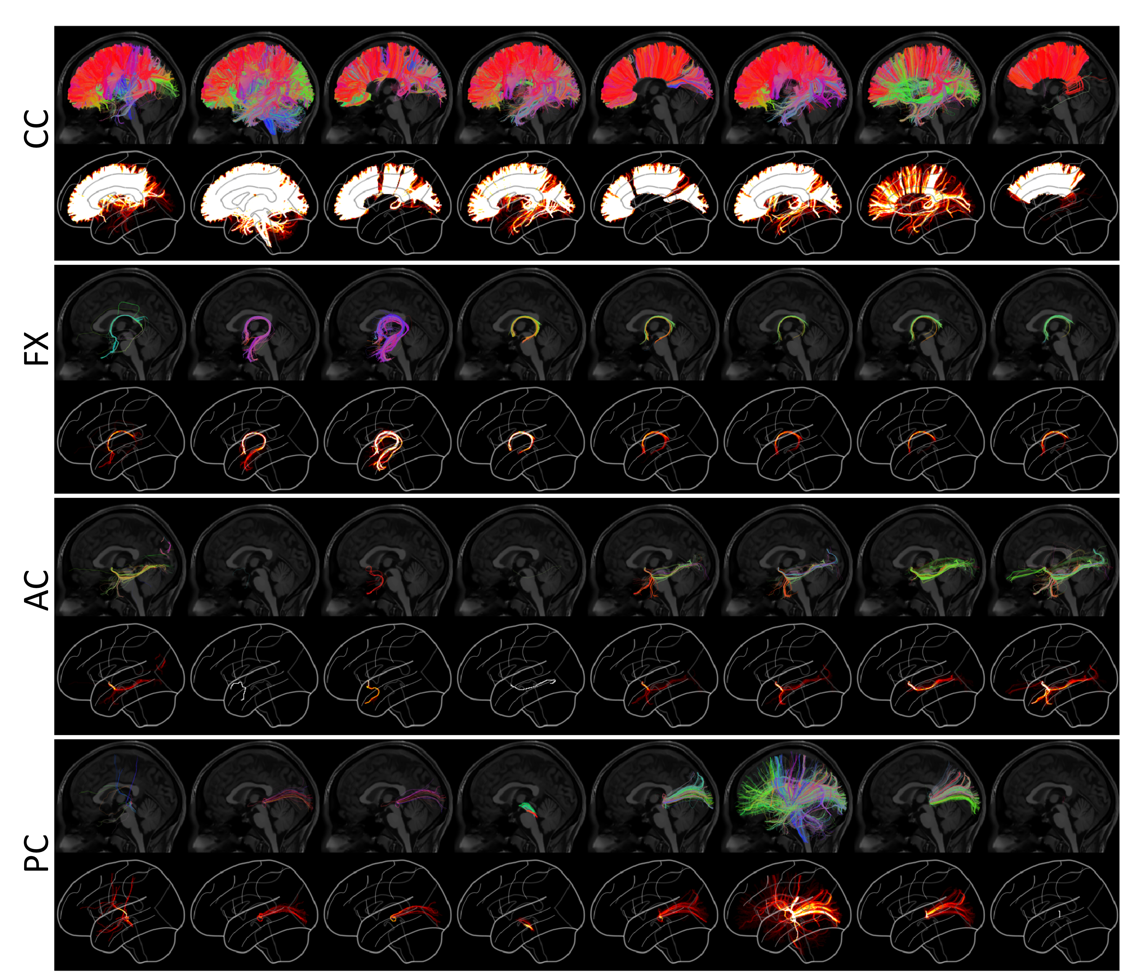

**Supplementary Figure 6.** Streamline and 3D density rendering of commissural pathways dissected from the deterministic set of streamlines.
