## Supplementary material for "Tractography dissection variability: what happens when 42 groups dissect 14 white matter bundles on the same dataset?": Results for all pathways

**what happens when 42 groups**

Supplementary Information

Abbreviations: Superior Longitudinal Fasciculus (SLF), Arcuate Fasciculus (AF), Optic Radiation (OR), Corticospinal Tract (CST), Cingulum (CG), Uncinate Fasciculus (UF), Corpus Callosum (CC), Middle Longitudinal Fasciculus (MdLF), Inferior Fronto-Occipital Fasciculus (IFOF), Inferior Longitudinal Fasciculus (ILF), Fornix (FX), Anterior Commissure (AC), Posterior Commissure (PC), and Parieto-Occipital Pontine Tract (POPT).

### Pathway-Specific Results

The following figures quantify inter-protocol variability. For each pathway, renderings show 25%, 50%, and 75% agreement on volume and streamlines for deterministic and probabilistic tractograms. Box-and-whisker plots of Dice overlap, density correlation, and bundle adjacency quantify inter-protocol, intra-protocol, and inter-subject variability (deterministic: red; probabilistic: blue). Each data-point in the plots is derived from the summary statistic of a single submission.

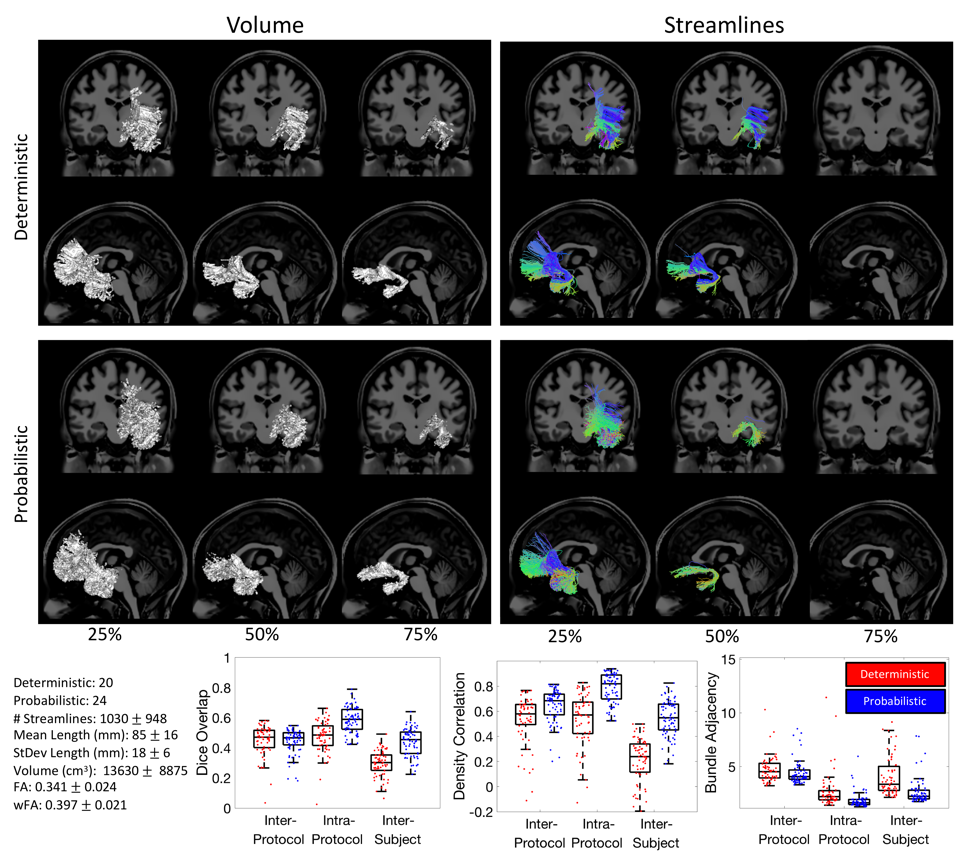

**Supplementary Figure 7.** UF Inter-protocol variability.

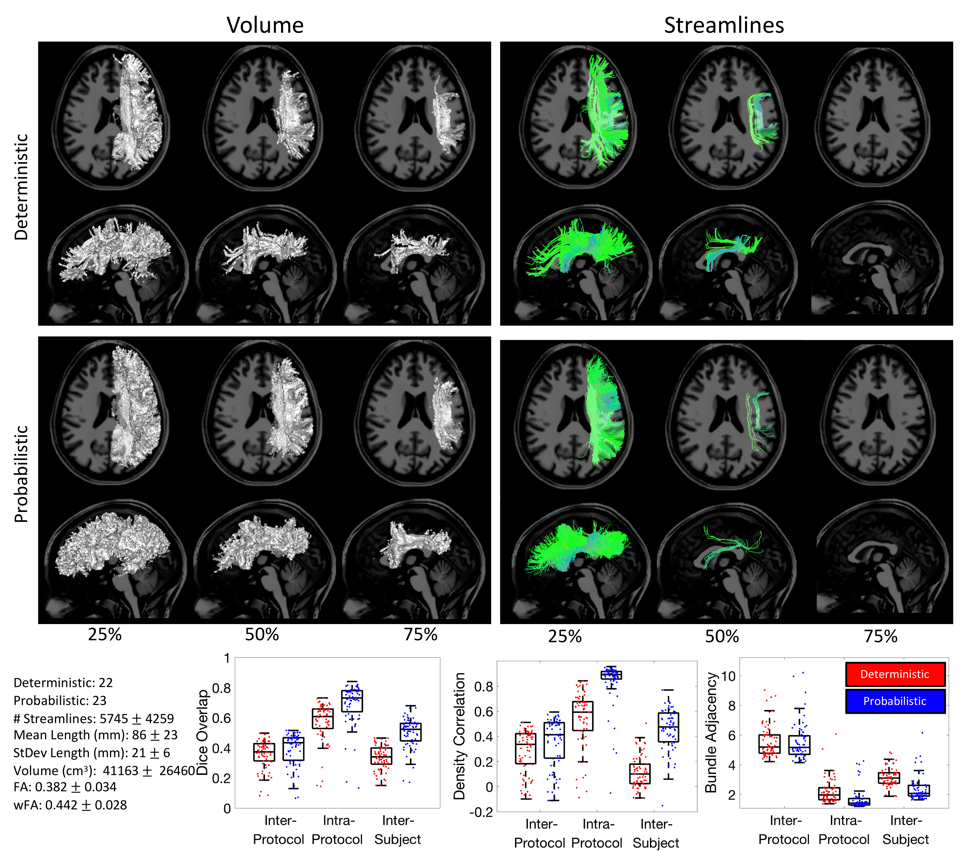

**Supplementary Figure 8.** SLF Inter-protocol variability.

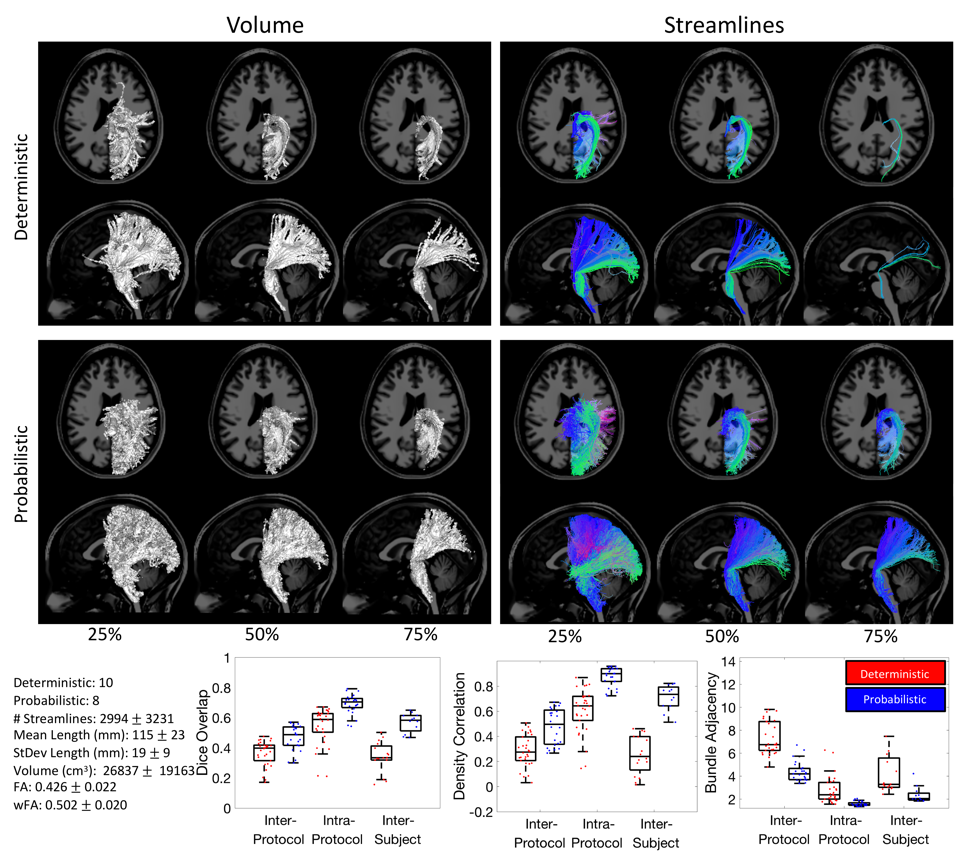

**Supplementary Figure 9.** POPT Inter-protocol variability.

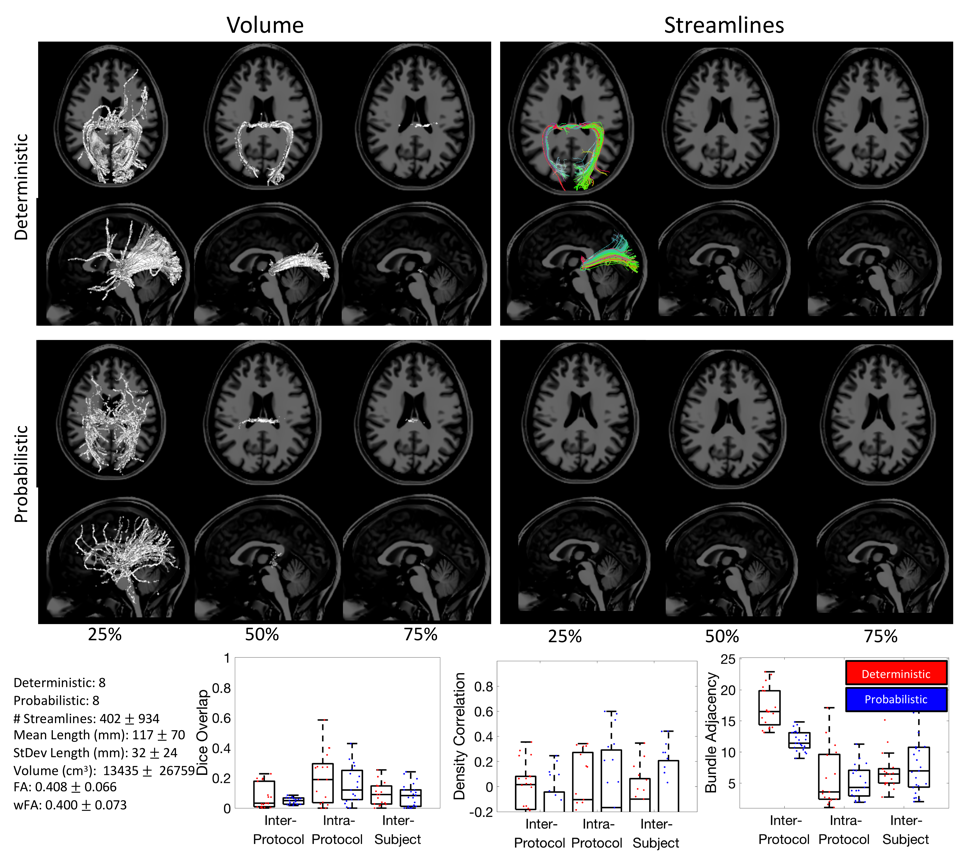

**Supplementary Figure 10.** PC Inter-protocol variability.

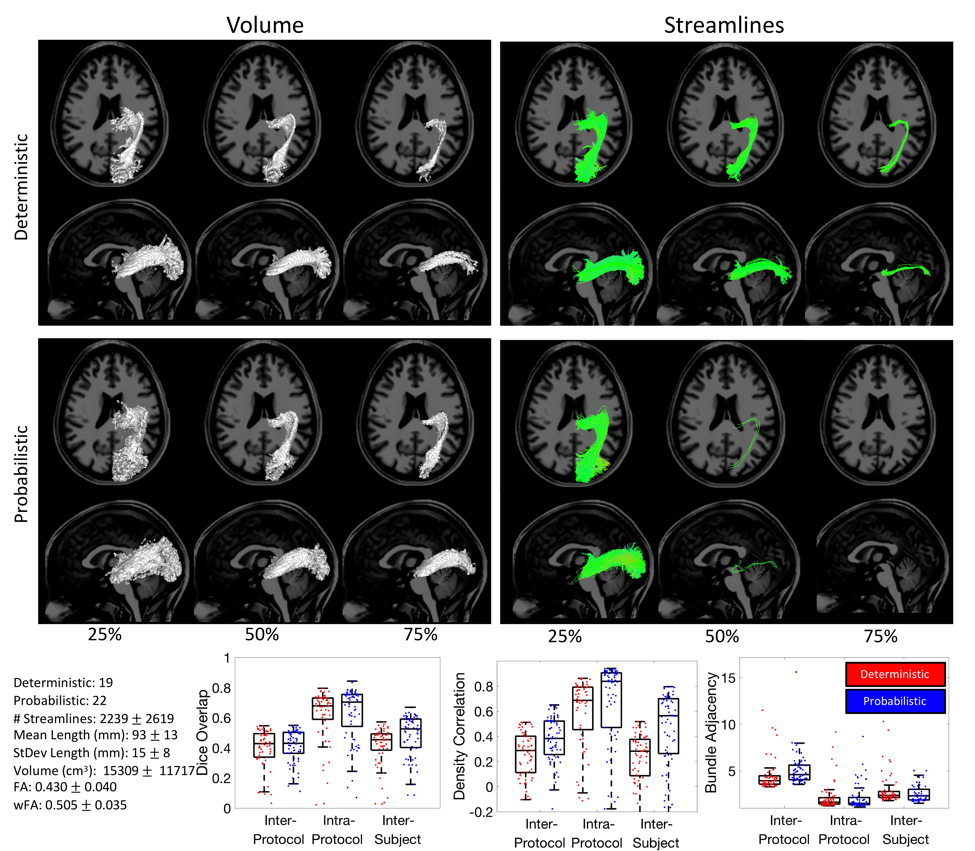

**Supplementary Figure 11.** OR Inter-protocol variability.

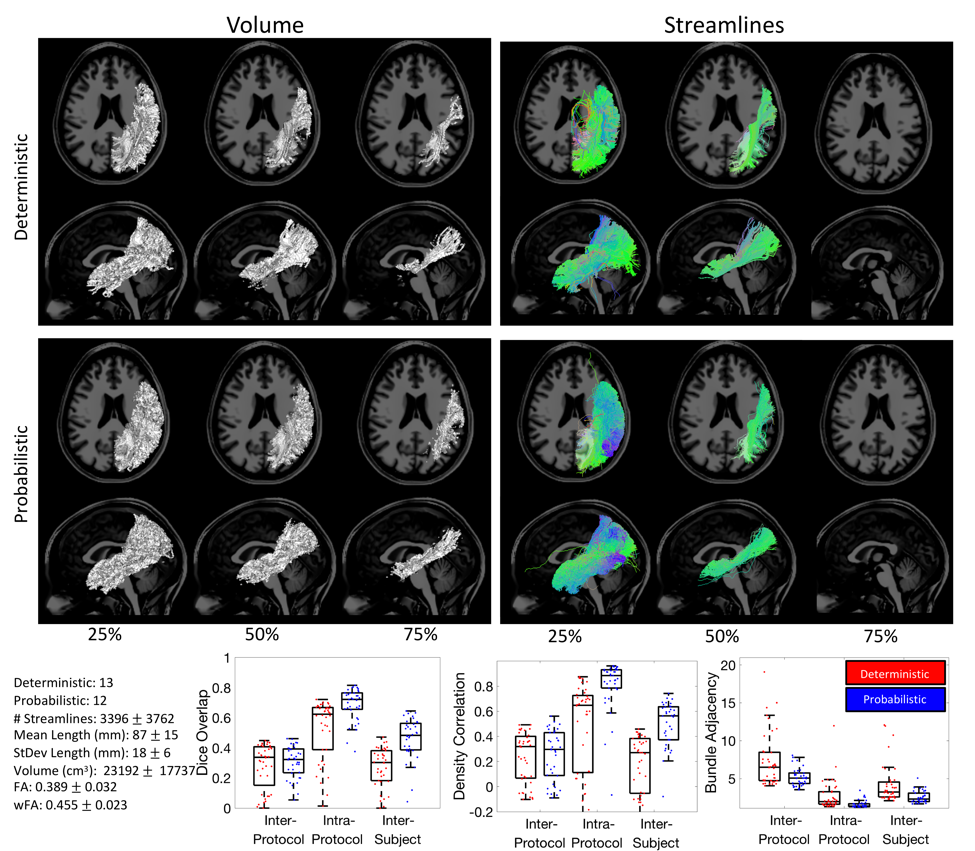

**Supplementary Figure 12.** MdLF Inter-protocol variability.

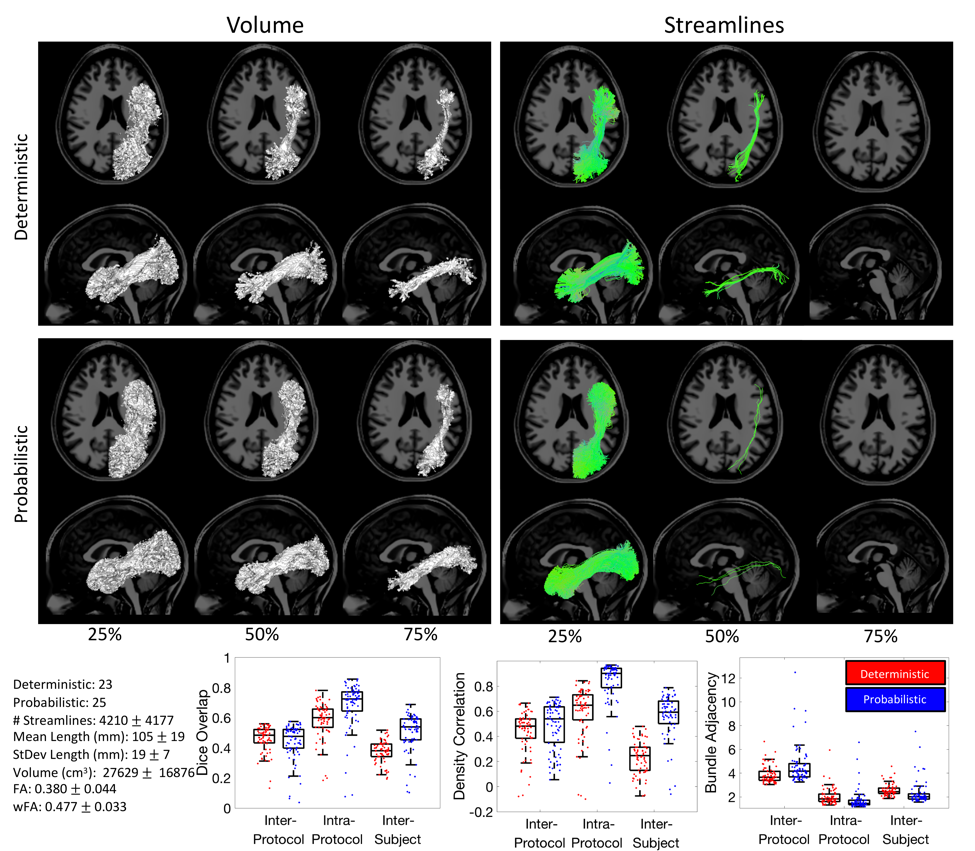

**Supplementary Figure 13.** ILF Inter-protocol variability.

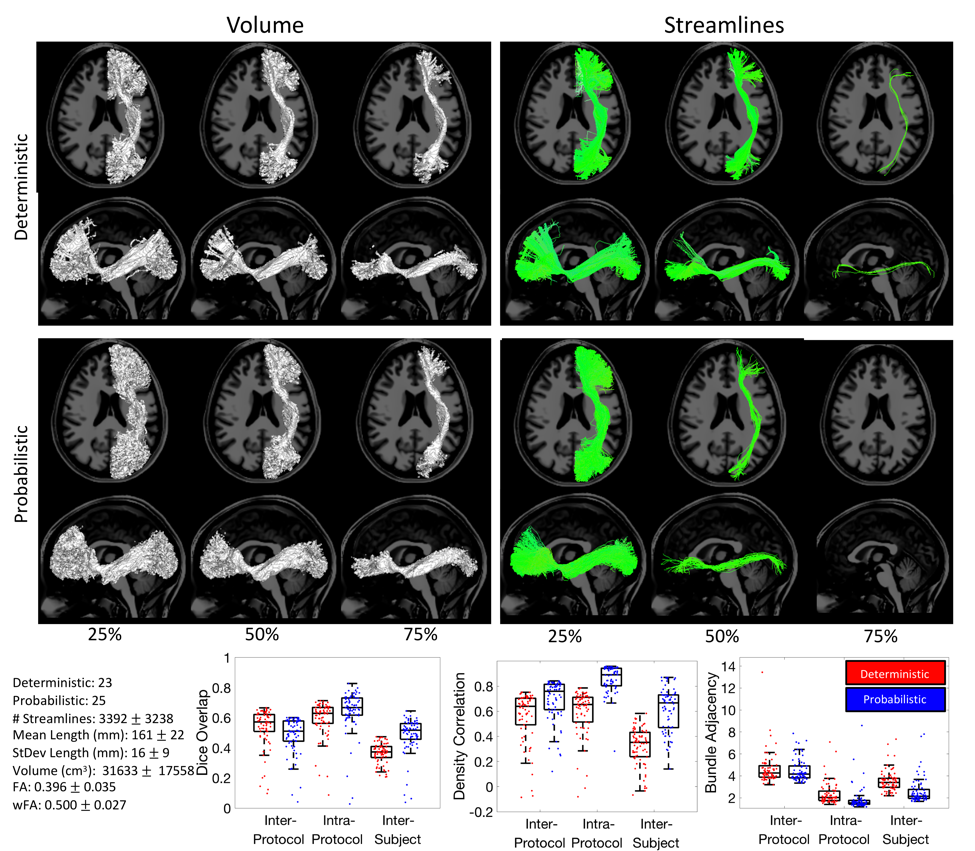

**Supplementary Figure 14.** IFOF Inter-protocol variability.

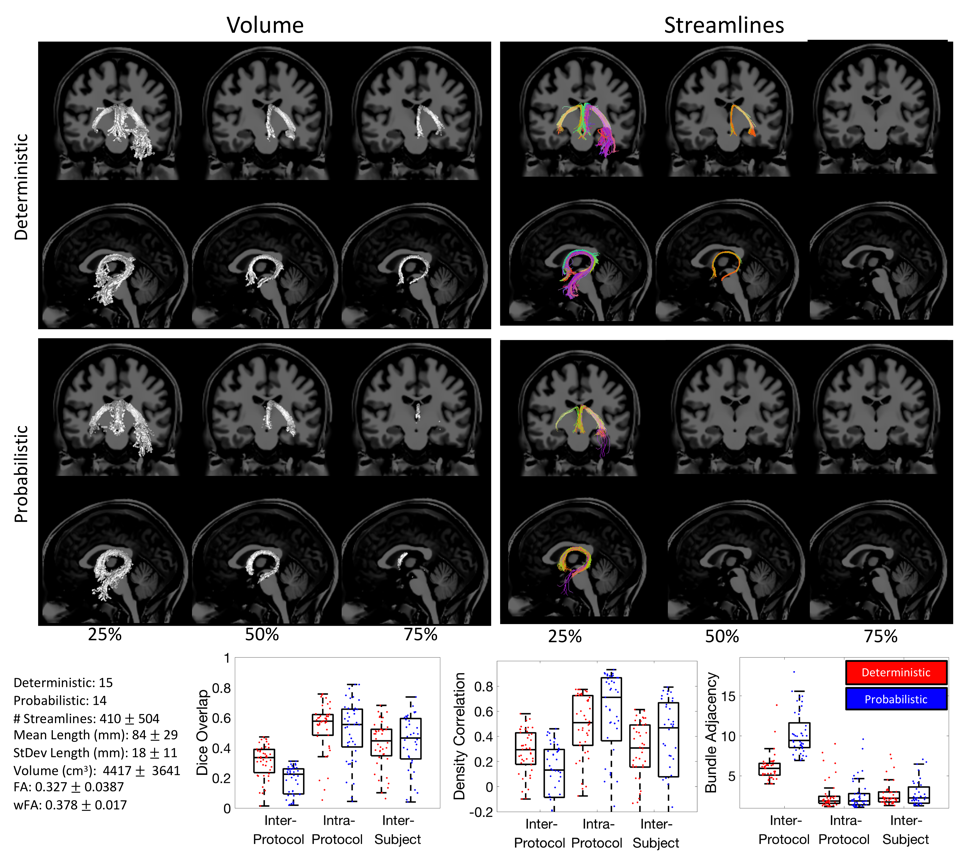

**Supplementary Figure 15.** FX Inter-protocol variability.

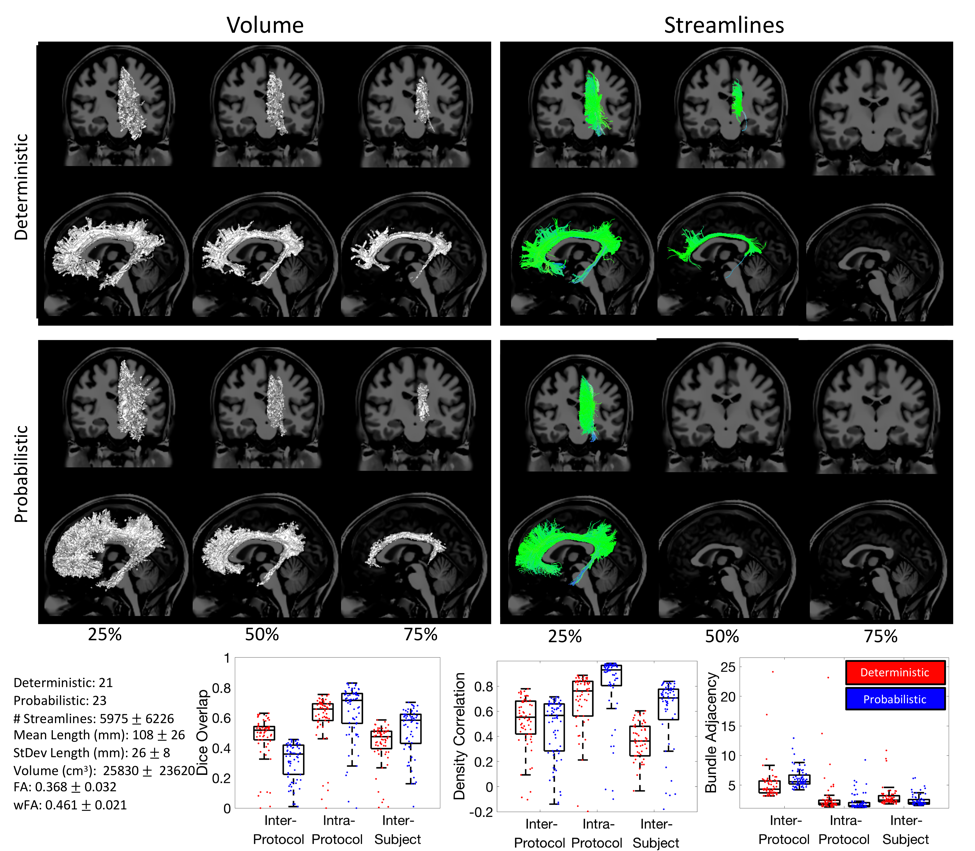

**Supplementary Figure 16.** CG Inter-protocol variability.

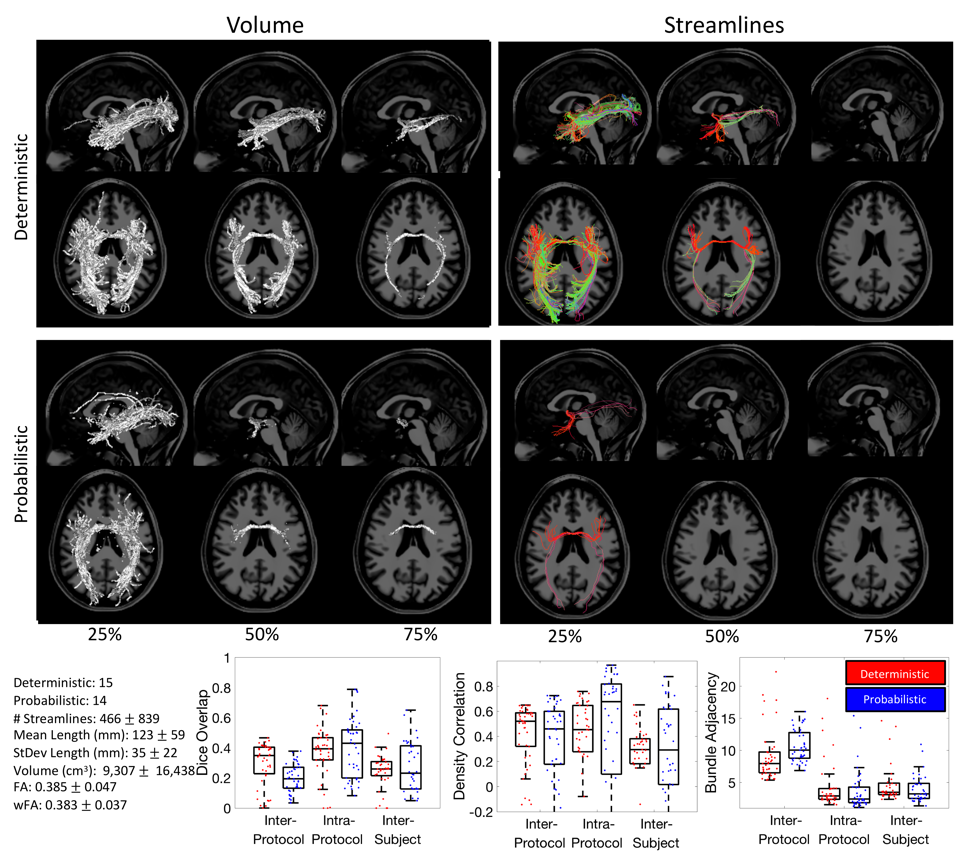

**Supplementary Figure 17.** AC Inter-protocol variability.
