## Supplementary material for "Tractography dissection variability: what happens when 42 groups dissect 14 white matter bundles on the same dataset?": QA 1 of 2

**what happens when 42 groups**

Supplementary Information

Abbreviations: Superior Longitudinal Fasciculus (SLF), Arcuate Fasciculus (AF), Optic Radiation (OR), Corticospinal Tract (CST), Cingulum (CG), Uncinate Fasciculus (UF), Corpus Callosum (CC), Middle Longitudinal Fasciculus (MdLF), Inferior Fronto-Occipital Fasciculus (IFOF), Inferior Longitudinal Fasciculus (ILF), Fornix (FX), Anterior Commissure (AC), Posterior Commissure (PC), and Parieto-Occipital Pontine Tract (POPT).

### Quality Assurance

The following figures show all streamline submissions on a single subject, for all pathways. Quality assurance was performed to ensure that (1) streamlines existed in the tractograms, (2) streamlines had correct header information (thus were in the correct subject-space), and (3) no obvious mis-labelling was performed. We note that we did not attempt to classify any pathways as “wrong”, and accepted tractograms as a valid representation of the intended white matter bundle as long as they passed all criteria above.

We note that this is only a single subject, and analysis in the study is based on a total of six subjects (thus, we show only 1/6 the total number of tractograms below).

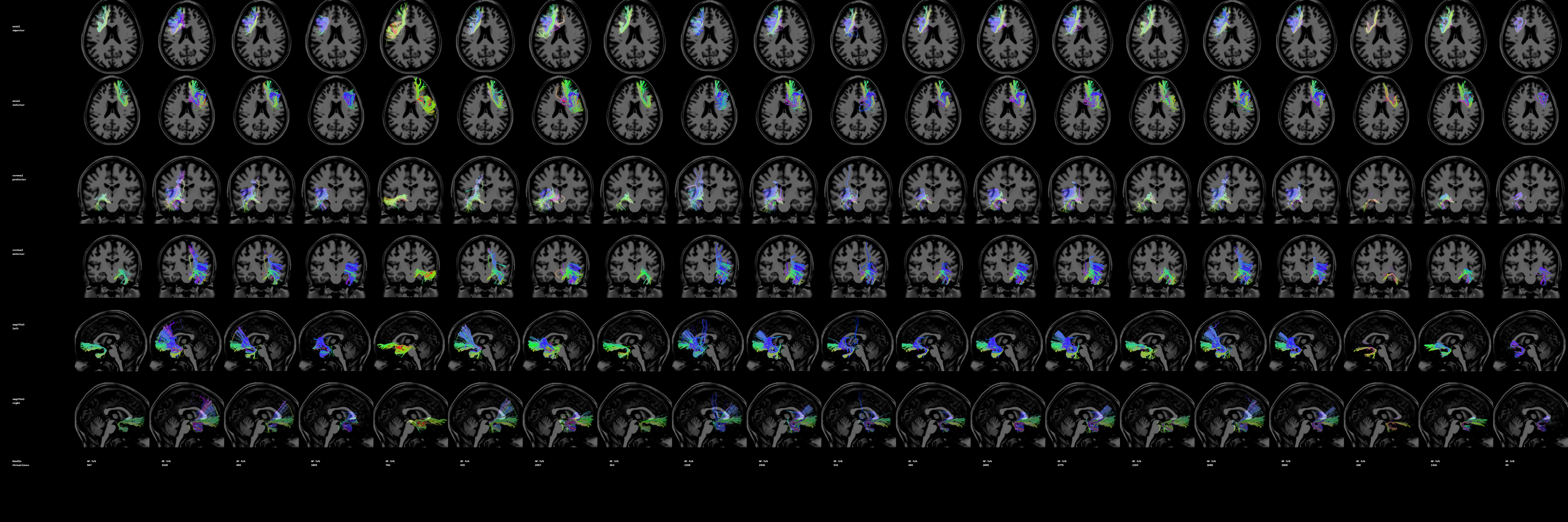
**Supplementary Figure 18.** UF deterministic submissions for a single subject.

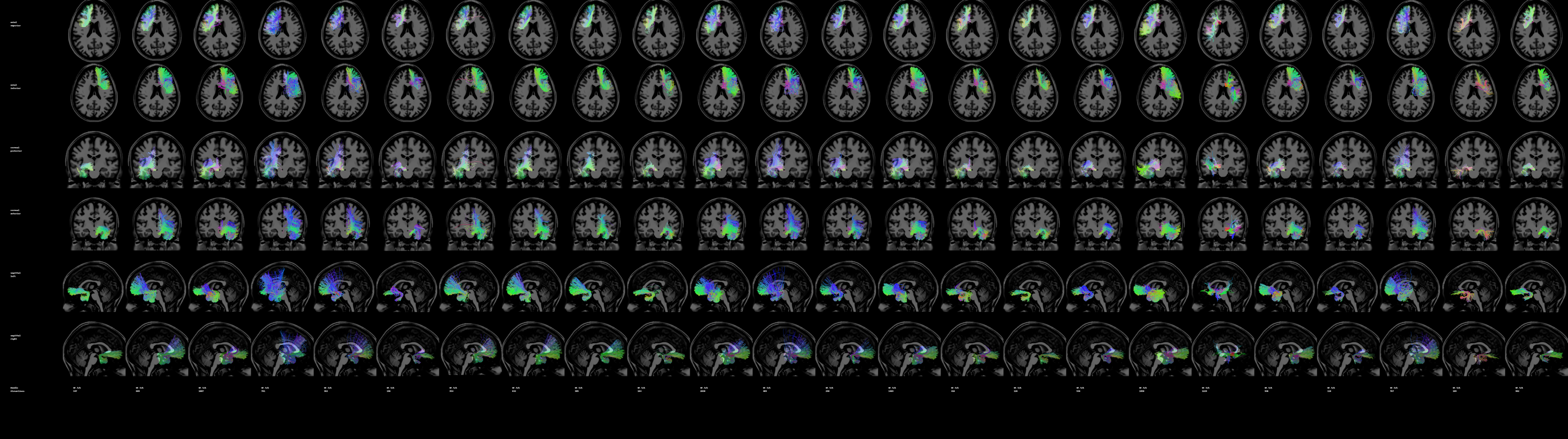
 **Supplementary Figure 19.** UF probabilistic submissions for a single subject.

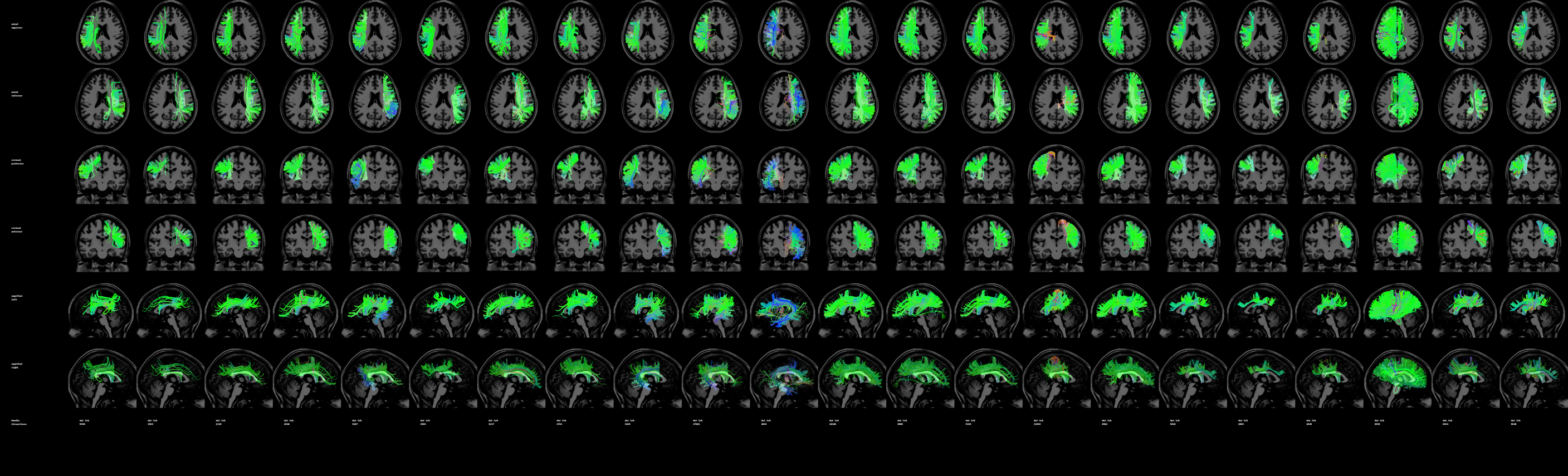

**Supplementary Figure 20.** SLF deterministic submissions for a single subject.

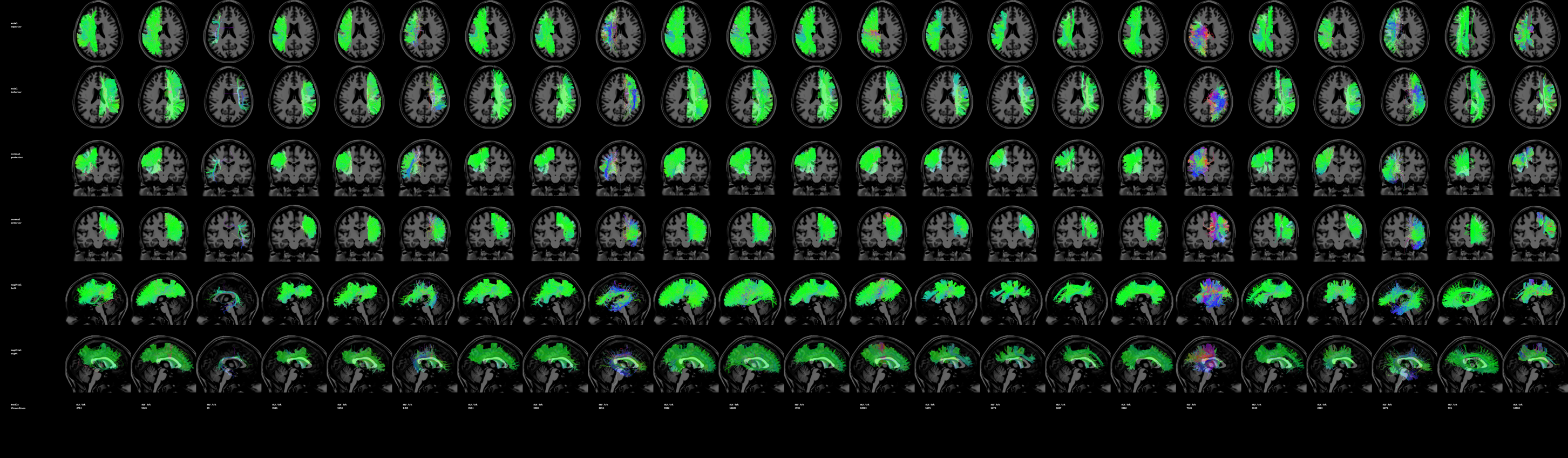

**Supplementary Figure 21.** SLF probabilistic submissions for a single subject.

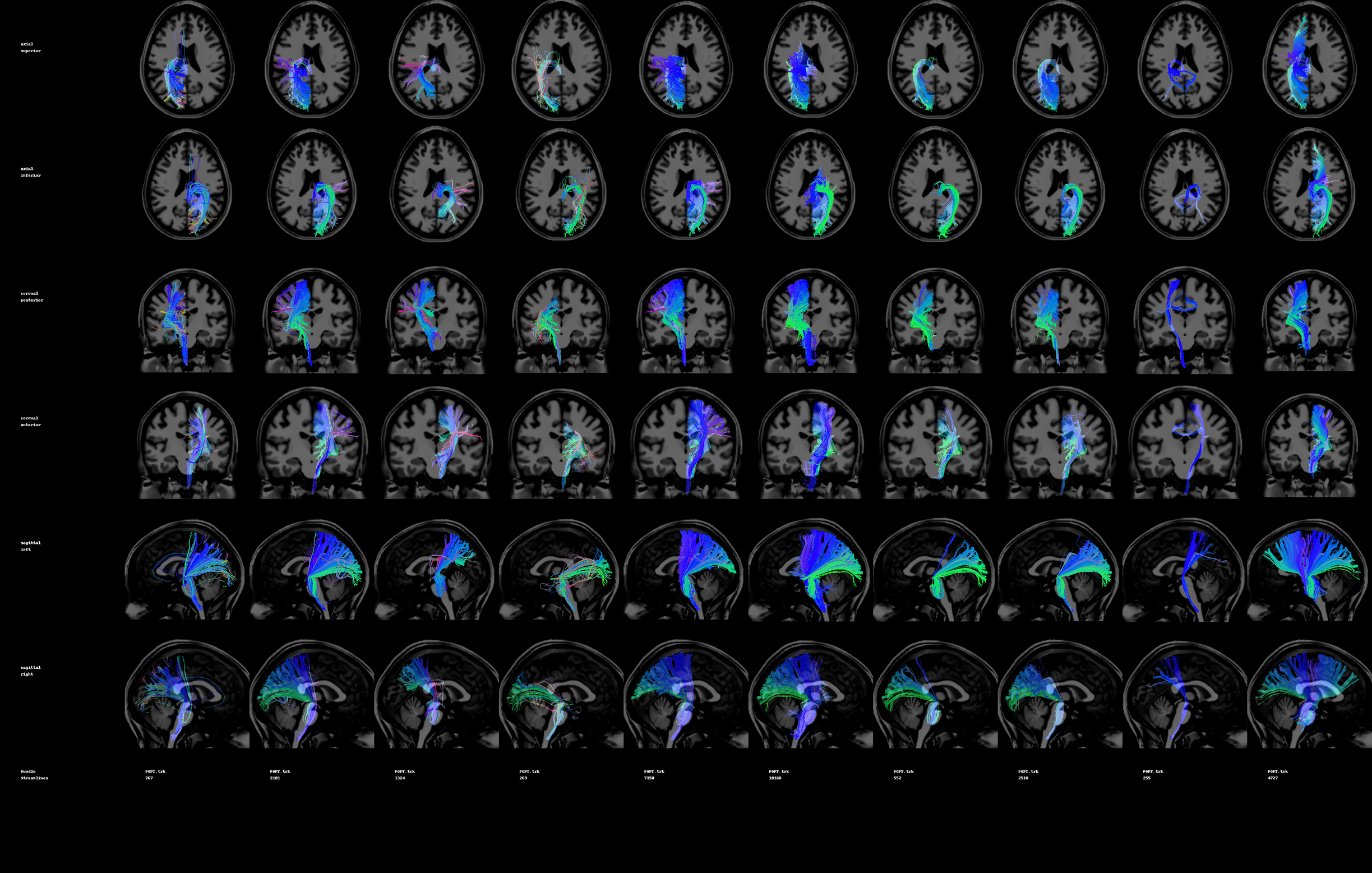

**Supplementary Figure 22.** POPT deterministic submissions for a single subject.

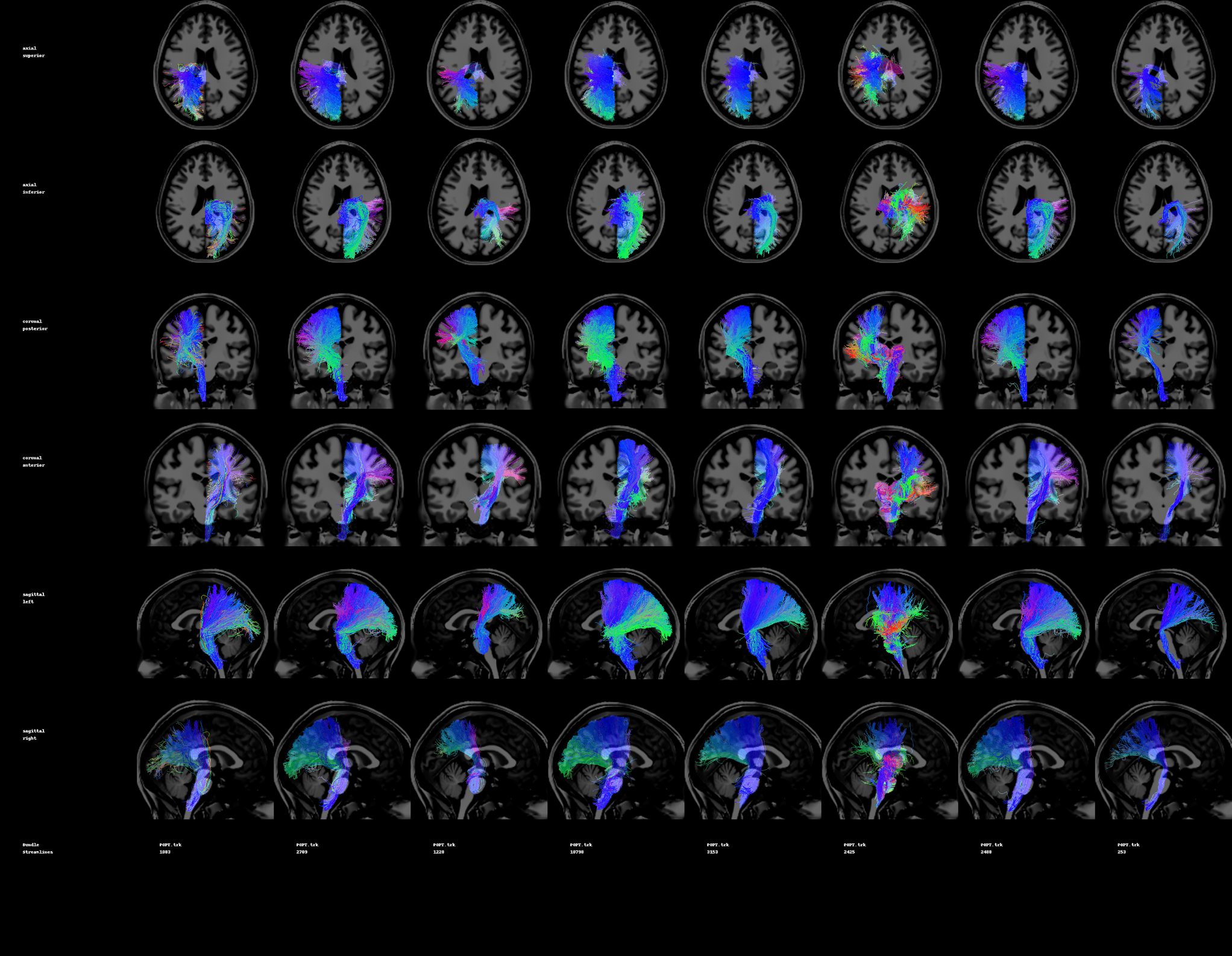

**Supplementary Figure 23.** POPT probabilistic submissions for a single subject.

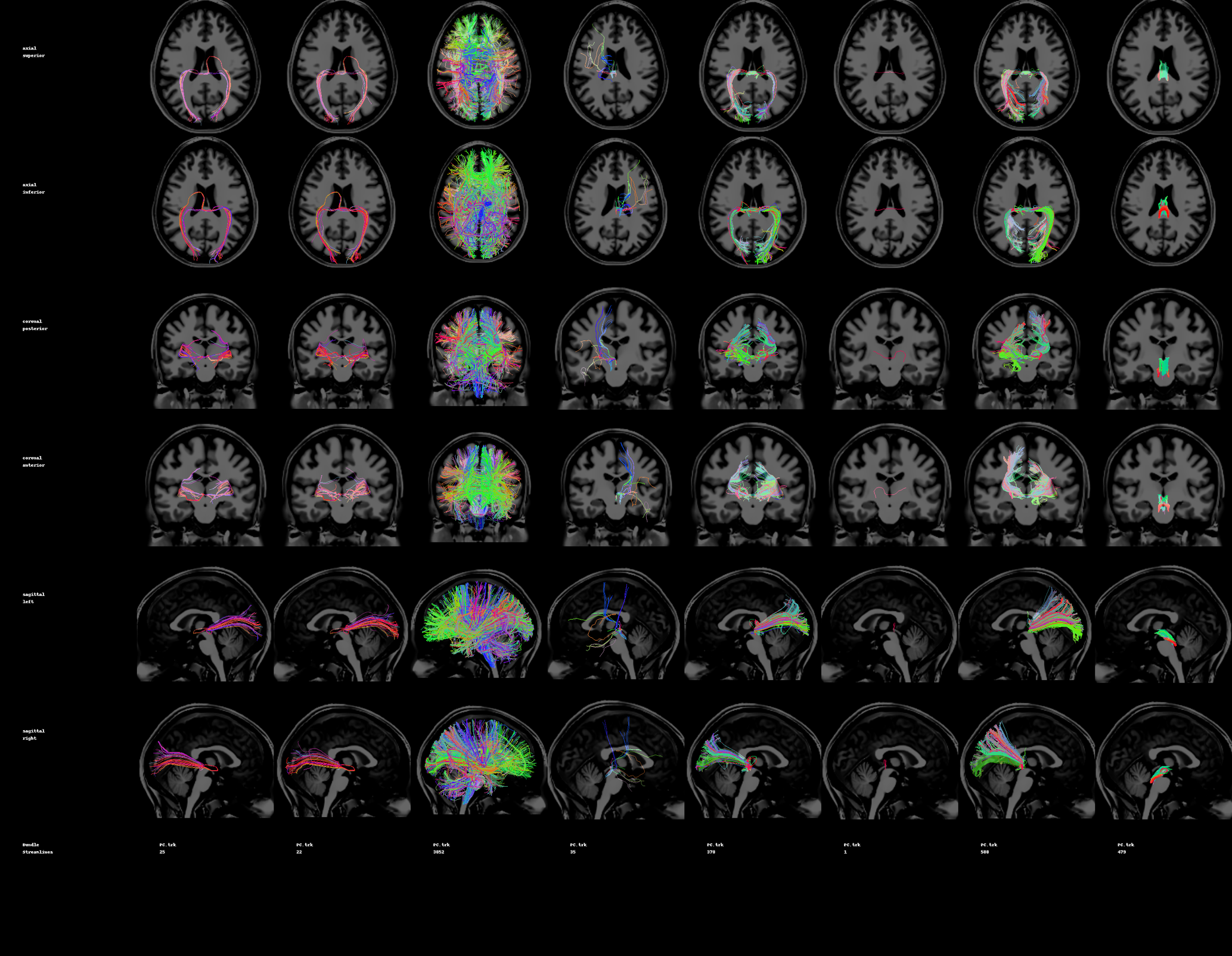

**Supplementary Figure 24.** PC deterministic submissions for a single subject.

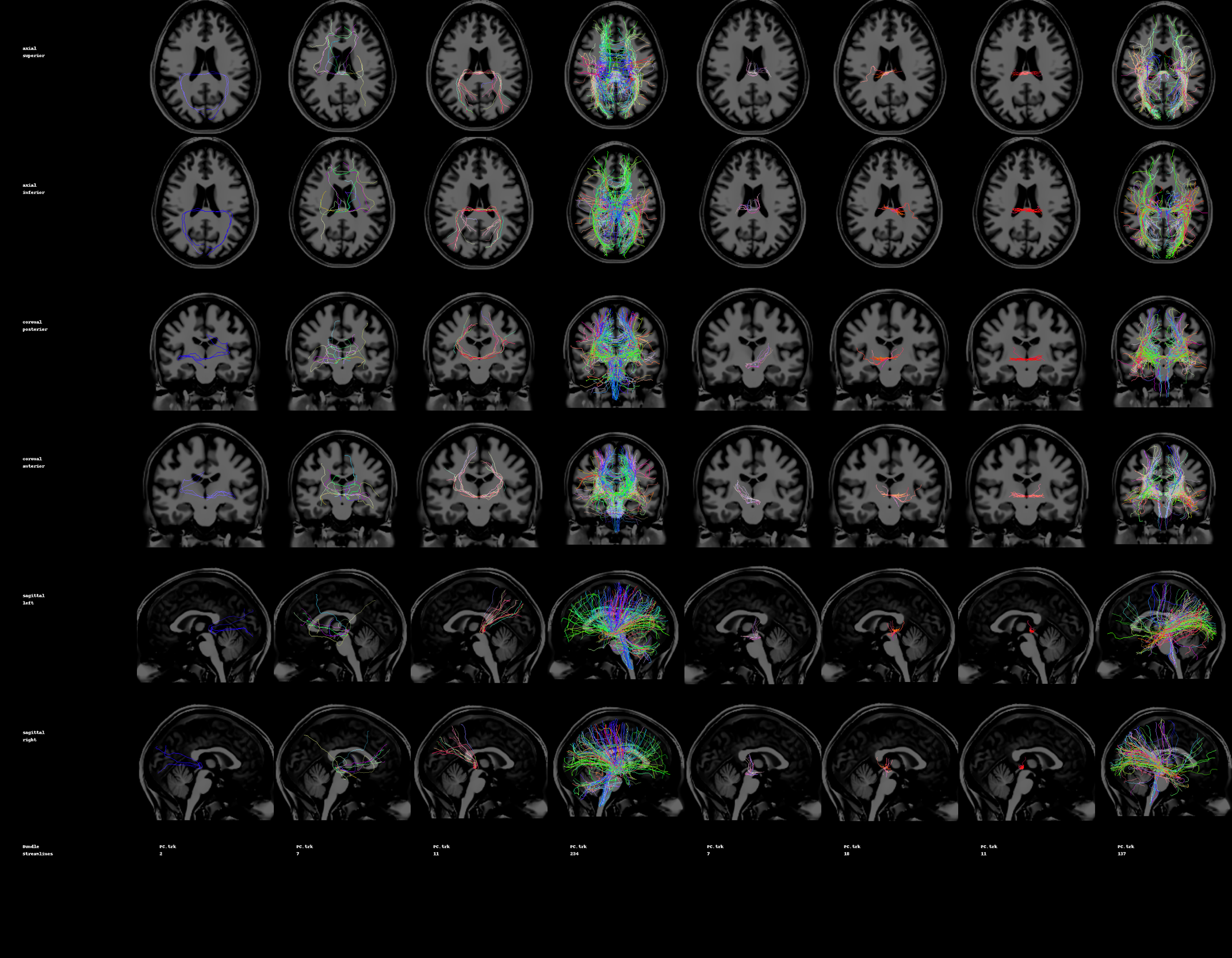

**Supplementary Figure 25.** PC probabilistic submissions for a single subject.

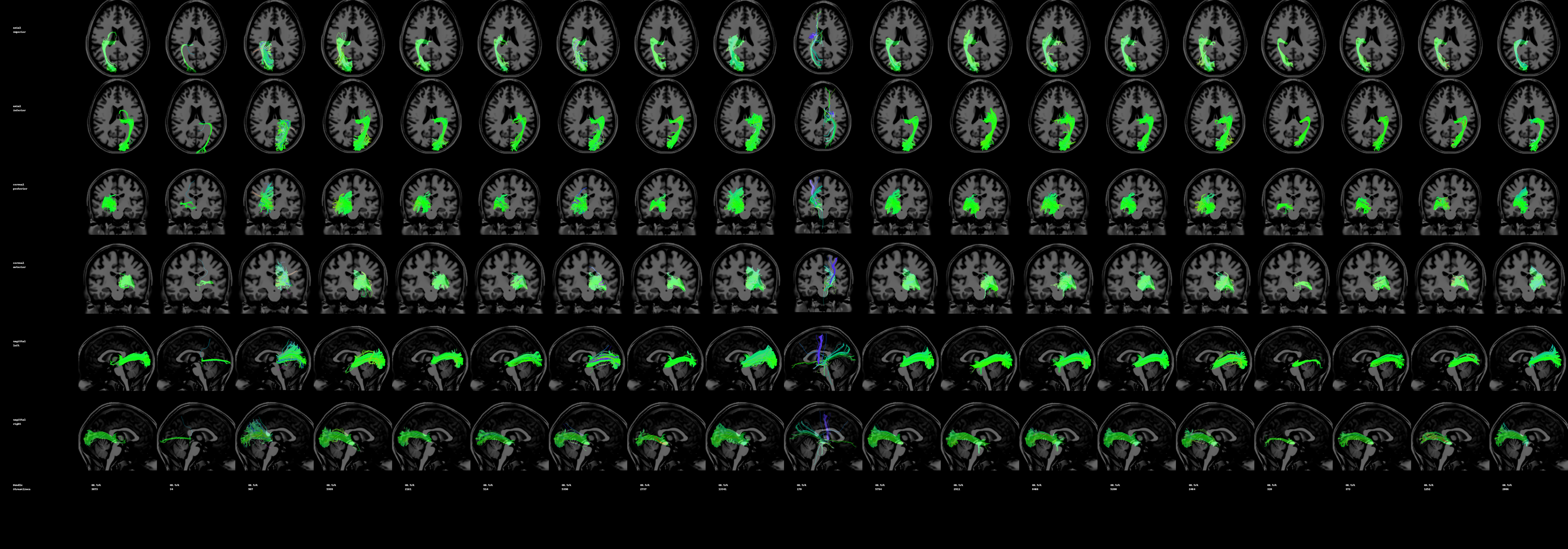

**Supplementary Figure 26.** OR deterministic submissions for a single subject.

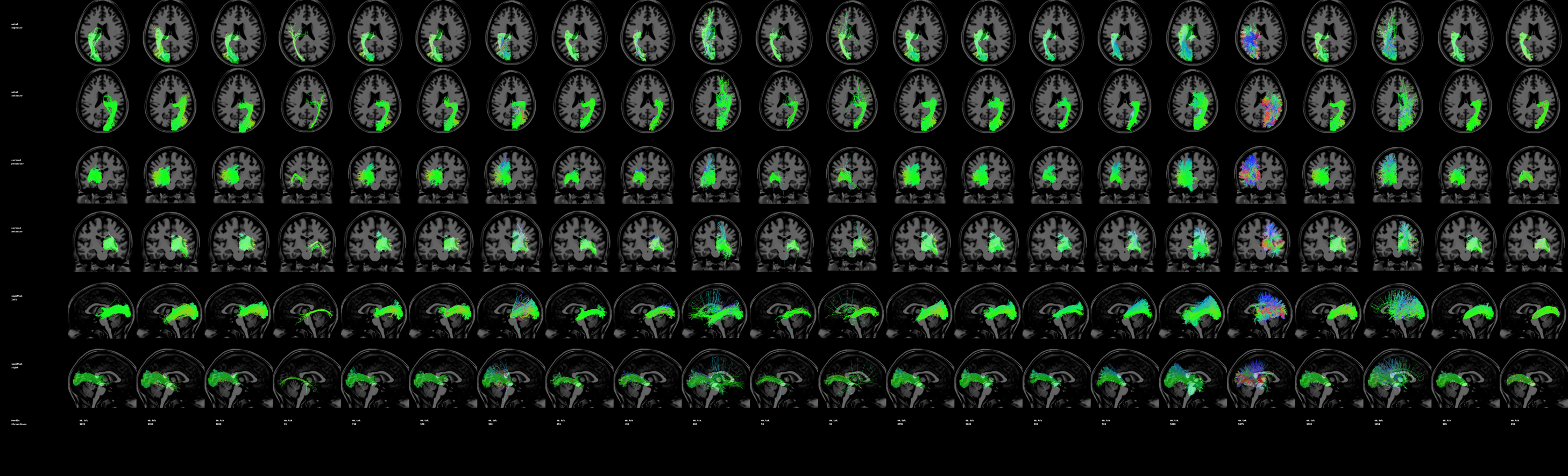

**Supplementary Figure 27.** OR probabilistic submissions for a single subject.

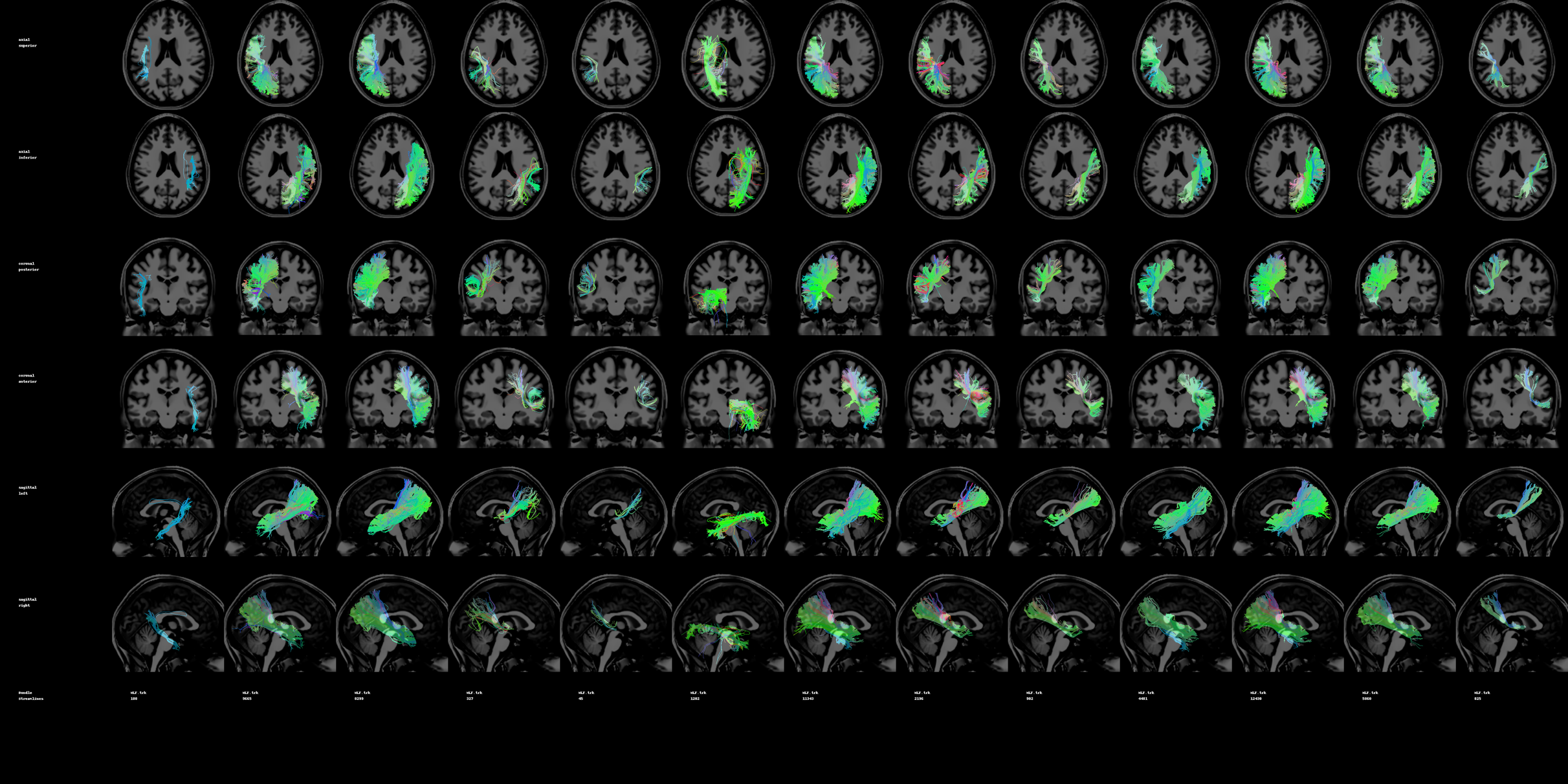

**Supplementary Figure 28.** MdLF deterministic submissions for a single subject.

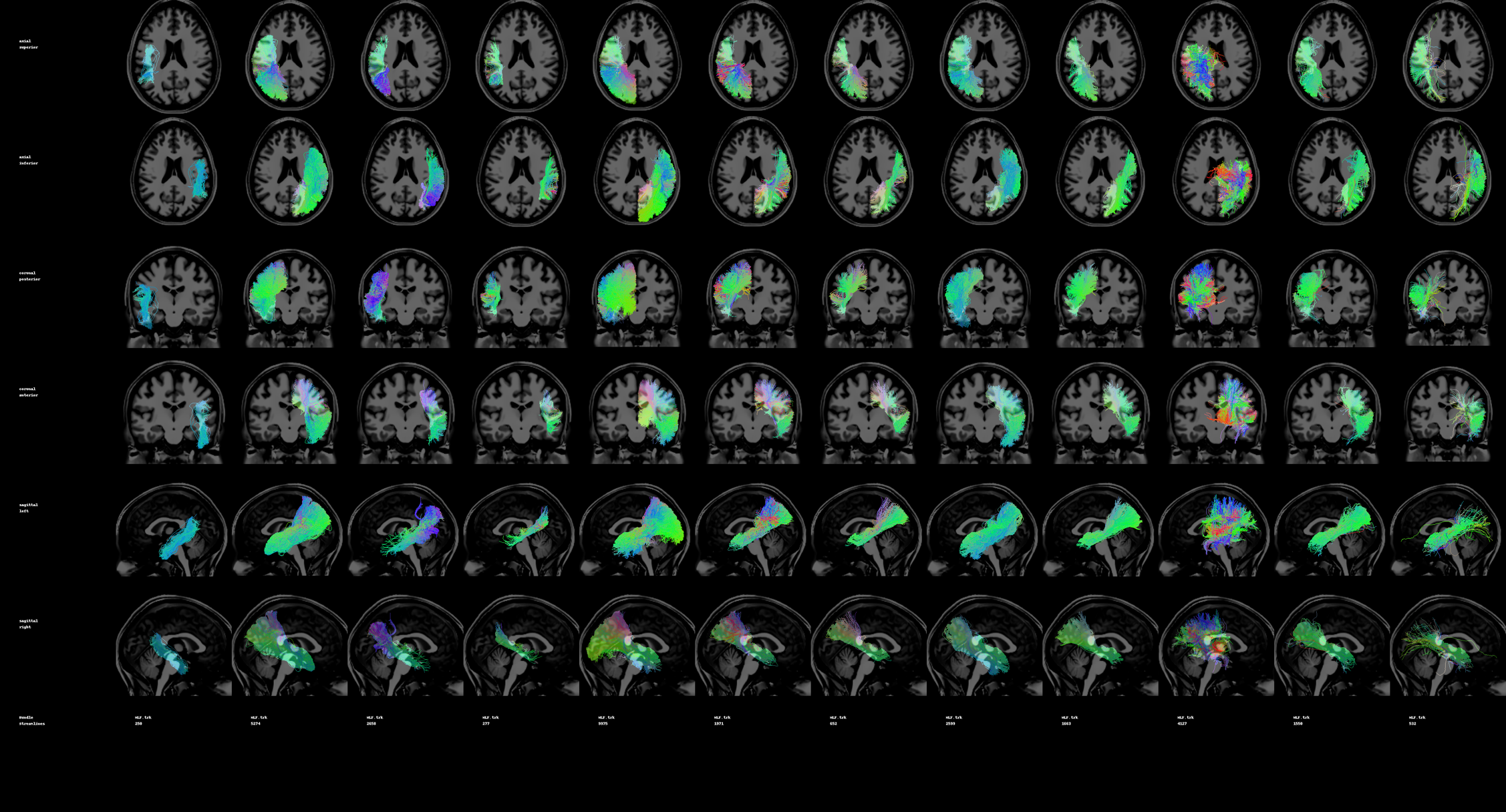

**Supplementary Figure 29.** MdLF probabilistic submissions for a single subject.

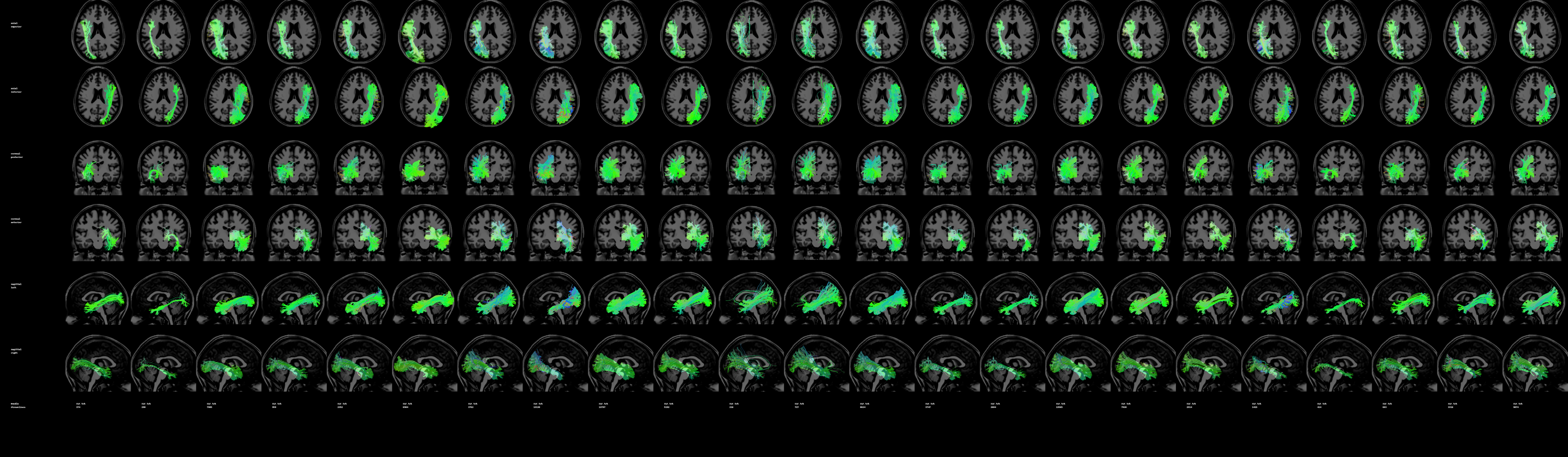

**Supplementary Figure 30.** ILF deterministic submissions for a single subject.

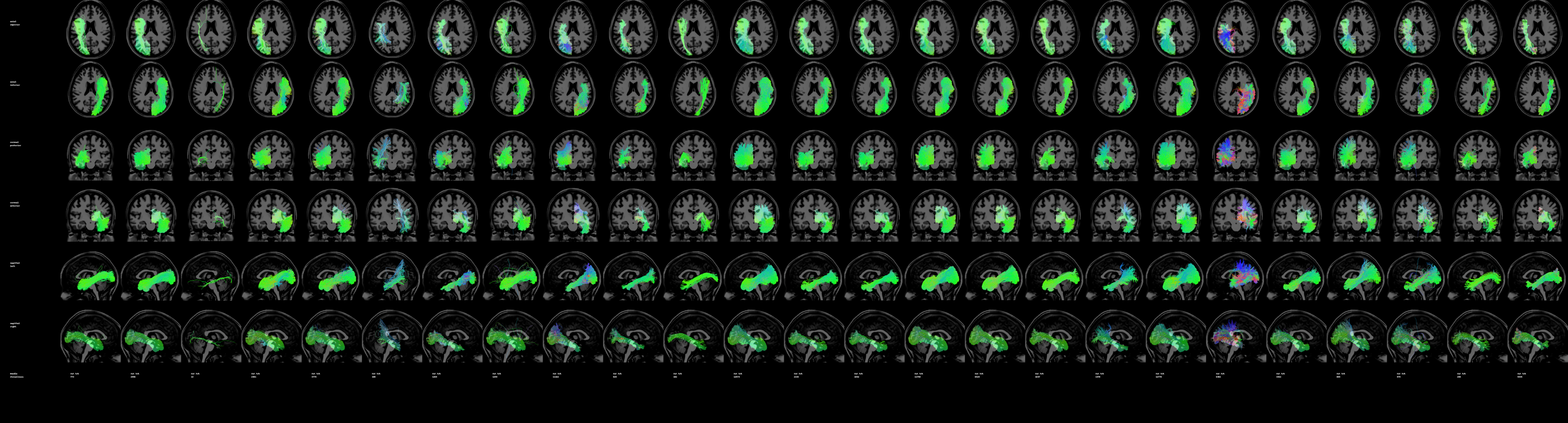

**Supplementary Figure 31.** ILF probabilistic submissions for a single subject
