## Supplementary material for "Tractography dissection variability: what happens when 42 groups dissect 14 white matter bundles on the same dataset?": QA 2 of 2

**what happens when 42 groups**

Supplementary Information

Abbreviations: Superior Longitudinal Fasciculus (SLF), Arcuate Fasciculus (AF), Optic Radiation (OR), Corticospinal Tract (CST), Cingulum (CG), Uncinate Fasciculus (UF), Corpus Callosum (CC), Middle Longitudinal Fasciculus (MdLF), Inferior Fronto-Occipital Fasciculus (IFOF), Inferior Longitudinal Fasciculus (ILF), Fornix (FX), Anterior Commissure (AC), Posterior Commissure (PC), and Parieto-Occipital Pontine Tract (POPT).

### Quality Assurance

The following figures show all streamline submissions on a single subject, for all pathways. Quality assurance was performed to ensure that (1) streamlines existed in the tractograms, (2) streamlines had correct header information (thus were in the correct subject-space), and (3) no obvious mis-labelling was performed. We note that we did not attempt to classify any pathways as “wrong”, and accepted tractograms as a valid representation of the intended white matter bundle as long as they passed all criteria above.

We note that this is only a single subject, and analysis in the study is based on a total of six subjects (thus, we show only 1/6 the total number of tractograms below).

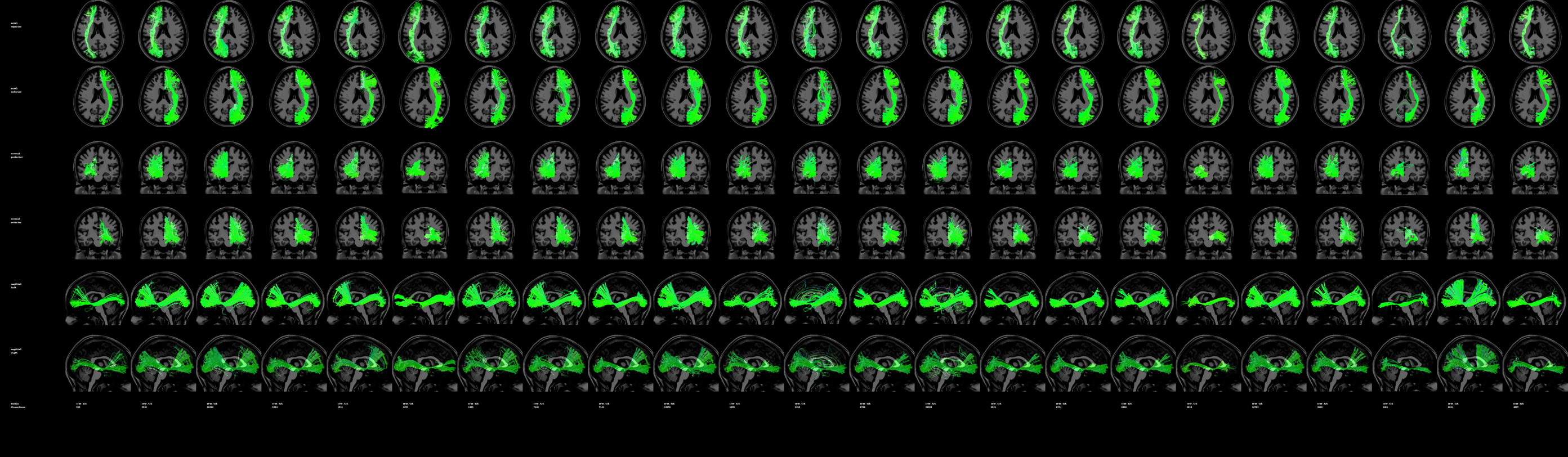

**Supplementary Figure 32.** IFOF deterministic submissions for a single subject.

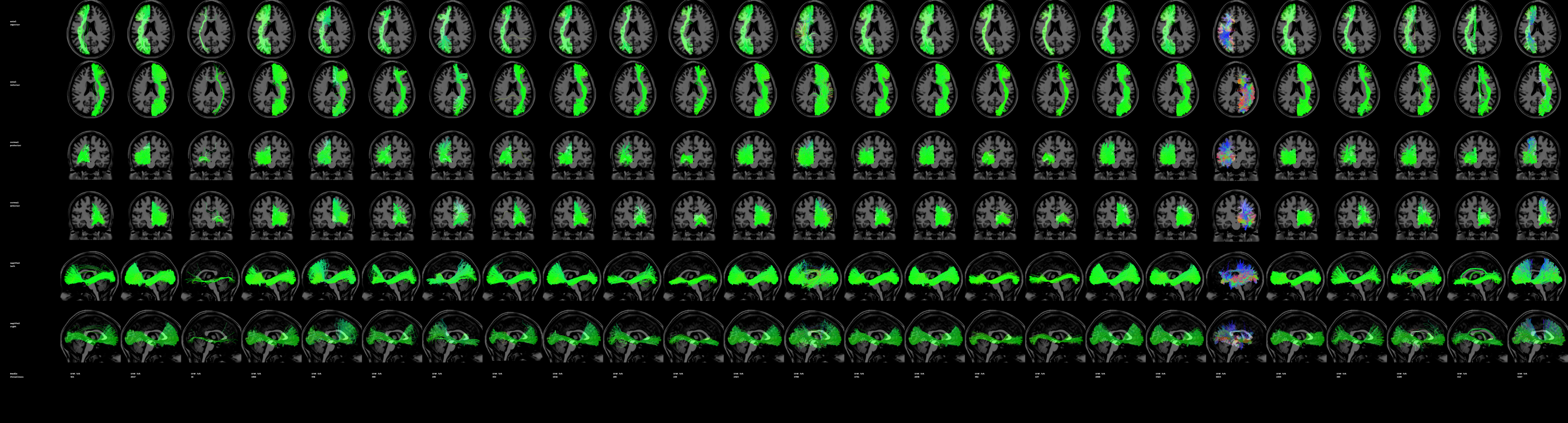

**Supplementary Figure 33.** IFOF probabilistic submissions for a single subject.

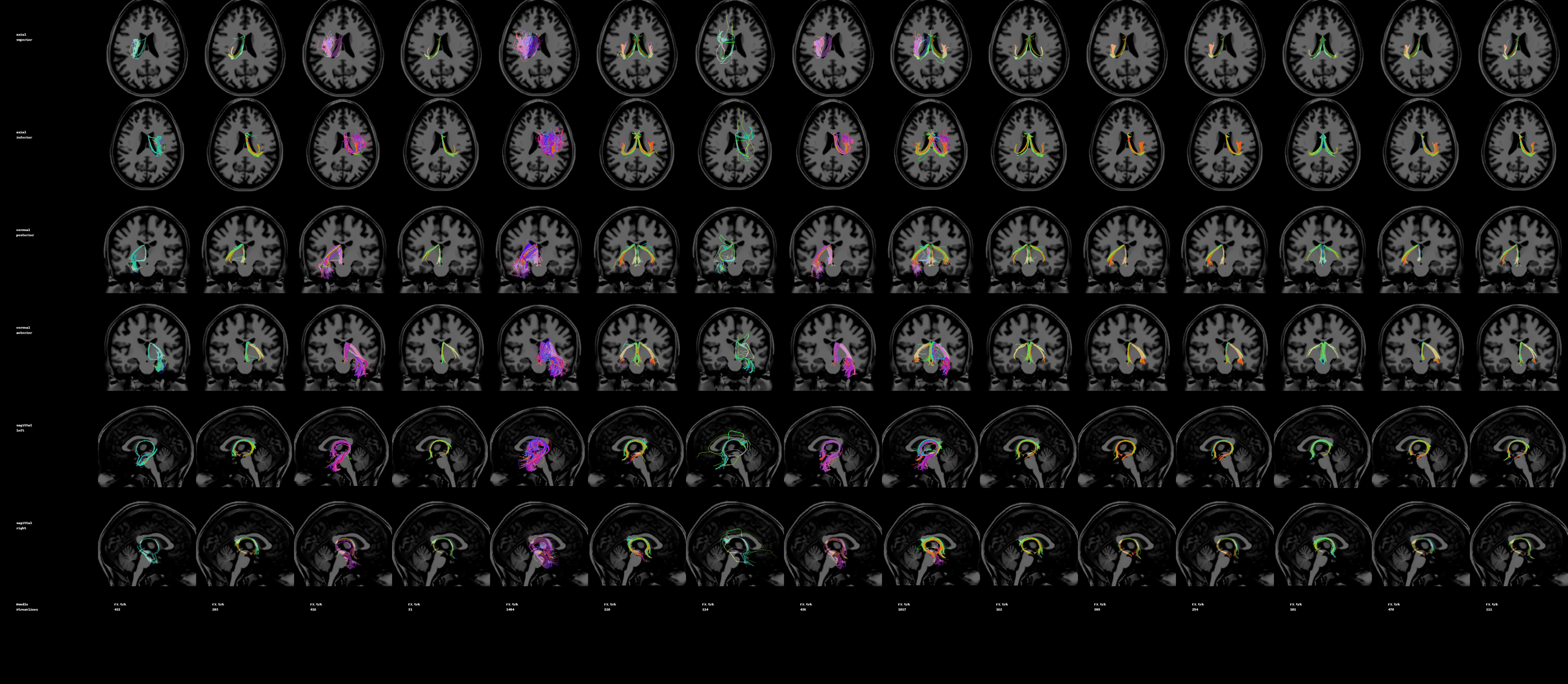

**Supplementary Figure 34.** FX deterministic submissions for a single subject.

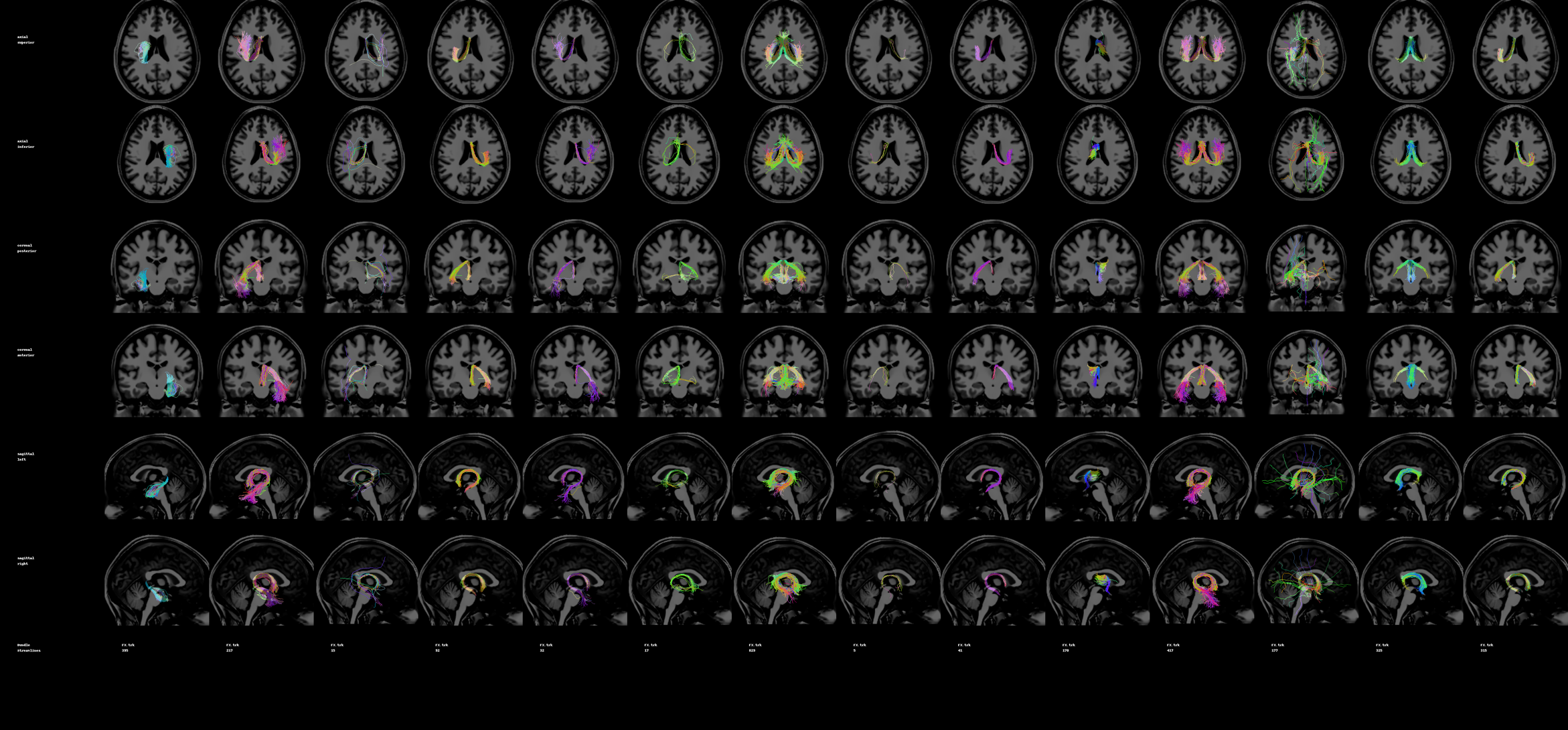

**Supplementary Figure 35.** FX probabilistic submissions for a single subject.

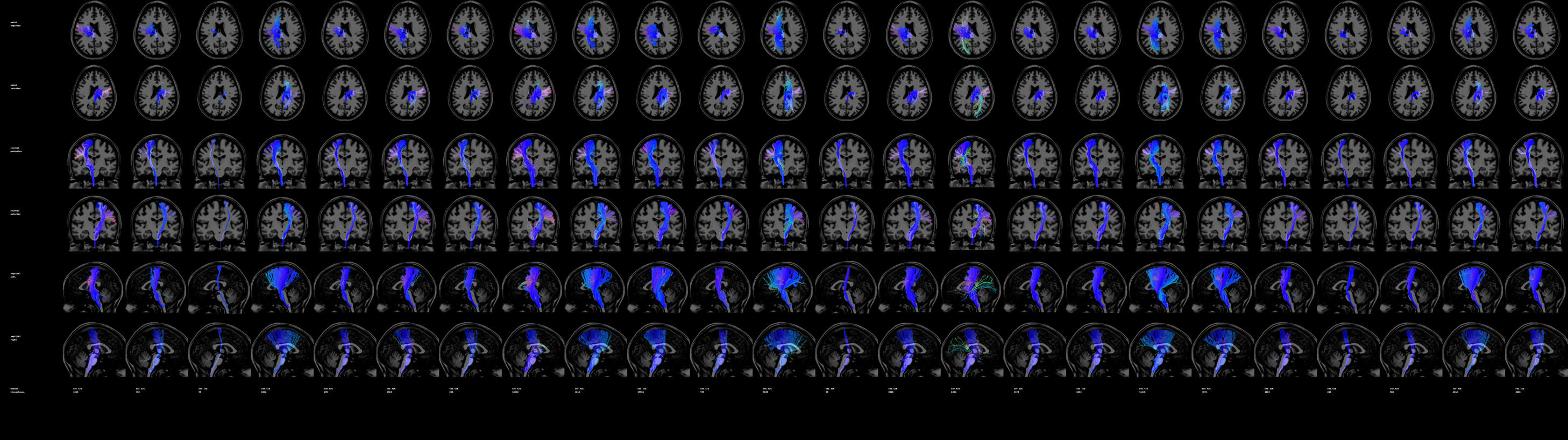

**Supplementary Figure 36.** CST deterministic submissions for a single subject.

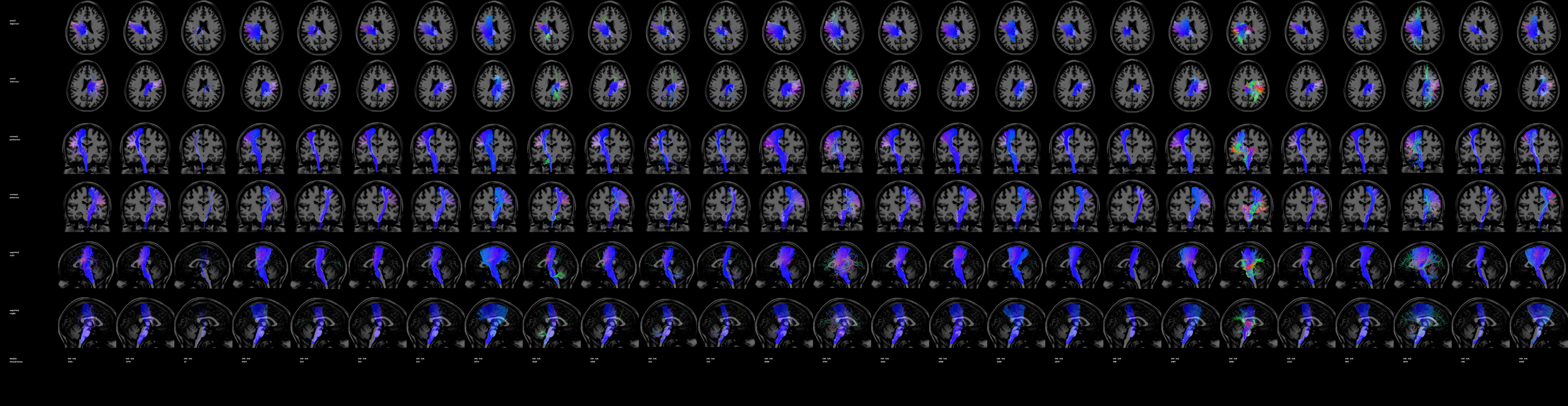

**Supplementary Figure 37.** CST probabilistic submissions for a single subject.

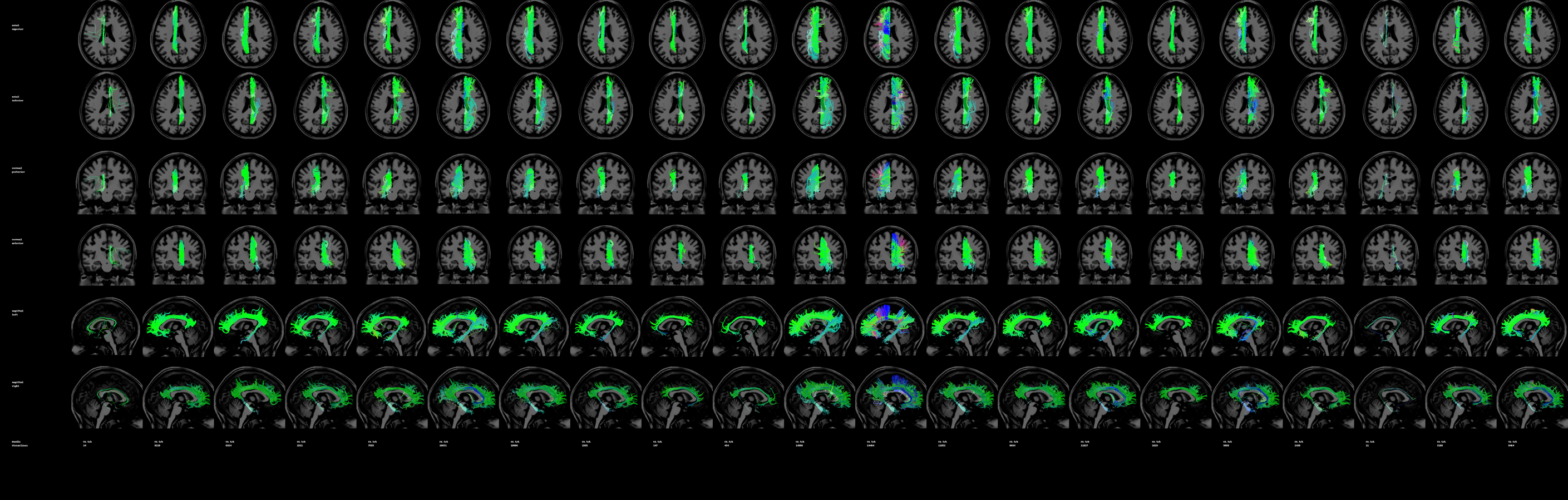

**Supplementary Figure 38.** CG deterministic submissions for a single subject.

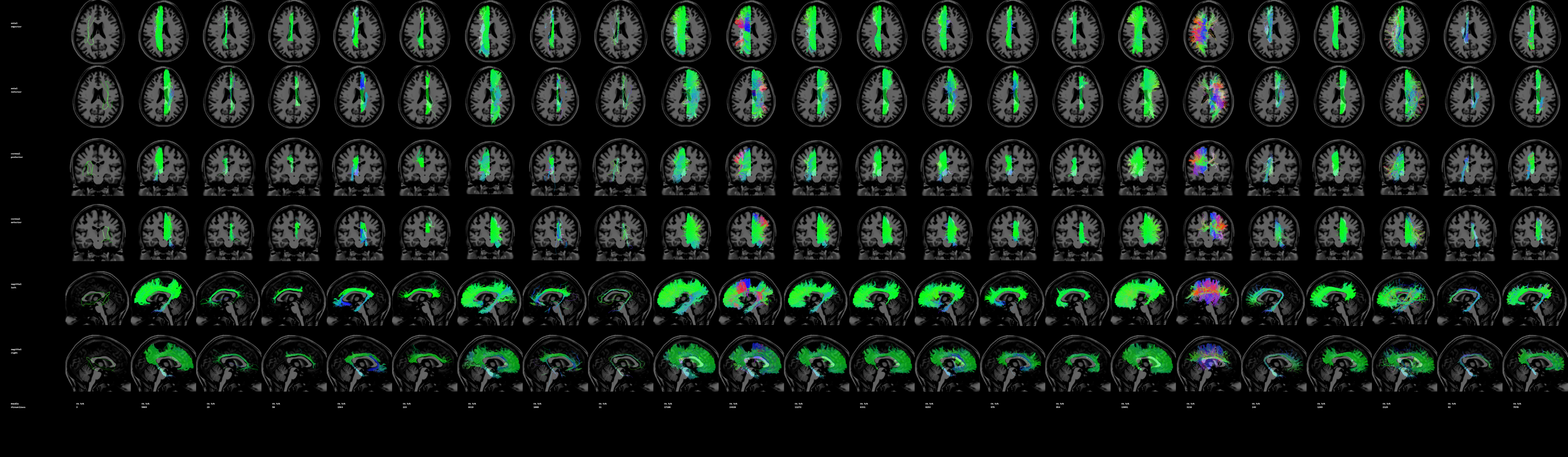

**Supplementary Figure 39.** CG probabilistic submissions for a single subject.

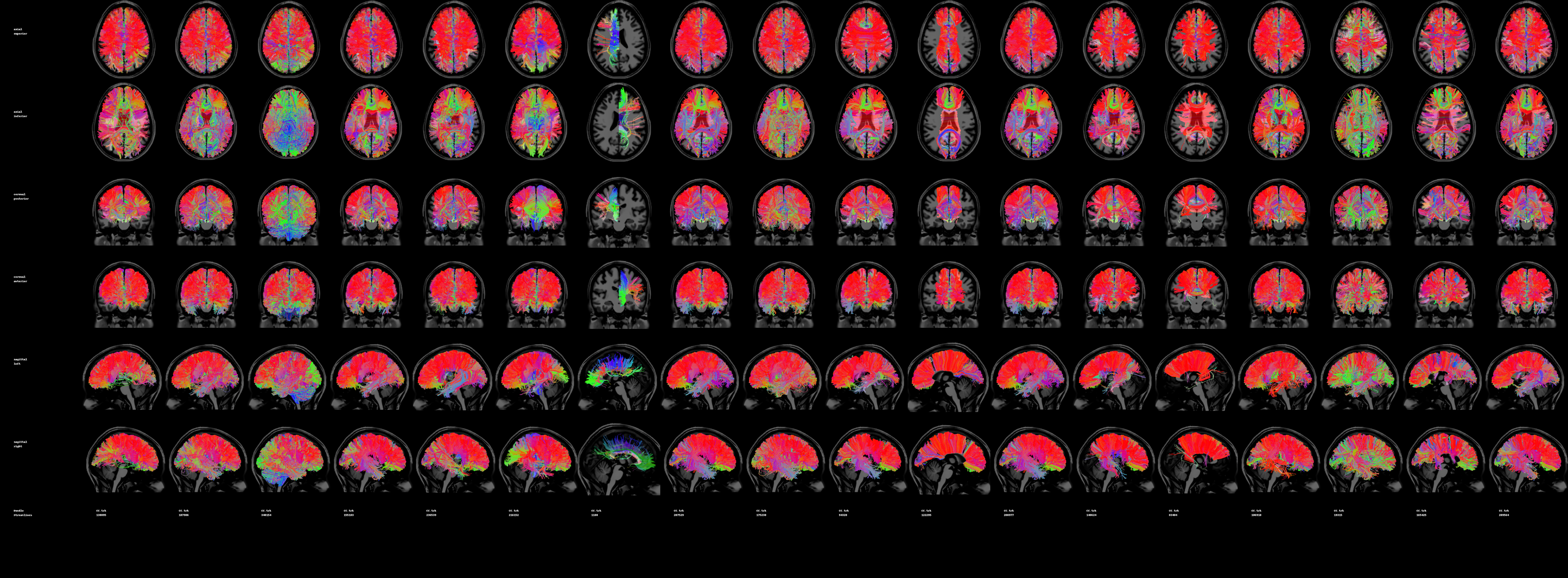

**Supplementary Figure 40.** CC deterministic submissions for a single subject.

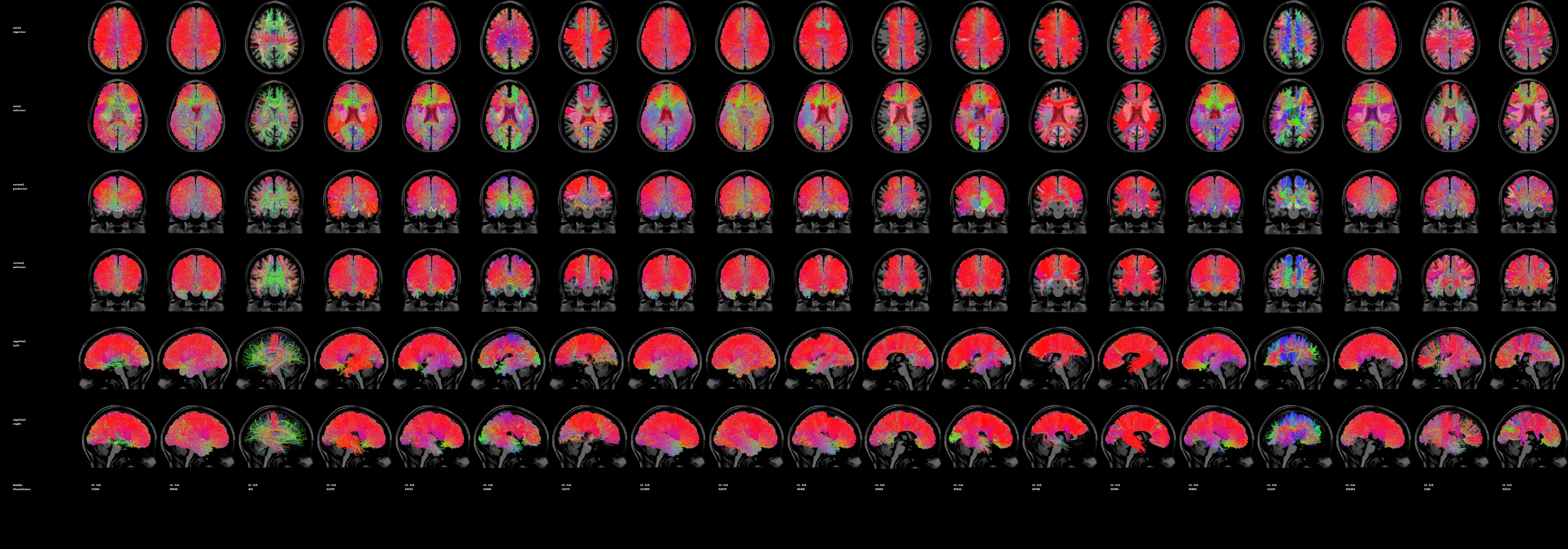

**Supplementary Figure 41.** CC probabilistic submissions for a single subject.

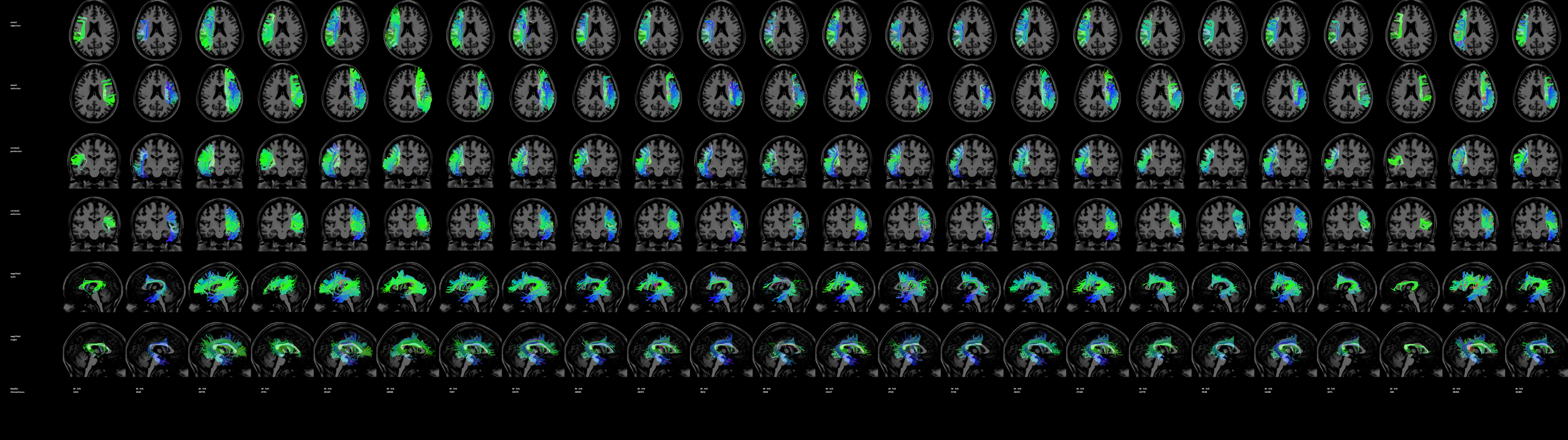

**Supplementary Figure 42.** AF deterministic submissions for a single subject.

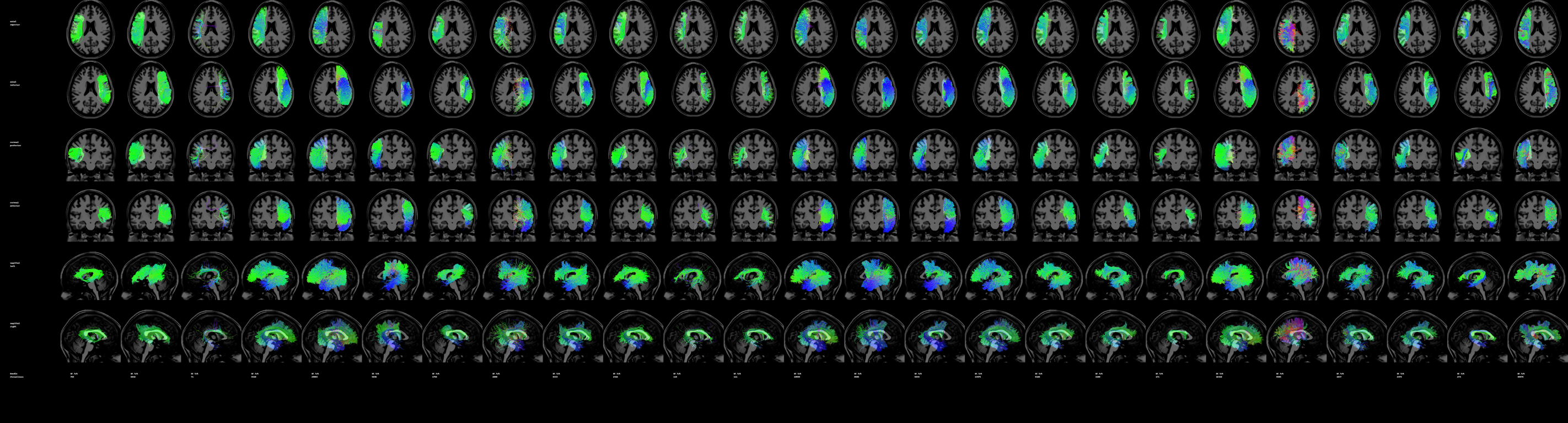

**Supplementary Figure 43.** AF probabilistic submissions for a single subject.

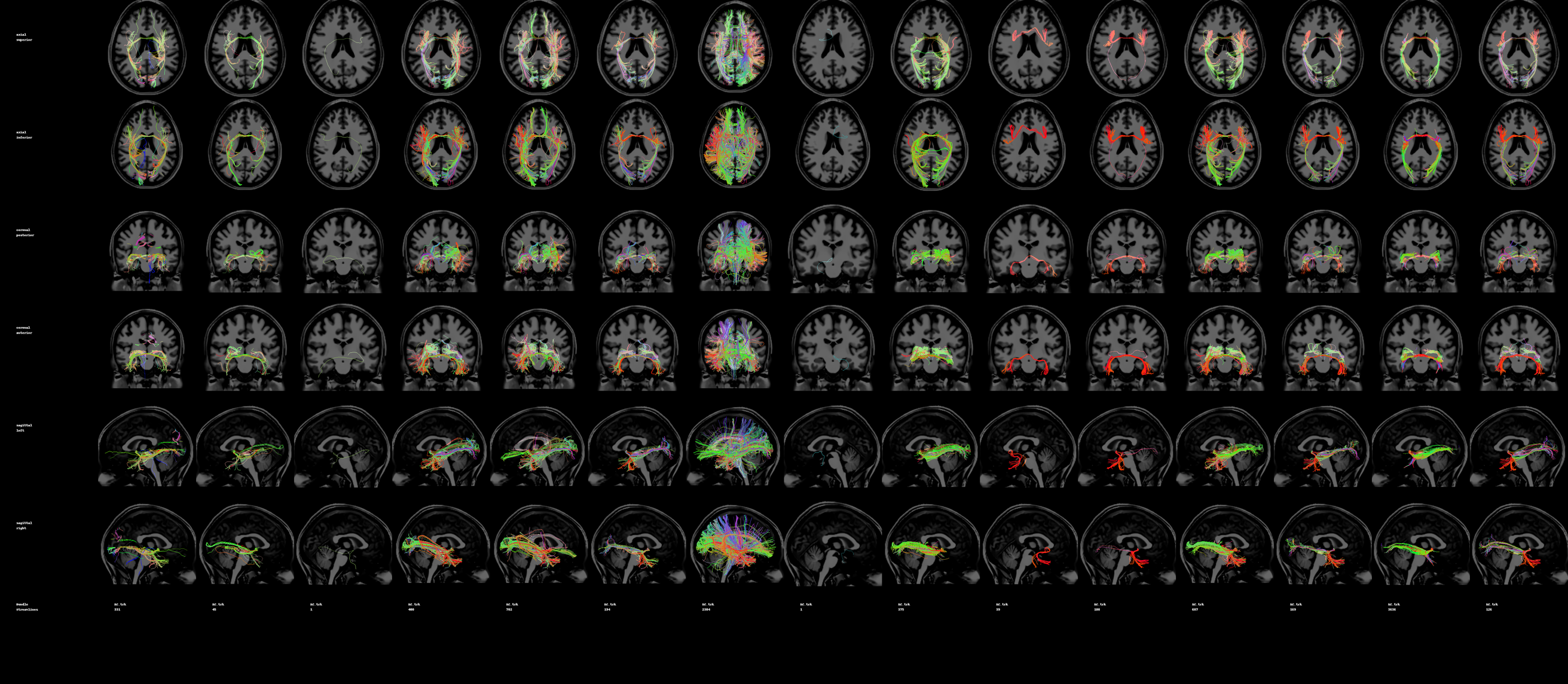

**Supplementary Figure 44.** AC deterministic submissions for a single subject.

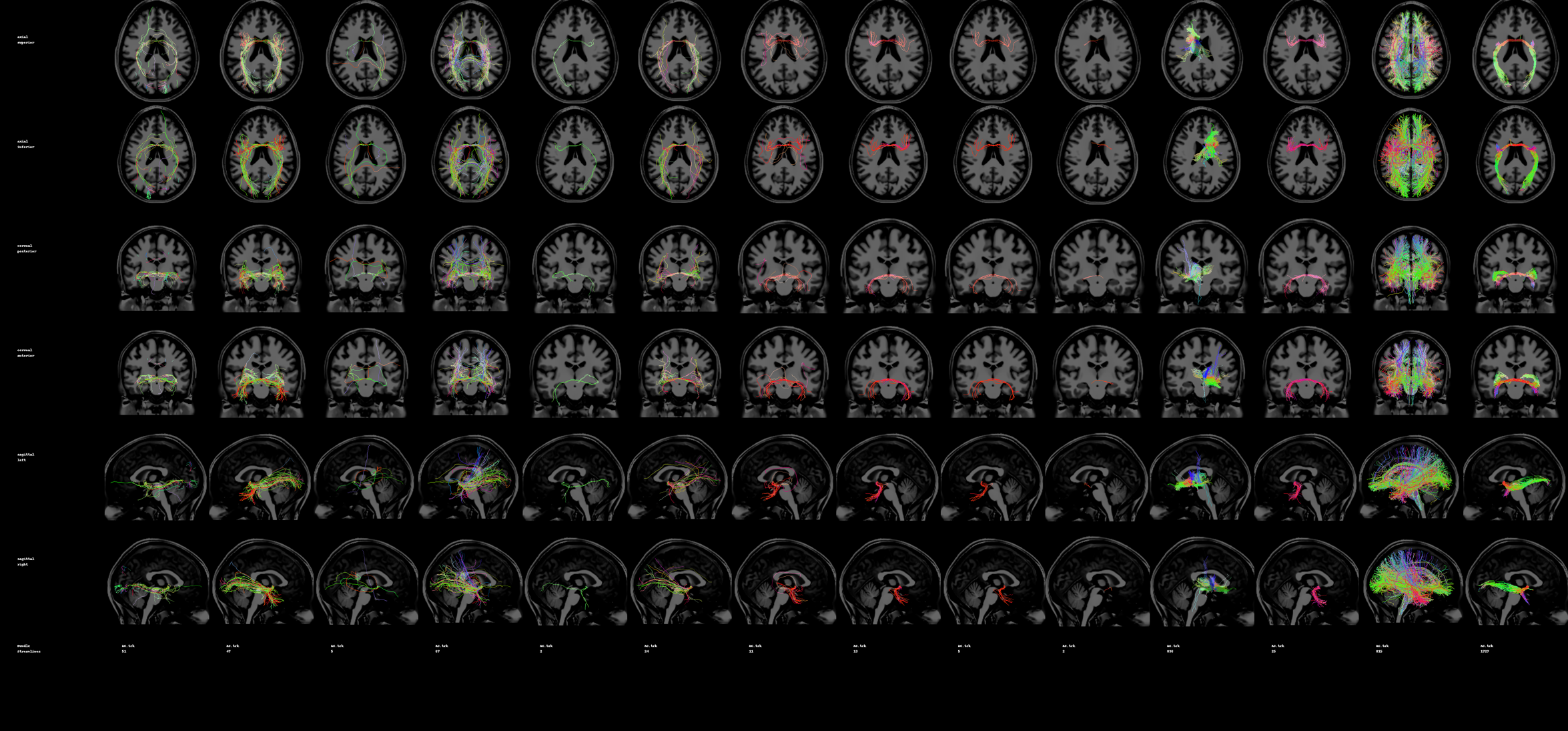

**Supplementary Figure 45.** AC probabilistic submissions for a single subject.
